# Dynamic patterning by the *Drosophila* pair-rule network reconciles long-germ and short-germ segmentation

**DOI:** 10.1101/099671

**Authors:** Erik Clark

## Abstract

*Drosophila* segmentation is a well-established paradigm for developmental pattern formation. However, the later stages of segment patterning, regulated by the “pair-rule” genes, are still not well understood at the systems level. Building on established genetic interactions, I construct a logical model of the *Drosophila* pair-rule system that takes into account the demonstrated stage-specific architecture of the pair-rule gene network. Simulation of this model can accurately recapitulate the observed spatiotemporal expression of the pair-rule genes, but only when the system is provided with dynamic “gap” inputs. This result suggests that dynamic shifts of pair-rule stripes are essential for segment patterning in the trunk, and provides a functional role for observed posterior-to-anterior gap domain shifts that occur during cellularisation. The model also suggests revised patterning mechanisms for the parasegment boundaries, and accounts for the *even-skipped* null mutant phenotype. Strikingly, a slightly modified version of the model is able to pattern segments in either simultaneous or sequential modes, depending only on initial conditions. This suggest that fundamentally similar mechanisms may underlie segmentation in short-germ and long-germ arthropods.

## INTRODUCTION

Like other arthropods, the fruit fly *Drosophila melanogaster* has a segmented body plan. Following a landmark genetic screen [1,2], three decades of genetic experiments have characterised the *Drosophila* “segmentation cascade” in great detail: during the first three hours of development, the anteroposterior (AP) axis of the *Drosophila* embryo is progressively patterned down to cellular-level resolution by an elaborate multi-tiered network of genes and their encoded transcription factors [3,4]. Many fundamental principles of transcriptional regulation have been revealed by studying the organisation and molecular logic of this network [5].

The early stages of *Drosophila* segment patterning have been studied in great detail in recent years, but the later stages, mediated by the so called “pair-rule” genes, have been largely neglected. Recently, Michael Akam and I proposed a revised structure for the pair-rule gene regulatory network, based on inferences made from pair-rule gene expression patterns in wild-type and mutant embryos [6]. Building on that paper, this manuscript is concerned with the reverse exercise, taking the identified genetic interactions as a starting assumption, and using models and simulations to analyse how they collectively lead to complex pattern formation.

This systems-level approach serves two purposes. First, it acts as an important test of the initial conclusions, by ensuring that the inferred network structure is indeed compatible with the experimental observations it is based upon. Second, it improves our understanding of *Drosophila* segmentation by rendering explicit a number of patterning mechanisms that, while implicit in the structure of the genetic network, are far from intuitively obvious.

### An overview of *Drosophila* segment patterning

Current understanding of *Drosophila* segmentation can be summarised as follows. The *Drosophila* embryo is initially polarised by localised maternal factors, such as the homeobox protein, Bicoid, which acts as a graded anterior determinant [7–9]. On the basis of this broad-stroke positional information, transcription factors encoded by the zygotic “gap” genes are expressed in partially overlapping expression domains along the axis. Four in particular are relevant to the patterning of the trunk: Hunchback [10], Krüppel [11], Knirps [12], and Giant [13]. Extensive cross-regulatory interactions between the gap genes lead to the maintenance and refinement of their early patterns as the syncitial blastoderm cellularises (reviewed in Jaeger 2011).

The gap factors are responsible for establishing the first periodic patterns of gene expression within the embryo, via their regulation of the next tier of transcription factors, the pair-rule genes. There are seven canonical pair-rule genes: *hairy* [15], *even-skipped* (*eve,* Macdonald et al. 1986), *runt* [17], *fushi tarazu* (*ftz,* Hafen et al. 1984), *odd-skipped* (*odd,* Coulter et al. 1990), *paired* (*prd,* Kilchherr et al. 1986), and *sloppy-paired* (*slp,* Grossniklaus et al. 1992). Individual stripes of a given pair-rule gene are initially patterned in an *ad hoc* manner through various “stripe-specific” enhancer elements, each of which responds to a given subset of gap inputs [22–25]. These elements act additively, and for certain pair-rule genes they lead to an overall pattern of seven equally-spaced stripes of double segment periodicity along the future trunk of the embryo (reviewed in Schroeder et al. 2011).

Like the gap genes, the pair-rule genes also cross-regulate one another, acting through enhancers known as “zebra elements”, that drive periodic expression [26–29]. For most pair-rule genes, cross-regulatory interactions are required during cellularisation for the completion and regularisation of the double-segment patterns established by the stripe-specific elements [26, 30–34]. For example, certain pair-rule genes lack stripe-specific elements for particular stripes, which are instead patterned purely on the basis of pair-rule inputs acting through the zebra elements. In addition, the stripes of different pair-rule genes need to be precisely-phased relative to one another. Finally, while the five “primary” pair-rule genes (*hairy, eve, runt, ftz,* and *odd*) each receive extensive gap inputs, the two “secondary” pair-rule genes (*prd* and *slp*) do not. They turn on in the trunk considerably later, and are patterned by existing pair-rule stripes through their zebra elements.

The double-segment (“pair-rule”) patterns of pair-rule gene expression established during cellularisation are responsible for establishing the final segment pattern [6,35,36]. Pair-rule factors spatially regulate “segment-polarity” genes, which turn on in narrow stripes of single-segment periodicity at gastrulation. At the same time, and on the basis of the same set of pair-rule inputs, a number of the pair-rule genes transition to segmental expression (i.e. patterns of 7 broad stripes are replaced by patterns of 14 narrow stripes). These late patterns of pair-rule gene expression play functional roles in the “segment-polarity” network, which also includes several components of the Wingless and Hedgehog signalling pathways. The segment-polarity network organises “parasegmental” compartment boundaries and will maintain them throughout development [37–41].

### Knowns, unknowns, and new discoveries

As mentioned above, the earlier, simpler stages of segment patterning are the best understood. The morphogen gradient generated by Bicoid has been the subject of increasingly sophisticated experiments and models (reviewed in Grimm et al. 2010). The gap network has been used as a case study for the application of dynamical systems [43–46] and information theory [47–49] approaches to developmental patterning. Finally, the stripe-specific elements of *eve* are a classic example of modular enhancers and have had their molecular logic dissected in fine detail [50–56] – although it is clear there is still a long way to go until they are fully understood [57].

In contrast, the *Drosophila* pair-rule network has received little systems-level attention this century, despite an initial flurry of developmental genetic studies in the 1980s and 1990s (for example, [34, 58–62]). As a consequence, the later stages of the segmentation process, regulated by the pair-rule genes (i.e. the refinement of the double-segmental pattern during cellularisation and the transition to the single-segmental pattern at gastrulation) have remained relatively mysterious.

The most recent models of pair-rule patterning date from more than ten years ago [63,36]. Since these were published, three important discoveries have been made about segment patterning, all of which challenge established assumptions about the *Drosophila* segmentation cascade, and all of which concern the pair-rule genes in some way. So long as the pair-rule network remains poorly-understood, key questions raised by these findings will go unanswered.

The first discovery is from comparative studies in other arthropod embryos. *Drosophila* is a “long-germ” embryo, patterning almost all of its segments simultaneously in the trunk prior to germband extension. However, the ancestral mode of arthropod development is “short-germ” embryogenesis, in which segmentation is sequential, and coordinated with germband extension [64]. Orthologs of the pair-rule genes play a key role in segment patterning in all arthropods yet studied (for example, [65–68]), but, in short germ embryos, their expression has been shown to oscillate in a posterior “segment addition zone” throughout germband extension [69,70]. This periodic dynamic expression indicates that in these organisms they are either components of, or entrained by, a segmentation “clock” [71]. How the expression of the pair-rule genes in long-germ embryos such as *Drosophila* relates to their expression in short-germ embryos (for example the model beetle, *Tribolium castaneum)* is unclear. It is thus not understood how long-germ segmentation was derived from short-germ segmentation, an important evolutionary transition that has occurred multiple times independently within the higher insects [72].

Second, quantitative studies have revealed that the domains of gap gene expression in the trunk of the embryo shift anteriorly across the cellularising blastoderm over the course of nuclear division cycle 14 [73–75]. These shifts have been used to argue that the *Drosophila* segmentation cascade does not simply transduce positional information present in the initial maternal signals, but rather that internal network dynamics play a crucial role in the specification of final cell states [73,76,77]. However, while the mechanistic origins of the shifts have been probed extensively [43,44,78,46], their functional role (if any) remains unclear. The main job of the *Drosophila* gap system is to spatially regulate the pair-rule genes, and indeed the shifting domains of gap expression are reflected by shifting stripes of pair-rule gene expression [74,79]. Given that the posterior gap domains are a relatively recent invention within the dipteran lineage (reviewed in Jaeger 2011), determining the downstream effects of the shifts on the pair-rule system might therefore shed light on the evolution and function of the gap network.

The final key finding relates to the structure of the pair-rule network itself. In our recent paper on the pair-rule network [6], Michael and I showed that many regulatory interactions between pair-rule genes are temporally regulated, requiring either the presence or absence of a broadly expressed transcription factor, Odd-paired [80], which mediates the transition from double-segment to single-segment periodicity at gastrulation. We argued that the pair-rule network is best viewed as two distinct networks, one operating during cellularisation and one during gastrulation, each with a specific topology and resultant dynamics. Analysing the “early” (cellularisation-stage) and “late” (gastrulation-stage) pair-rule networks separately should lead to a better understanding of pair-rule patterning, and might also shed light on why the network shows this bipartite organisation in the first place.

In this manuscript, I build on the regulatory analysis presented in the Odd-paired paper to firmly establish the topology of the “early” pair-rule gene network (responsible for regulating the double-segmental stripes during cellularisation). Through dynamical analysis of this network, I demonstrate a role for the anterior shifts of gap gene expression in correctly phasing the pair-rule stripes. I then model and simulate the pair-rule system as a whole, revealing important downstream effects of these early shifts on the final segment pattern. I also find that graded Eve stripes are not strictly necessary for pair-rule patterning, and I explain the aetiology of the surprisingly severe *eve* null mutant phenotype. Finally, I show that a slightly modified version of the *Drosophila* pair-rule network gains the capacity to pattern in both simultaneous and sequential modes, conceptually reconciling long and short germ segmentation.

## RESULTS

### 1. Topology of the early pair-rule network

During cellularisation, pair-rule gene expression patterns are influenced by both gap and pair-rule factors. For the most part, these effects consist of gap factors acting directly on stripe-specific elements, and pair-rule factors acting directly on zebra elements. While the regulatory regions of all seven pair-rule genes have been carefully characterised, and all such enhancer elements are thought to have been identified [26], the relative roles of the two classes of elements are not entirely clear. Originally, gap factors were thought to govern the expression of *hairy, eve,* and *runt,* while pair-rule factors governed the expression of *ftz, odd, prd,* and *slp* [30,33,58]. More recently, it has been proposed that stripe-specific elements play the major role in patterning the expression of all five primary pair-rule genes, with zebra elements responsible only for minor stripe refinements and secondary pair-rule gene expression [26,81].

Resolving this issue involves carefully evaluating the contribution of cross-regulation to pair-rule patterning during cellularisation. To do this, it is important both to determine which particular cross-regulatory interactions are active during this stage, and to clarify the extent of their effects on expression. Early and late regulation of the pair-rule genes was analysed in Clark & Akam 2016 [6]. Extracting the relevant regulatory interactions from that work yields the network structures diagrammed in Figure 1.

**Figure 1:**
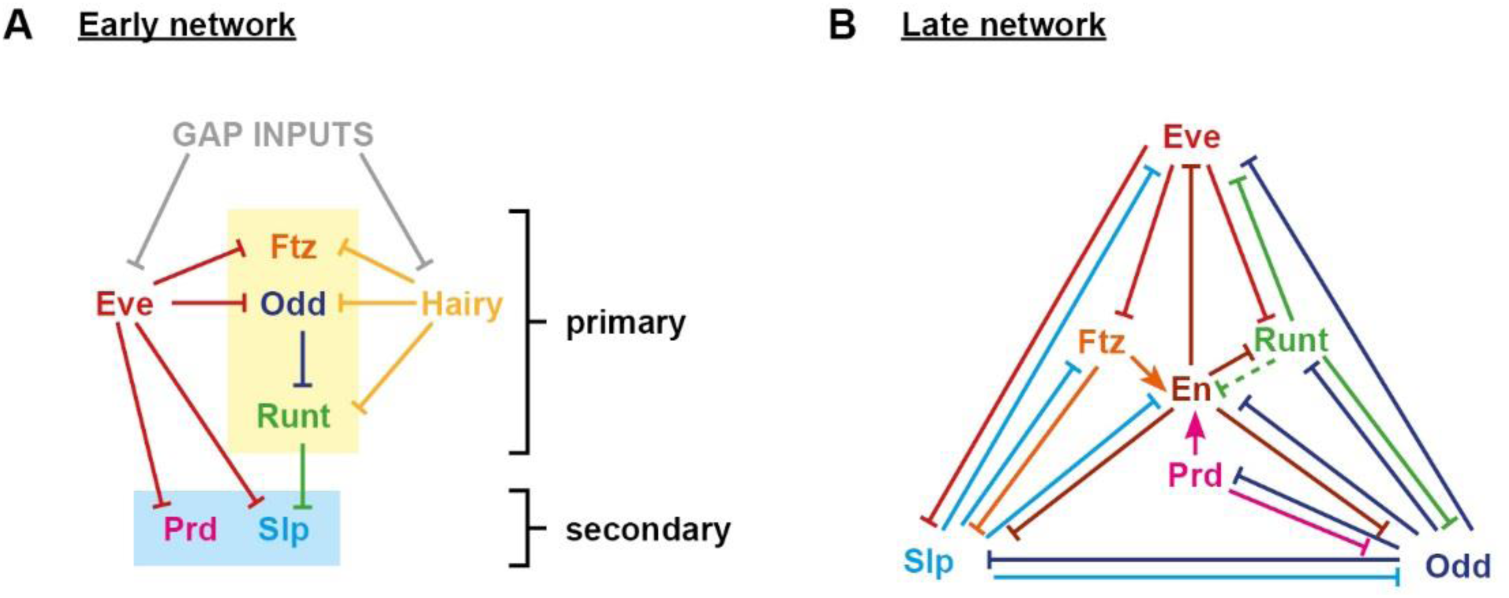
Topologies of the early and late pair-rule networks. **A**: Cross-regulatory interactions between pair-rule genes during cellularisation (note that gap inputs into *ftz, odd,* and *runt* are not depicted). The zebra elements *of ftz, odd,* and *runt* (yellow box in **A**) turn on earlier than those of *prd* and *slp* (blue box in **A**). **B**: Cross-regulatory interactions between pair-rule genes during gastrulation. Hammerhead arrows represent repression; pointed arrows represent activation. The dotted repression of En by Runt in **B** represents the finding that while the odd-numbered *en* stripes are sensitive to Runt, the even-numbered stripes are regulated differently [139]. Note that the regulation of Runt in the late network reflects the regulatory logic of the “seven-stripe” element, rather than that of the “six-stripe” element [82]. The *runt* seven-stripe element drives strong expression in the primary stripes, and eventually drives expression in the secondary stripes as well. The *runt* six-stripe element drives strong expression in the secondary stripes, which initially overlap with Odd expression (see Clark & Akam 2016 for details).

Unsurprisingly, the late network (Figure 1B) is a dense web composed largely of mutually repressive pairs of interactions, consistent with its function of specifying and maintaining discrete segment-polarity states. In contrast, the early network (Figure 1A) is relatively sparse, including only six regulatory interactions between the five primary pair-rule genes, and only nine regulatory interactions when all seven pair-rule genes are included. For a more in-depth justification of the early network topology, see Supplement 1, in which I analyse cellularisation stage patterns of pair-rule gene expression in pair-rule mutant embryos, and discuss this evidence in the context of the wider experimental literature relating to pair-rule gene regulation.

As described previously [6], almost all of the cross-regulatory interactions depicted in the early and late pair-rule networks were identified many years ago by way of genetic experiments (see also Appendix 1). However, as yet there has been no consensus structure for the pair-rule system as a whole (for various different summary topologies see [26,36,66]). Given that accurate knowledge of the structure of a gene regulatory network is a prerequisite for correctly predicting and analysing the expression dynamics it produces, the well-substantiated network diagrams presented here provide a foundation for new insights into segment patterning.

In the first instance, the topology of the early network (Figure 1A) permits a number of broad conclusions to be drawn about *Drosophila* blastoderm patterning:

1) *Two classes of primary pair-rule genes* Two of the primary pair-rule rule genes, *hairy* and *eve*, are “input-only” factors in the network, as their expression does not seem to be directly and systematically affected during cellularisation by the expression of any pair-rule factor. Their regular seven stripe patterns are thus presumably directly specified by regulation from the gap genes. This phenomenological inference corresponds well with what is known of the regulatory landscape of each gene: both *hairy* and *eve* possess a full complement of stripe-specific elements, and both lack a canonical zebra element (Schroeder et al. 2011 and references therein). Note that *eve* does however possess a periodically expressed “late” element [23,24] that turns on towards the end of cellularisation, corresponding to the time at which significantly aberrant *eve* expression patterns are first observed in pair-rule mutant embryos [32]. In contrast, the genetic evidence suggests that the expression patterns of the remaining primary pair-rule genes - *runt, ftz,* and *odd* - are specified largely by inputs from other pair-rule factors, consistent with findings that each of the three possesses a zebra element expressed throughout cellularisation [26,27,82].
2) *Patterning by repression* The interactions between the pair-rule genes are generally repressive, consistent with established regulatory models of ubiquitous activation (by, for example, maternally provided factors) combined with precisely positioned repression from other segmentation genes [60,83–85]. While certain of the pair-rule factors have been shown to quantitatively upregulate the expression of other pair-rule genes (for example Ftz is thought to directly activate *odd, prd,* and itself [86–88]), for the most part these effects do not seem to be important for qualitatively determining the spatial pattern of gene expression (but see point 3 below). Notably, the primary pair-rule genes *runt, ftz,* and *odd* are each repressed by two other pair-rule factors, with the result that anterior borders of each set of stripes are positioned by a different repressor to that which patterns the posterior borders. Specifically, *ftz* and *odd* are both repressed by Eve from the anterior and by Hairy from the posterior, while *runt* is repressed by Hairy from the anterior and by Odd from the posterior (see Figure 2). These asymmetrical regulatory inputs are not straightforward to reconcile with the fairly symmetrical expression patterns of these genes: *ftz* and *odd* expression is largely complementary with *eve* (Figure 2F,G), and *runt* expression is largely complementary with *hairy* (Figure 2E). I provide an explanation for this discrepancy below.
3) *Dominance of pair-rule repression over gap inputs* While *runt*, *ftz*, and *odd* all possess at least some stripe-specific elements in addition to their zebra elements [26], it is clear that regulatory inputs received from the pair-rule factors largely override those provided by the gap factors (at least during the latter half of cellularisation). For example, ectopic expression of Eve or Hairy protein has been shown to result in the loss of all *odd* and *ftz* stripes [59,89,90], indicating that repression mediated by pair-rule factors also shuts off expression from the stripe-specific elements. In contrast, *ftz* and *odd* expression expands in *hairy* or *eve* mutant embryos [30,31] – including into regions outside the expression domains of the stripe-specific elements – indicating that repression from gap factors does not influence expression governed by zebra elements. This phenomenological evidence is suggestive of zebra element “enhancer dominance”, and is consistent with studies showing that pair-rule factors tend to act as long-range repressors along a stretch of DNA, while gap factors act only at short-range [91–93]. The inference is that, while the stripe-specific elements of *runt, ftz,* and *odd* likely play a role in the early establishment of a periodic pattern, the precise boundaries of their stripe domains soon come to be specified largely by their zebra elements. However, note that an apparent exception is presented by the anterior boundaries of the *ftz* stripes. *ftz* possesses stripe-specific elements for all stripes except stripe 4 [26]. During cellularisation, the anterior boundary of *ftz* stripe 4 aligns closely with the anterior boundary of *odd* stripe 4 (reflecting their common regulation by Eve), but the anterior boundaries of the remaining *ftz* stripes are anteriorly shifted relative to the anterior boundaries of the *odd* stripes, by roughly one cell row [6]. It seems likely that this discrepancy reflects specific positioning of the *ftz* anterior boundaries by gap inputs (something that may become “locked in” by direct Ftz autoregulation through the zebra element). For the purposes of this manuscript I ignore this effect: while it may well contribute to the robustness of pair-rule patterning, it cannot be absolutely required for the process since a normal parasegment boundary is still produced by stripe 4, which lacks gap inputs.
4) *Hierarchical flow of positional information* The topology of the early network (Figure 1A) implies a hierarchical flow of positional information within the pair-rule system during cellularisation. First, the gap factors pattern the expression of Hairy and Eve. Second, the resulting periodic inputs from Hairy and Eve organise (directly or indirectly) the expression of the other three primary pair-rule genes. Finally, Eve and Runt together pattern the early expression of the two secondary pair-rule genes, *prd* and *slp.* Significantly, all regulatory interactions in the early network are unidirectional – there are no mutually repressive, “switch-like” interactions like those in the late network – suggesting that inputs from Hairy and Eve unilaterally impose patterns onto their targets (both direct and indirect). The primary pair-rule genes *odd* and *runt* play intermediary roles in this process: Odd, patterned by Hairy and Eve, regulates the expression of Runt, while Runt, patterned by Hairy and Odd, regulates the expression of *slp.* In contrast, Ftz does not significantly affect the spatial limits of expression of any pair-rule gene (other than itself) during cellularisation and is thus an “output-only” factor in the network.

**Figure 2:**
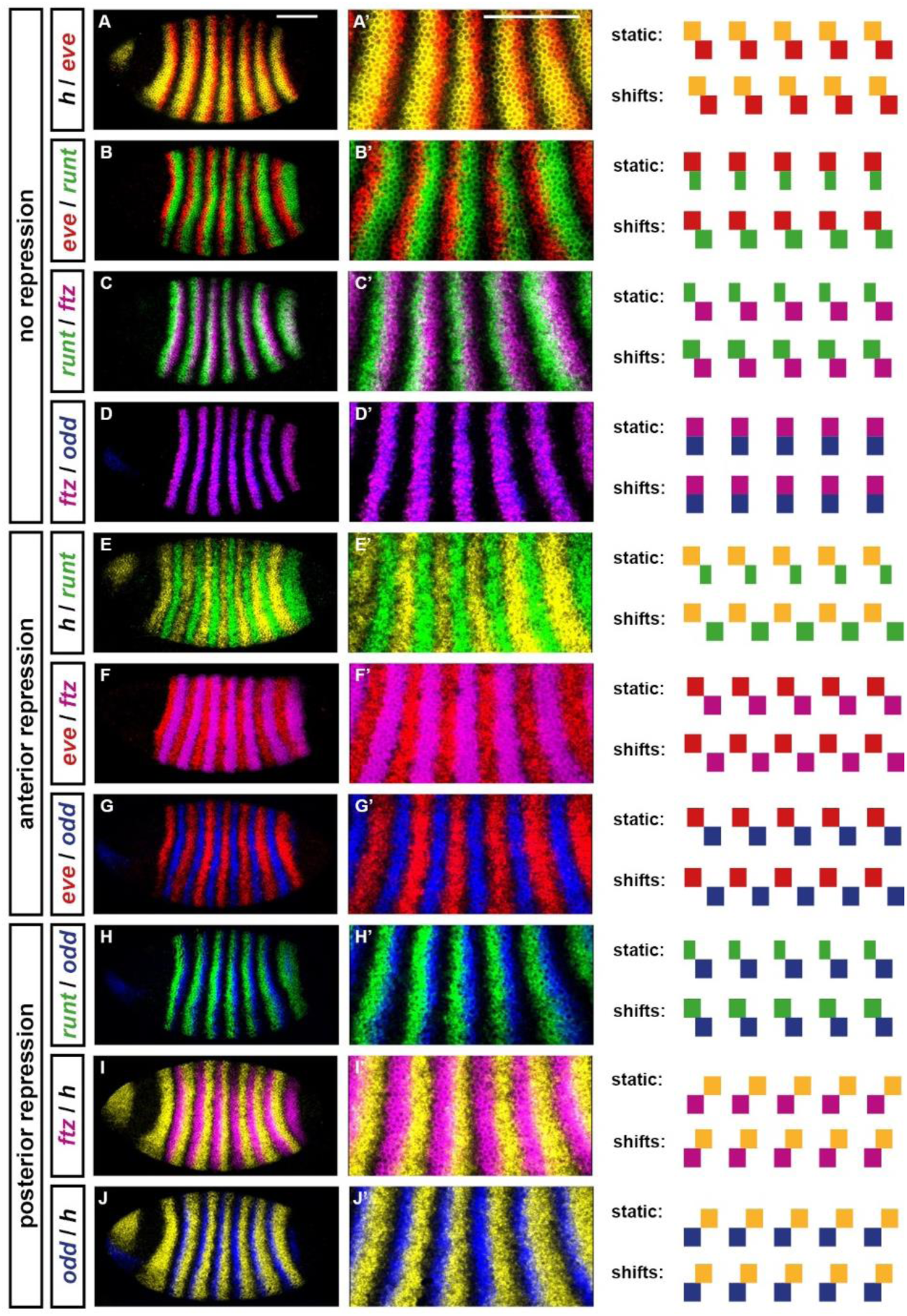
stripe phasings of the primary pair-rule genes during cellularisation. Double fluorescent *in situs* for all pairwise comparisons of the five primary pair-rule genes: *hairy, eve, runt, ftz,* and *odd.* All embryos are at approximately mid cellularisation. Fluorescence channels are false coloured, with each gene’s transcripts assigned a different colour (yellow=*hairy*; red=*e*ve; green=*runt*; magenta=*ftz*; blue=odd). **A-J**: lateral views of whole embryos; anterior left, dorsal top. **A’-J’**: enlarged views of stripes 2-6. In panels **A-D** the stripes do not regulate each other; in panels **E-G** one set of stripes patterns the anterior borders of the other (Hairy represses *runt;* Eve represses *ftz/odd);* in panels **H-J** one set of stripes patterns the posterior borders of the other (Odd represses *runt;* Hairy represses *ftz/odd).* To the right of the *in situ* images are schematics showing predicted stripe phasings from simulating the early pair-rule network (Figure 1A) with either static or shifting gap inputs (see Figure 4). Shifting inputs predict the observed patterns more accurately: for example, note the width and relative phasings of the *runt* stripes (**B,C,E,H**), or the fairly complementary expression patterns in cases of anterior repression **(E-G)**. Scale bars 100 μm.

### 2. Potential sources of spatial resolution

The final segment pattern is precise and elaborate, consisting of at least six distinct domains of segment-polarity gene expression within each double-segment repeat. At the minimum, this template consists of a repeating tripartite pattern of mutually exclusive En, Odd, and Slp domains, which collectively determine the position and polarity of parasegment boundaries (Figure 3A). This threefold division of segments conforms to prescient theoretical predictions made by Hans Meinhardt in the early 1980s [94–96].

**Figure 3:**
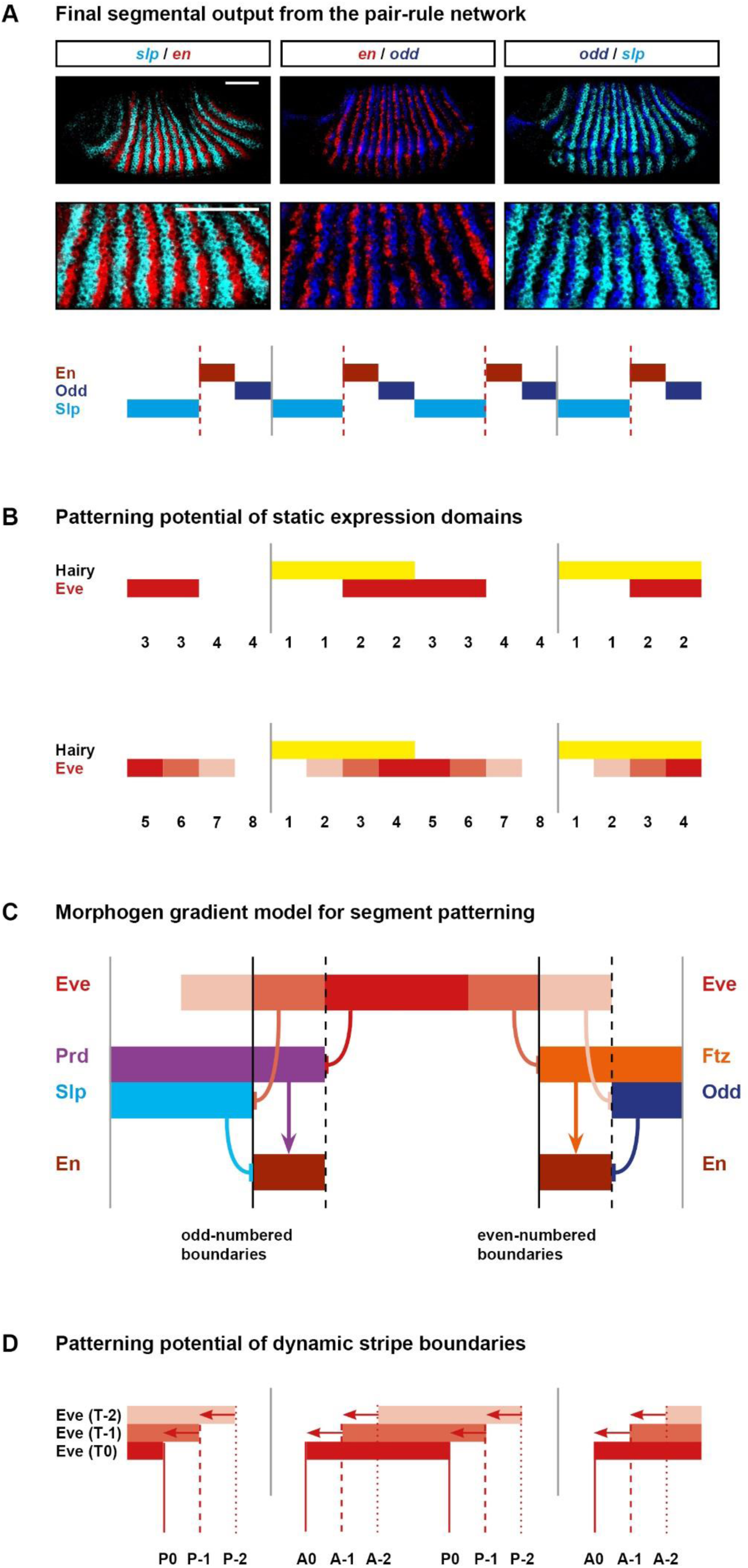
Static *versus* dynamic models for pair-rule patterning. **A:** The template for polarised parasegment boundaries is formed by a repeating pattern of En, Odd, and Slp stripes. Top: whole mount double fluorescent *in situs* (anterior left, dorsal top) showing that *en, odd,* and *slp* are expressed in abutting, mutually exclusive domains. Middle: blown up views of the stripes. The *en* and *odd* stripes are about one cell wide, while the *slp* stripes are about two cells wide. Bottom: schematic of the overall pattern (anterior left). The grey vertical lines indicate the span of an initial pair-rule repeat relative to the final output pattern. Parasegment boundaries (dotted red lines) will form at the interface between En and Slp domains. The Odd stripes act as buffers between En and Slp, precluding ectopic boundary formation. For details, see Jaynes & Fujioka 2004; Mullen & DiNardo 1995; Cadigan et al. 1994; Meinhardt 1984. Scale bars 100μm. **B:** Schematics indicating the number of distinct states that can be specified by static domains of Hairy and Eve expression. Top: Hairy and Eve are both Boolean variables. There are only four possible expression states (1: Hairy on, Eve off; 2: Hairy on, Eve on; 3: Hairy off, Eve on; 4: Hairy off, Eve off). This is not sufficient to pattern a double-segment repeat (typically 7-8 cells across) to single-cell resolution. Bottom: Hairy is still Boolean, but Eve is now a multi-level variable. Different shades of red represent different levels of Eve activity: low (lightest), medium, or high (darkest). There are now 8 different possible combined expression states of Hairy and Eve, sufficient to uniquely specify position within a double-segment repeat. **C:** Summary of the morphogen gradient model for the patterning of the segmental *engrailed* stripes [61,63,100]. *prd, slp, ftz,* and *odd* are all repressed by specific concentration thresholds of Eve protein. Graded Eve stripes therefore result in distinct posterior boundaries of Prd and Slp, and distinct anterior boundaries of Ftz and Odd. En is activated anywhere where Prd or Ftz are expressed but Slp and Odd are not, and therefore turns on in narrow stripes at the margins of the Eve stripes. Parasegment boundaries (solid black lines) will form to the anterior of the En domains. **D:** Kinematic expression shifts can also increase the positional information conveyed by a given stripe domain, if expression boundaries of the downstream genes are defined at different times. Boolean Eve stripes are depicted travelling from posterior to anterior over time (darker red represents a more recent position). The anterior and posterior boundaries of the stripes (vertical lines) are located in different positions at each of the three time points.

Most of the segmental stripes of pair-rule and segment-polarity gene expression are just a single cell wide. These “late” patterns are specified cell-autonomously over the course of gastrulation, on the basis of inputs from the “early” pair-rule patterns set up during cellularisation (intercellular signalling is not functional until germband extension, DiNardo et al. 1988). Therefore, the early patterns must necessarily contain sufficient spatial information to pattern the anteroposterior axis down to cellular level resolution. This can be accomplished by precisely phasing the individual pair-rule stripes within each double segment repeat, so that at gastrulation each nucleus starts off with a unique combination of pair-rule factors, permitting it to follow an expression trajectory different from that of its immediate neighbours [35].

Appropriate relative phasing of the pair-rule stripes could theoretically originate from independent patterning of each gene by its own specific set of gap inputs. However, the topology of the early pair-rule network presented in the previous section implies that (almost) all of the spatial information in the final segment pattern must eventually trace back to the stripe-specific elements of *hairy* and *eve*. How is it possible that just two independent spatial signals are able to give rise to such a precise and high-resolution final output?

#### The current hypothesis: morphogen gradients

Since the early 1990s, the answer has been thought to be that quantitative information inherent in the initial pair-rule signals is instructive for generating the final segment pattern [59,61,98–100]. In particular, attention has focused on the graded margins of the early Eve stripes. These have been thought to act as local morphogen gradients, repressing different target genes at different concentration thresholds, and thus differentially positioning their respective expression boundaries. Four functionally distinct levels of Eve activity (HIGH, MEDIUM, LOW, and OFF) would be sufficient to provide single-cell resolution within a double segment repeat (around 7-8 cells wide on average). Combining this information with a second, Boolean (i.e. ON/OFF) signal – provided by Hairy – that differentiated the anterior half of these symmetrical gradients from the posterior half, the identity of each cell in the repeat could be uniquely specified (Figure 3B). These assumptions form the basis of the most recent dynamical model of pair-rule patterning [63].

Specifically, “French flag”-style patterning of target genes by Eve is thought to be responsible for the patterning of both sets of *en* stripes, with odd- and even-numbered boundaries being specified by the anterior and posterior margins of the Eve stripes, respectively (Fujioka et al. 1995; Figure 3C). The odd-numbered *en* stripes are activated by Prd and repressed by Slp, both of which are repressed by Eve. Eve is thought to repress *slp* at a lower concentration threshold than required to repress *prd,* therefore differentially patterning the posterior boundaries of the *prd* and *slp* expression domains, and consequently permitting *en* expression only in a single cell row at the anterior edge of the Eve stripes, where Prd is present but Slp is not.

Along similar lines, the even-numbered *en* stripes are activated by Ftz but repressed by Odd, both of which are (like Prd and Slp) repressed by Eve. *odd* is thought to have a higher sensitivity to repression by Eve than does *ftz,* thus resulting in narrow stripes of Ftz-positive, Odd-negative cells at the posterior borders of the Eve stripes, in which *en* is activated.

While this patterning model seems to follow in a straightforward manner from the genetic evidence – the relevant expression boundaries of *prd, slp, ftz,* and *odd* are all mispatterned in *eve* mutant embryos (see below, and Supplement 1) – there are several potential problems with it. First, Eve shows no evidence of differentially patterning *ftz* and *odd* during cellularisation, with the *ftz* / *odd* offsets instead seemingly stabilised after gastrulation by the combined activity of Opa and Runt [6]. Second, in order for the model to work, not only would the cell-autonomous transcriptional response of Eve targets to differences in Eve concentration have to be extremely reliable, but the Eve stripes themselves would have to have extremely precise intensity profiles in order to provide the necessary spatial information to these targets. Neither of these conditions are likely to be met in reality: transcription in the blastoderm is intrinsically highly stochastic [101,102], while Eve shows significantly different expression levels between different stripes throughout most of cellularisation [74].

#### An alternative hypothesis: temporal dynamics

How, then, might the spatial resolution of the pair-rule pattern be explained? Classic models of *Drosophila* segment patterning assume that segmentation genes are expressed in static domains [35,58,96,103]. However, as discussed in the Introduction, quantitative studies have revealed that in the posterior half of the embryo (which encompasses pair-rule stripes 3-7), the gap gene expression domains shift gradually towards the anterior over the course of cellularisation. Systems-level analyses have shown that these shifts arise due to feedback interactions within the gap gene network: more posteriorly-expressed factors asymmetrically repress more anteriorly-expressed factors and therefore progressively displace them (reviewed in Jaeger 2011).

In the previous section, I discussed how *hairy* and *eve* are not affected by pair-rule gene expression, and are therefore presumably regulated by gap inputs throughout the entire duration of cellularisation. Because the stripes of Hairy and Eve are responsible for organising the expression of the remaining pair-rule genes, their sustained link to the gap system transfers gap domain expression dynamics to the pair-rule system, resulting in subtle but observable posterior-to-anterior shifts during cellularisation [74].

Because pair-rule stripe boundaries are consequently located in different places at different timepoints during cellularisation, these shifts present a source of additional spatial information that might be utilised for blastoderm patterning (Figure 3D). In the following section, I use simulations to explore the ways in which these shifts are likely to influence pair-rule gene expression, and propose that they play a fundamental role in segmentation. (Details of all simulations are given in Supplement 2.)

### 3. Expression shifts are required for appropriate phase 2 expression patterns

#### The early pair-rule network requires dynamic inputs to produce observed stripe phasings

During cellularisation, the primary pair-rule genes are expressed in precisely phased, partially overlapping stripes of double-segment periodicity (Figure 2). From anterior to posterior within a pattern repeat, stripes are arranged in the order *hairy; eve; runt; ftz/odd; hairy,* with substantial overlaps between neighbouring pairs within the sequence (i.e. between *hairy* and eve, between *eve* and *runt,* between *runt* and *ftz/odd,* and between *ftz/odd* and *hairy*).

We know that this pattern is established by the early pair-rule network, given wild-type expression of *hairy* and *eve*. For reasons outlined above, it is unlikely that the stripes of Hairy and Eve provide extensive quantitative information to their target genes. Given these three assumptions: 1) the network topology (as in Figure 1A); 2) partially overlapping stripes of *hairy* and *eve* (as in Figure 2A); and 3) approximately Boolean regulatory activity, is it possible to account for observed pair-rule stripe phasings?

Assuming static gap inputs and hence static expression of Hairy and Eve, it is not possible to recover the observed pattern (Figure 4A,B; Movie 1). The early network has only four possible stable states (Hairy; Hairy + Eve; Eve + Runt; Ftz + Odd) and therefore when provided with partially overlapping domains of Hairy and Eve, it yields a pattern where the *runt* stripes are narrower than the others, and half of the observed expression overlaps are missing. Overlaps between *hairy* and *eve* and between *eve* and *runt* are present, but overlaps between *runt* and *ftz/odd* and between *ftz/odd* and *hairy* are not. This is because Odd represses *runt* and Hairy represses *ftz* and *odd,* precluding any stable coexpression of these particular pairs of genes.

**Figure 4:**
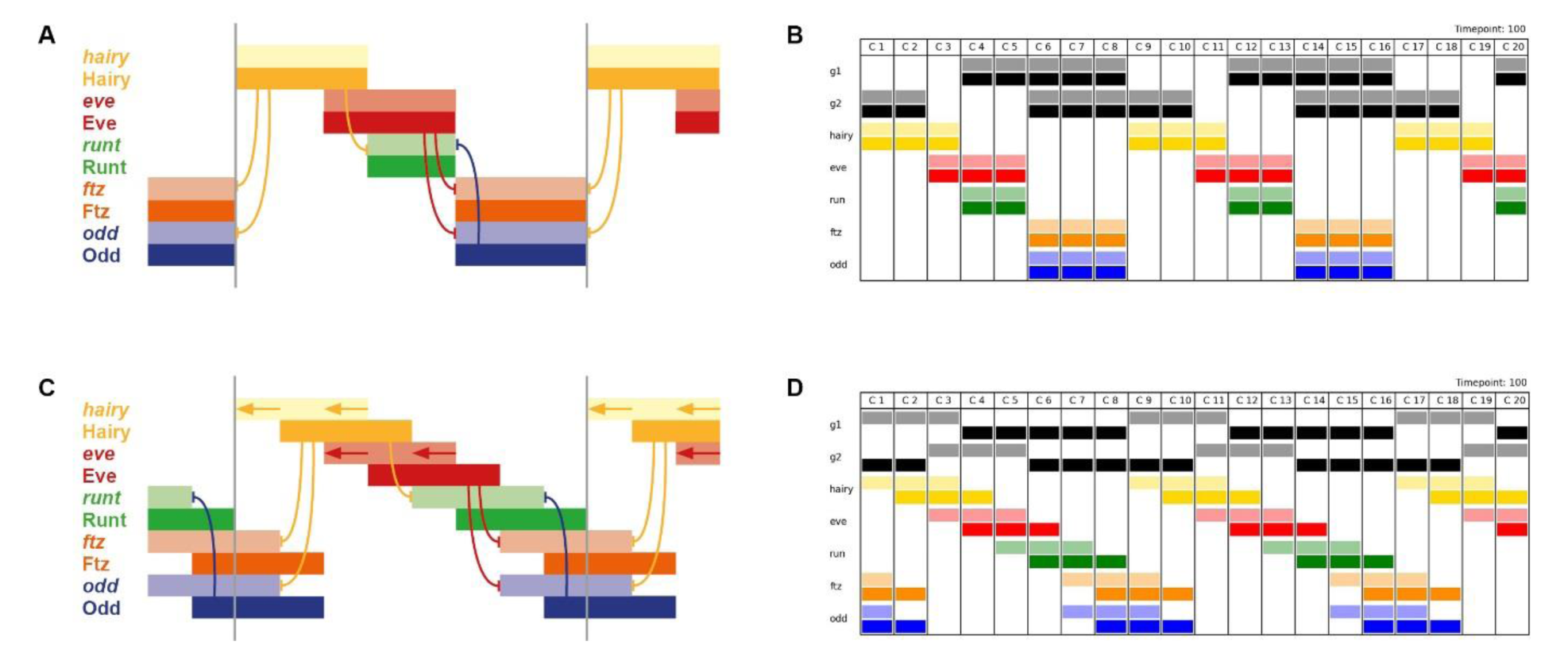
Dynamic gap inputs are required for regularly phased stripes of primary pair-rule gene expression. **A,B:** Expression patterns generated by simulation of the early pair-rule network, assuming static stripes of Hairy and Eve. *ftz* and *odd* do not overlap with *runt* or *hairy,* and transcript (pale colour) and protein (dark colour) domains coincide exactly. **C,D:** Expression patterns generated by simulation of the early pair-rule network, assuming Hairy and Eve stripes shift from posterior to anterior over time. *ftz* and *odd* overlap with both *runt* and *hairy*, and protein activity lags behind transcript expression. **A,C:** Regulatory schematics showing how Hairy and Eve together organise the expression of the other primary pair-rule genes, *runt*, *ftz*, and *odd*. Hammerhead arrows represent repression, grey vertical lines indicate the span of a double-segment repeat. **B,D:** Simulation output showing the same expression patterns as **A** and **C**, respectively (see Movies 1 and 2). The columns C1 to C20 represent different cells along an idealised AP axis. Pale colours represent transcription of a given gene, dark colours represent protein activity. Generalised “gap inputs” G1 and G2 repress *hairy* and *eve* expression, respectively. In **B**, G1 and G2 expression domains are static, due to autoactivation. In **D**, G1 and G2 expression is dynamic, due to autorepression. In all panels, anterior is to the left.

However, as we now know, the stripes of Hairy and Eve are not static, but instead shift anteriorly over time, driven by dynamic gap gene expression. Strikingly, when shifting gap inputs are included in the starting assumptions, it becomes possible for wild-type stripe phasings to be produced by the network (Figure 4C,D; Movie 2). If the rate of expression shifts is on the same timescale as the time delays involved in protein synthesis and protein decay, discrepancies between domains of transcript expression and domains of protein expression are predicted to occur. This phenomenon has been explicitly demonstrated for gap domains [73]. It is therefore likely to also hold true for the pair-rule genes, which have similarly compact transcription units (FlyBase) to the gap genes, and produce protein products which turn over extremely rapidly [83,86]. As shown in Figure 4 and Movie 2, the existence of such transcript/protein offsets would generate the transcript/transcript and protein/protein overlaps between repressors and their targets that are missing in the static models.

The additional overlaps produced by the dynamic model arise because domains of active transcription do not necessarily reflect domains of protein activity, and *vice versa* (Figure 4C,D). Overlaps between *runt* and *odd* at the anteriors of the *odd* domains reflect the absence of Odd protein activity, while overlaps between Runt and Odd at the posteriors of the Runt domains occur in the absence of *runt* transcription (and similarly for *odd/hairy* and Odd/Hairy).

This phenomenon also resolves the apparent conflict between gene regulation and gene expression noted above for *runt, ftz,* and *odd.* The offsets between domains of active transcription and domains of protein activity will act to space out the stripes of a target and a repressor when the repression comes from the anterior, but will cause overlaps when the repression comes from the posterior (compare model predictions for static *versus* shifting inputs in Figure 2E-G and Figure 2H-J). Therefore, cases of anterior repression (i.e. Hairy/*runt*, Eve/*odd*, and Eve/*ftz*) are predicted to produce fairly complementary expression patterns, giving the false impression of symmetrical regulation, and masking the role of the posterior repressors.

#### Pair-rule gene expression is locally unstable but results in a persistent global pattern

As illustrated by the simulation output for the dynamic model (Movie 2), the overlaps produced by the transcript/protein offsets are of course unstable, and are thus transient, resolving over a period of some minutes within each nucleus. However, because the inputs from Hairy and Eve keep changing within any given cell, the system does not stabilise and a certain fraction of nuclei within the tissue will be in these transient states at any given moment. So long as the gap gene shifts continue (and while the early network persists), the overall pattern will be maintained, with all the stripes travelling across the tissue in concert like an elaborate Mexican wave.

The appropriate shift speed to recapitulate the *Drosophila* pair-rule pattern is that covering the distance of one nucleus in roughly the same time as it takes to synthesise new protein or decay existing protein (Figure 4–supplementary figure 1; Movies 1-5). If the shift speed is too fast (Figure 4–supplementary figure 1D,E; Movies 4,5), the relative phasing of the various pair-rule stripes changes significantly. If the shifts are too slow (Figure 4–supplementary figure 1B; Movie 3), the pattern just looks similar to the static model.

#### How much shifting is theoretically required?

It seems clear that some degree of posterior-to-anterior shifting occurs in wild-type embryos for all pair-rule stripes posterior to stripe 2. However, due to the limited duration of cycle 14, these shifts are of fairly small magnitude (see Discussion). Is it plausible to suggest that they play a functional role in patterning the blastoderm? To address this question, I used the dynamic model to determine the minimum amount of shifting required to produce an appropriate pattern of primary pair-rule gene expression (see Supplement 2 for details).

If *hairy* and *eve* are the only genes that receive gap inputs, as in the original simulation (Movie 2), a relatively long time is required for the wild-type pattern to first establish. This initial time delay corresponds to 3x the synthesis/decay time delay and therefore a minimum shift magnitude of three nuclei. However, by mimicking the effect of the stripe-specific elements and including independent gap patterning of *runt* and/or *ftz/odd* during the early stages of the simulation, the time required to produce the final pattern is considerably reduced.

Specifically, if either *runt* (Movie 6) or *ftz/odd* (Movie 7) are controlled by gap inputs until the pattern is established, the time delay is only 2x the synthesis/decay time, equating to a shift of two nuclei. Moreover, if both *runt* and *ftz/odd* are controlled by gap inputs while the pattern is established (Movie 8), it is possible for an appropriate expression pattern to emerge immediately, corresponding to the shortest possible establishment time (1x the synthesis delay) and a shift of only one nucleus.

Thus, the stripe-specific elements of *runt, ftz,* and *odd* may serve to increase patterning speed within the *Drosophila* blastoderm and therefore slightly reduce the total time required for embryonic development (see Discussion). They may also be involved in compensating for inappropriate early expression from Hairy and Eve. For example, the “interstripe” (gap) between *hairy* stripes 3 and 4 resolves relatively late, correlating with an early absence of *runt/ftz/odd* zebra element reporter expression throughout the corresponding region of the embryo [26]. However, endogenous *runt/ftz/odd* stripe 3 expression is nevertheless established in a timely fashion, as all three genes possess a stripe-specific element for this particular stripe.

### 4. Dynamic early expression is required for appropriate late expression patterns

#### A logical model of the pair-rule system

In summary, dynamic gap inputs seem to be involved in properly patterning early primary pair-rule gene expression. But what are the consequences for downstream segment patterning? In order to explore this question, I built a simple “toy model” of the entire pair-rule system and simulated its behaviour in the presence and absence of posterior-to-anterior shifts.

A detailed description of this model is provided in Supplement 2. Briefly, the model simulates the known regulatory interactions between nine transcription factors: all seven pair-rule factors, the segment-polarity factor En, and the temporal factor Opa. Additional temporal information is provided by a hypothetical signal “X”, which controls the onset of secondary pair-rule gene expression. Finally, as in the simulations of primary pair-rule gene expression (Figure 4; Movies 1-8), two sets of repressive “gap” inputs (“G1” and “G2”) regulate *hairy* and *eve* respectively. Note that “X”, “G1”, and “G2” are not necessarily meant to represent specific transcription factors *per se,* but rather are the simplest way to impart the extrinsic spatiotemporal regulatory information known to influence the pair-rule system.

Each of the 12 total components in the model takes the form of a Boolean variable, which may be on or off in any given “nucleus” along an abstracted region of “AP axis” (a one-dimensional array of 20 autonomous cells). The system is iteratively simulated by simultaneous discrete time updates, with simple time delay rules governing the synthesis of “protein” from “transcript”, and the degradation of protein after the cessation of active transcription (see Supplement 2). The system is fully cell-autonomous, with the state of each nucleus evolving independently of the states of its neighbours. This last assumption is realistic for the pair-rule system: because pair-rule transcripts are apically localised, pair-rule cross-regulation occurs in an effectively cell-autonomous environment even before the blastoderm fully cellularises [104,105].

So far as possible, the control logic of each component is programmed to reflect validated genetic interactions between the relevant *Drosophila* genes (Figure 1 provides a good overview). As in real embryos, these interactions are context-dependent, with “early network” interactions operating in the absence of Opa protein, and “late network” interactions operating in the presence of Opa [6]. Uncertain interactions and any *ad hoc* assumptions introduced for modelling purposes are clearly flagged within Supplement 2.

#### Given dynamic inputs, the model accurately recapitulates pair-rule patterning

If I run the model with static gap inputs, it performs very poorly (Figure 5A; Movie 9). This is unsurprising, because only four distinct states are present in the initial conditions, precluding fine scale patterning. Specifically, offsets between Prd/Slp and Ftz/Odd never emerge, and En expression is consequently absent from the final pattern. Moreover, the periodicity of this final pattern remains entirely double segmental.

**Figure 5:**
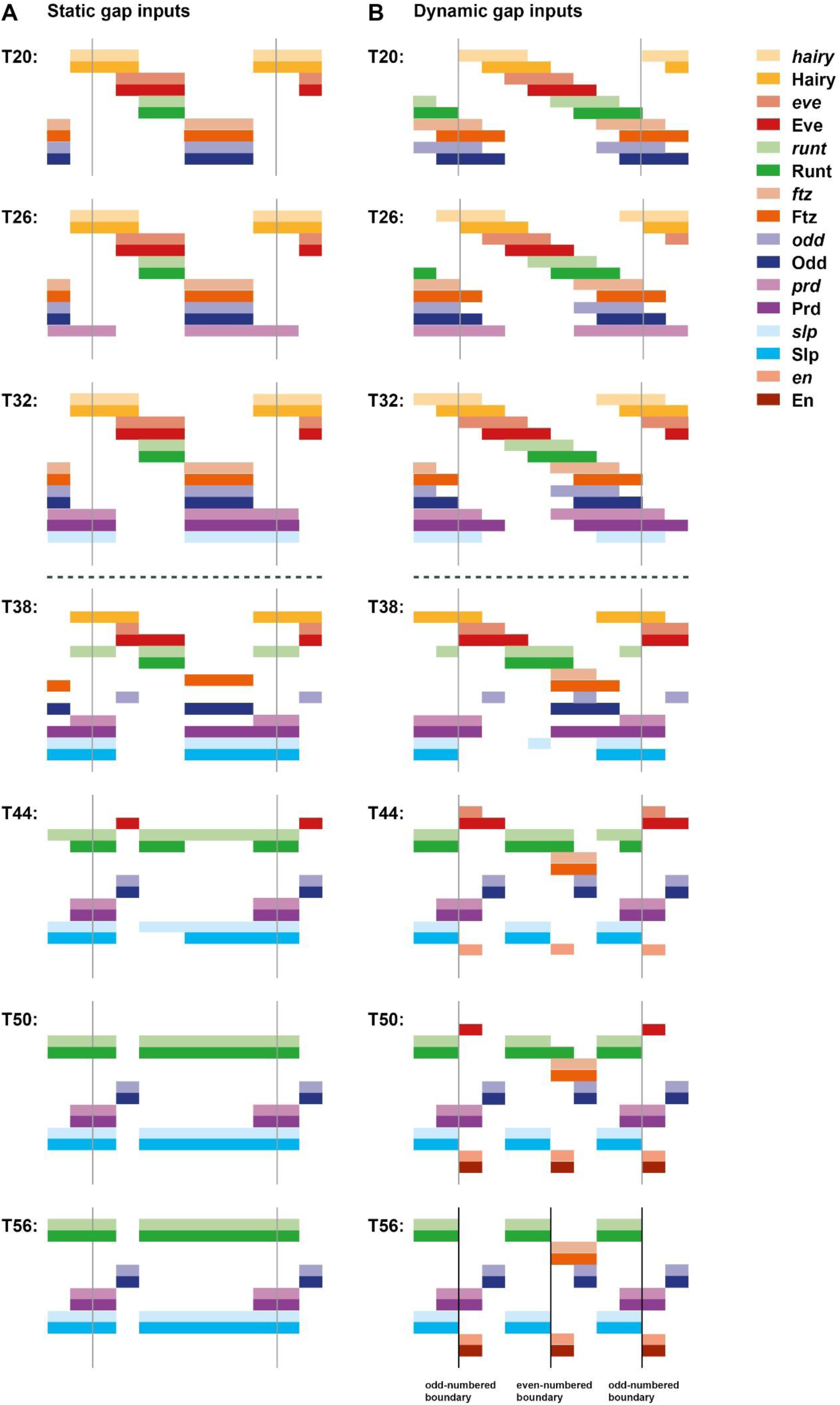
Given dynamic gap inputs, a Boolean model of the pair-rule network produces an appropriate segmental pattern. Pair-rule gene expression patterns generated by simulating a Boolean model of the pair-rule network, assuming either static (**A**) or dynamic (**B**) “gap” inputs (see Movies 9 and 10 for the original simulations). In each panel, the horizontal axis represents the AP axis (anterior left), while the vertical axis represents the different gene products that might be expressed in a given “cell” (column). Pale colours represent active transcription; dark colours represent active protein (see colour key at top right for details). Grey vertical lines indicate the span of an idealised double segment repeat of 8 “cells”. The same seven timepoints are shown for each simulation; the gap of 6 timesteps between each pictured timepoint is equivalent to the synthesis / decay time delay used for the simulations. The transition from the “early” network to the “late” network occurs between the third (T32) and fourth (T38) panels (dashed horizontal line). In the first panels (T20, equivalent to the situation shown in Figure 4), the pair-rule stripes of the primary pair-rule genes have been established, but the secondary pair-rule genes are still repressed. *prd* and *slp* expression is first seen in the second (T26) and third (T32) panels, respectively. Expression patterns change significantly in the fourth panels (T38), due to the new regulatory logic from the late network. These patterns resolve over the remaining timepoints. By the final panel (T56), both simulations have reached steady state. **A**: En is not expressed at any point; the final, stable pattern is broad pair-rule stripes of Slp alternating with narrow pair-rule stripes of Odd. **B**: An appropriate segment-polarity pattern is produced by the end of the simulation (compare the stable patterns of En, Odd, and Slp in the final panel with the schematic in Figure 3A). The black vertical lines in the final panel (T56) indicate the locations of prospective “parasegment boundaries” (i.e. interfaces between abutting domains of Slp and En expression).

However, when the gap inputs are programmed to shift anteriorly over time, the model recapitulates wild-type gene expression remarkably well (Figure 5B; Figure 6; Figure 6-figure supplement 1; Movie 10). The key patterning events all occur: 1) Prd/Slp and Ftz/Odd offsets both emerge, with Prd activating odd-numbered En stripes, and Ftz activating even-numbered En stripes. 2) Odd, Runt, and Slp all correctly transition from double-to single segment periodicity after the appearance of Opa. 3) An accurate final segment-polarity pattern is produced, consisting of mutually exclusive domains of En, Odd, and Slp within each segment (compare Figure 3A). Surprisingly for such a simple model, a number of fairly subtle features of pair-rule gene expression are also reproduced: for example, the segmental Runt stripes are initially of uneven width, requiring En activity to refine completely [106].

**Figure 6:**
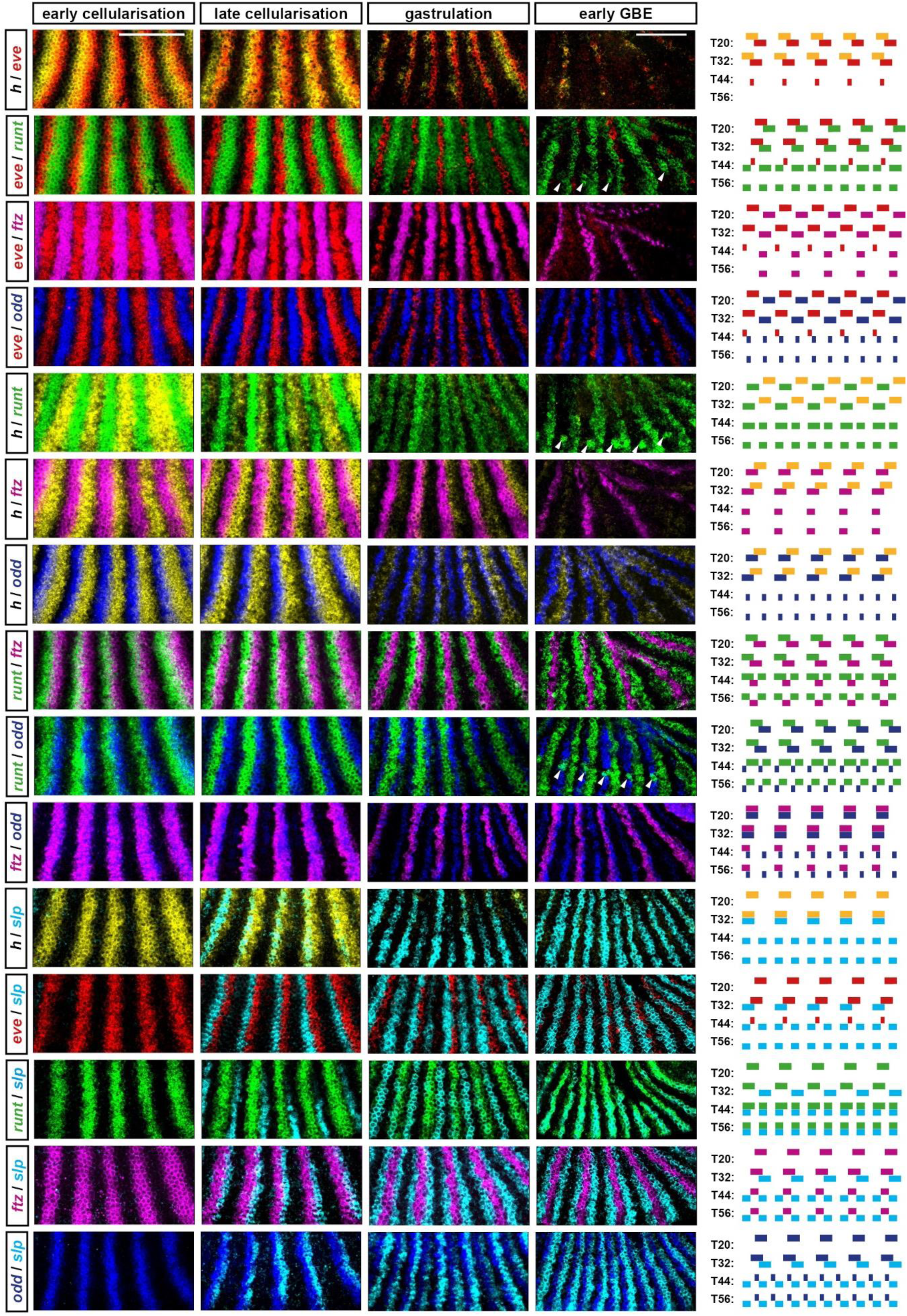
The “dynamic” simulation accurately recapitulates the spatiotemporal expression of the pair-rule genes. This figure compares the observed spatiotemporal expression patterns of *hairy, eve, runt, ftz, odd,* and *slp* (i.e. all the pair-rule genes except *prd*) with the patterns generated by the dynamic simulation shown in Figure 5B and Movie 10. Each row shows one of the 15 possible pairwise combinations of the six genes. On the left, double fluorescent *in situ* images are shown of the stripe 2-6 region of embryos of four different ages: early cellularisation, late cellularisation, gastrulation, and early germband extension (GBE). (Uncropped views of all the individual embryos used are shown in Figure 6–figure supplement 1.) Anterior is to the left, different transcripts are assigned different colours (yellow=*hairy*, red=*eve*; green=*runt*; magenta=*ftz*; blue=*odd*; cyan=*slp*). On the right, simulated transcriptional output of the relevant pair of genes is shown for four timepoints: T20, T32, T44, and T56 (stripe phasings are identical to those in Figure 5B). The four timepoints from the simulation can be thought of as roughly equivalent to the four developmental ages shown in the *in situ* images; in general, the simulation accurately recapitulates the observed patterns. Scale bars 100 μm; note that a slightly different scale is used for the “early GBE” *in situ* images, to compensate for cell rearrangements and morphogenetic movements that stretch the overall pattern during germband extension. Note also that early neurogenic *runt* expression (arrowheads) is apparent in some of the early germband extension images, in addition to the segmental stripes that are the focus of this project.

The only significant deviation from wild-type pair-rule gene expression relates to the late patterning of *prd* (Figure 6-figure supplement 2). The “A” stripes of *prd* (i.e. the narrow stripes formed from anterior portions of the early broad stripes) do not emerge and therefore the *prd* pattern remains pair-rule throughout. This failure to recover the wild-type pattern probably stems from the simplicity of the model, and indicates that additional complexities influence *prd* regulation in real embryos. These could include 1) additional spatial or temporal regulatory inputs missing from the model, 2) quantitative information from existing spatial or temporal inputs that is not captured by the use of Boolean variables, or 3) differential synthesis/degradation rates of particular segmentation genes not accounted for by the parsimonious equal rates used by the model (see Supplement 2).

The discrepancy from wild-type expression identifies *prd* as a gene whose regulation is currently not adequately understood, and should therefore be the focus of future study. Nevertheless, the model, although unrealistically simplistic, still captures most of the important aspects of pair-rule patterning. Indeed, the *prd* “P” stripes – which are patterned fine by the model – are the only *prd* expression domains reflected in the cuticle phenotype of *prd* mutant embryos, as the “A” stripes have relatively minor effects on segment-polarity gene expression [35,62,107].

The speed of patterning by the model is also appropriate to the real pair-rule system. Assuming a synthesis/decay time delay somewhere between 6 and 10 minutes [83,86] the simulation requires a minimum of 42-70 minutes to complete patterning (and could go slightly faster if provided with more extensive gap inputs, see Discussion). These numbers are consistent with the duration of the process in the *Drosophila* embryos, where everything happens within the roughly 1 hour (at 25° C) period from the beginning of cycle 14 (Bownes stage 5) through to the end of gastrulation (Bownes stages 6 and 7) [108,109].

In summary, a Boolean formulation of the pair-rule network is able to faithfully recapitulate pair-rule gene expression and successfully pattern both odd-numbered and even-numbered sets of parasegment boundaries, but only when provided with dynamic spatial inputs. To determine why this is so, I look in turn at the patterning of the odd-numbered boundaries and that of the even-numbered parasegment boundaries, both of which fail to emerge in the static scenario.

#### Dynamic patterning of the odd-numbered parasegment boundaries

As mentioned above, accounting for the patterning of the odd-numbered parasegment boundaries consists in explaining the relative expression of Eve, Prd, and Slp. The relevant patterning occurs towards the end of cellularisation; odd-numbered parasegment boundaries are present in *opa* mutant embryos, and therefore do not require late, Opa-dependent regulatory interactions [6,80,110].

At the end of cellularisation, Slp and Eve are expressed in directly abutting domains, with domains of Prd expression spanning the boundary between them [111]. Together, the expression domains of these three factors are sufficient to pattern mutually-exclusive, single cell wide domains of wg, *en,* and *odd* (Figure 7A; Figure 7–supplementary figure 1).

**Figure 7:**
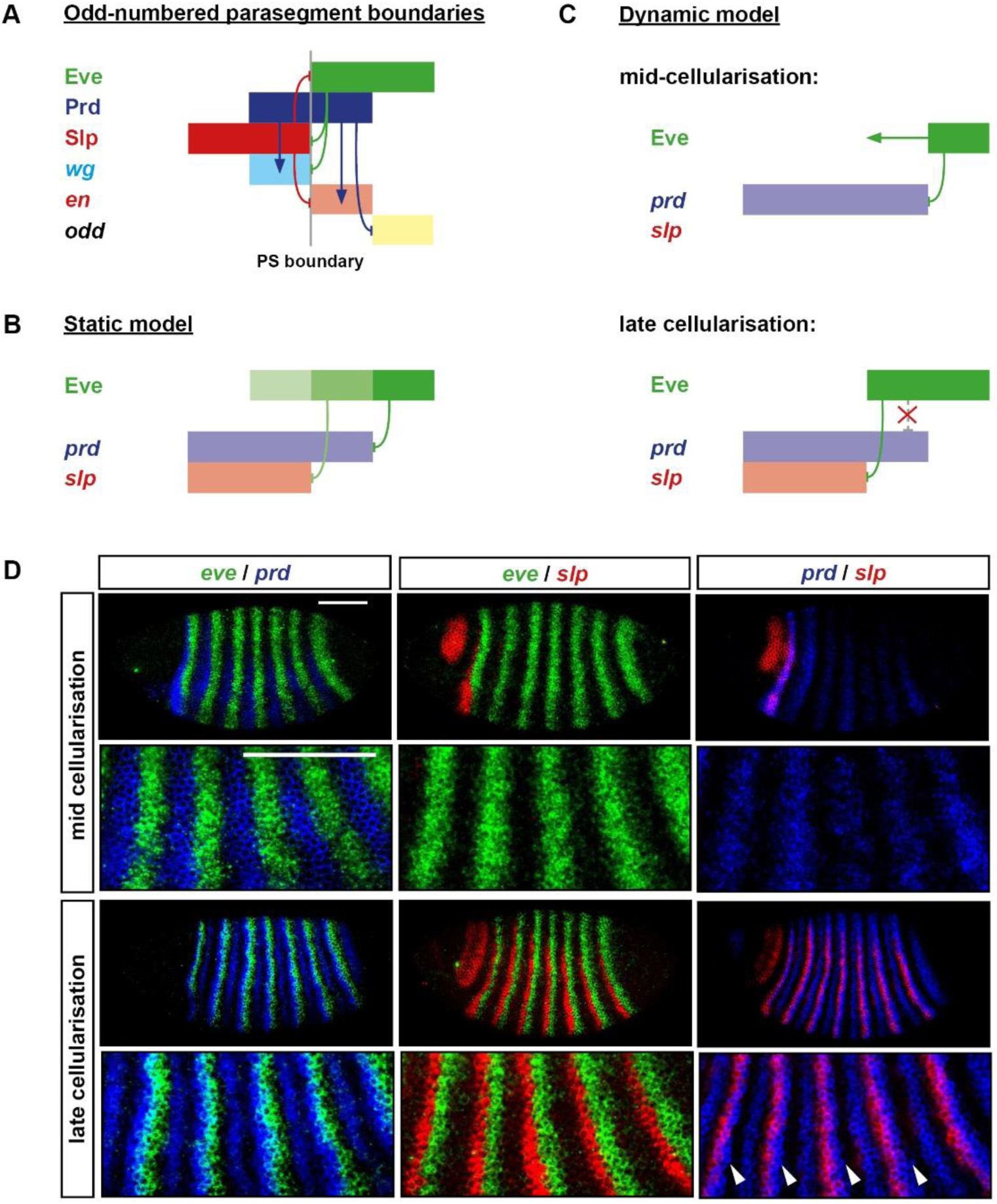
Dynamic patterning of the odd-numbered parasegment boundaries. **A:** Schematic of gene expression and regulation at the odd-numbered parasegment boundaries, where domains of Slp, Prd, and Eve expression pattern segment-polarity stripes of *wg, en,* and *odd.* Anterior left; hammerhead arrows represent repressive interactions; pointed arrows represent activatory interactions; dark colours represent protein expression; pale colours represent transcript expression; gray vertical line represents a prospective parasegment boundary. See Figure 7–figure supplement 1 for relevant *in situ* data. **B:** Static, “morphogen gradient” model for the patterning of the *prd* and *slp* posterior borders by Eve. The anterior margin of the Eve stripe is graded, with higher levels of Eve protein (darker green) present more posteriorly. High Eve (dark green) is required to repress *prd,* but only medium Eve (medium green) is required to repress *slp.* Based on Fujioka et al. 1995; Jaynes & Fujioka 2004. **C:** Dynamic model for the patterning of the *prd* and *slp* posterior borders by Eve. *prd* is activated earlier (mid-cellularisation) than *slp* (late cellularisation). In between these time points, the anterior border of the Eve domain shifts anteriorly. The posterior border of the *prd* domain is patterned by Eve at mid-cellularisation, but the posterior border of the *slp* domain is patterned by Eve at late cellularisation, resulting in a more anterior location. *prd* is no longer repressed by Eve at late cellularisation, resulting in stable, overlapping expression of Eve and *prd.* **D:** Double fluorescent *in situs* showing the relative phasing of *eve* (green), *prd* (blue), and *slp* (red) expression domains at mid-cellularisation and late cellularisation. Anterior left, dorsal top; enlarged views of stripes 2-6 are shown below whole embryo lateral views. At mid-cellularisation, *slp* is not expressed and the posterior borders of the *prd* stripes abut the anterior borders of the *eve* stripes. At late cellularisation, the posterior borders of the *prd* stripes overlap the anterior borders of *eve* stripes (note the regions that appear cyan), the posterior borders of the *slp* stripes sharply abut the anterior borders of the *eve* stripes, and the posterior borders of the *slp* stripes are offset anteriorly from the posterior borders of the *prd* stripes (arrowheads). These expression patterns seem more consistent with the dynamic model (**C**) than the static model (**B**). Scale bars 100 μm.

The regulatory interactions relevant to setting up the Slp/Prd/Eve pattern are as follows. Once Slp protein has been synthesised, Eve and Slp mutually repress each other (explaining the sharp boundaries between their respective stripes, as seen in Figure 7D). *prd* is at first repressed by Eve, which patterns its initial broad stripes during mid-cellularisation [58], but later becomes insensitive to Eve activity [61]; the cause of this regulatory change is currently unknown). Finally, *prd* is activated within the blastoderm around 10 minutes earlier than is *slp* (Figure 7D; [74]).

Given these interactions, it is easy to see why shifting Eve stripes are important for the formation of the odd-numbered parasegment boundaries. In the static case, posterior boundaries of *prd* and *slp* will coincide so long as Eve activity is effectively Boolean. However, because *prd* and *slp* have different temporal windows during which they are sensitive to Eve repression, dynamic Eve stripes provide an opportunity for differential spatial patterning. The posterior borders of the Prd stripes (which seem to be static once formed [74]) are patterned considerably earlier than the posterior borders of the Slp stripes (also static once formed), and thus correspond to slightly more posterior locations of the Eve stripe anterior boundaries.

Under this view, the observed one-nucleus-wide offsets between the *prd* and *slp* posterior boundaries correspond to one-nucleus shifts in the Eve stripe boundaries in between the patterning of *prd* and *slp* (Figure 7C), rather than one-nucleus-wide distances between specific Eve activity thresholds (Figure 7B). Consistent with this interpretation, the posterior boundaries of the *prd* stripes at first abut the anterior boundaries of the *eve* stripes and it is only later, when *slp* starts to be expressed, that overlaps between *prd* and *eve* become obvious (Figure 7D). In contrast, the morphogen gradient hypothesis (Figure 7B) would predict overlapping expression of *eve* and *prd* from the beginning.

In summary, the dynamic model suggests that the root causes for the spatial patterning of the odd-numbered parasegment boundaries lie in 1) the differential temporal regulation of *prd* and *slp* expression and 2) the temporal restriction of Eve repressive activity on *prd.* Both of these temporal phenomena are documented, but neither is currently explained.

#### Dynamic patterning of the even-numbered parasegment boundaries

The even-numbered parasegment boundaries are patterned during gastrulation, requiring Opa-dependent regulatory interactions [6]. The relevant regulatory inputs are provided by partially overlapping domains of Runt, Ftz, and Slp (Figure 8B). The patterning of the even-numbered parasegment boundaries thus relies on establishing an appropriate *runt* / *ftz* / *slp* pattern during cellularisation (Figure 8A).

**Figure 8:**
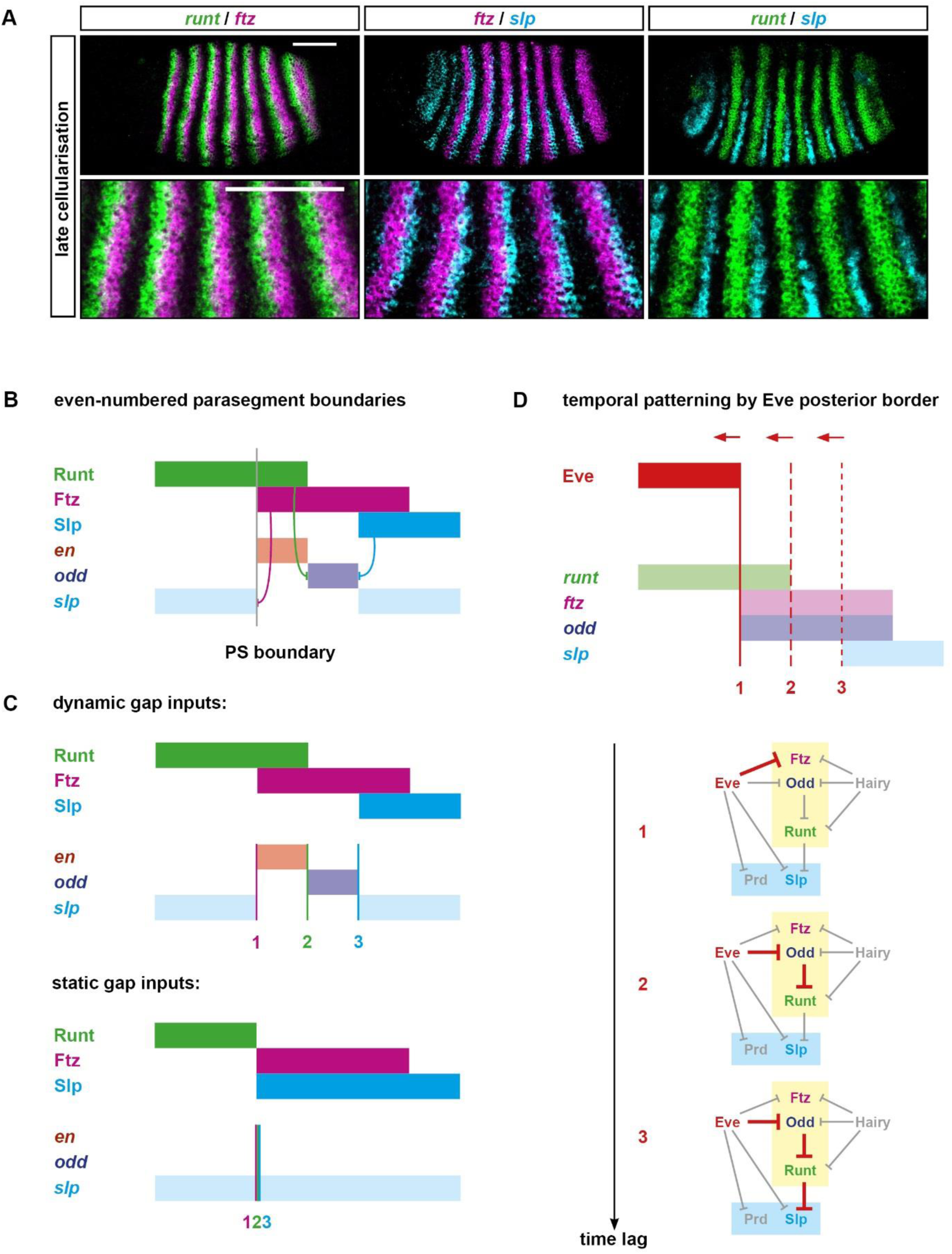
Dynamic patterning of the even-numbered parasegment boundaries. **A**: Double fluorescent in situs showing the relative expression of *runt, ftz,* and *slp* at late cellularisation. Enlarged views of stripes 2-6 are shown below whole embryo lateral views (anterior left, dorsal top). The *ftz* stripes overlap the posteriors of the *runt* stripes at their anteriors, and overlap the anteriors of the *slp* stripes at their posteriors. The posterior borders of the *runt* stripes are just slightly anterior to the anterior borders of the *slp* stripes. Scale bars 100 μm. **B**: Schematic showing the patterning of the even-numbered parasegment boundaries. At gastrulation, Runt, Ftz, and Slp are expressed in partially overlapping domains similar to their transcript expression at late cellularisation (see **A**). These overlapping domains provide a template for the segment-polarity stripes of *en, odd,* and *slp:* in particular, the anterior borders of the Ftz stripes define the posterior borders of the *slp* secondary stripes, the posterior border of the Runt stripes define the anterior borders of the *odd* primary stripes, and the Slp anterior borders define the posterior borders of the *odd* primary stripes. The even-numbered *en* stripes are activated by Ftz, but repressed by Odd and Slp, and so are restricted to the region of overlap between Runt and Ftz, where both *odd* and *slp* are repressed [6]. Hammerhead arrows represent repressive interactions; grey vertical line represents a prospective parasegment boundary; anterior left. **C**: Schematic explaining why the even-numbered parasegment boundaries require dynamic gap inputs in order to be patterned. Given dynamic inputs (top panel, see Figure 5B), the Ftz anterior boundary (1, pink vertical line), the Runt posterior boundary (2, green vertical line), and the Slp anterior boundary (3, blue vertical line) are each located at different AP positions (as in **B**), resulting in the segment-polarity pattern: *slp, en, odd, slp.* Given static inputs (bottom panel, see Figure 5A), all three boundaries will coincide, resulting only in broad *slp* expression. **D**: Schematic explaining the origin of the offset boundaries of *ftz, runt,* and *slp.* Top: the relative expression of Eve, *runt, ftz, odd,* and *slp* is shown at late cellularisation (equivalent to panel **A**, or T32 in Figure 5B). The solid red vertical line indicates the current position of the Eve posterior border, which coincides with the *ftz* anterior border (1). Dotted red vertical lines indicate previous positions of the dynamic Eve posterior border, coinciding with the *runt* posterior border (2), or the *slp* anterior border (3), respectively. Bottom: the regulatory chains responsible for patterning each of the three expression boundaries are highlighted in red on the early pair-rule network. All three boundaries trace back to Eve, but more posterior boundaries correspond to longer regulatory chains and so would incur a longer time lag to resolve, given a change in Eve expression. The three different genes (*ftz*, *runt,* and *slp*) are effectively patterned by increasingly earlier incarnations of the Eve stripes, and therefore the existence of spatial offsets between boundaries 1, 2, and 3 relies on the Eve posterior border shifting anteriorly over time.

As seen earlier in Figure 5, setting up the correct phasing of these three expression domains requires dynamic gap inputs: in their absence, rather than there existing three distinct expression boundaries (Ftz anterior; Runt posterior; Slp anterior), all three boundaries coincide (Figure 8C). Consequently, rather than these domains collectively specifying four distinct output states (Slp; En; Odd; Slp), the starting conditions for the En and Odd states are absent from the initial pattern, and the result is just broad expression of Slp.

Why are these important boundaries offset in the dynamic case, but coincident in the static case? It turns out that they all trace back eventually to the same upstream input, the posterior boundary of Eve expression (Figure 8D). *ftz* is directly repressed by Eve; *runt* is repressed by Odd, which is itself repressed by Eve; *slp* is repressed by Runt, whose regulation again traces back to Eve via Odd. Therefore, in the static case where the system is allowed to reach stable state, all three of these boundaries will correspond with the static posterior boundaries of the Eve stripes (see Figure 5A). However, each of the three regulatory chains is of a different length (1, 2, or 3 interactions, respectively), and will therefore take a different amount of time to conclude after a change in Eve expression. In the dynamic case where the Eve posterior boundaries are constantly shifting anteriorly, the resulting expression boundaries of Ftz, Runt, and Slp will all end up offset from the Eve boundary by a slightly different amount, producing the template for a high-resolution spatial pattern.

This patterning mechanism has two important implications. First, transient expression states generated by the early network are responsible for specifying crucial segment-polarity cell fates. The unstable combination of Ftz and Runt is required to specify *en* expression, while the unstable state of Ftz alone (without Runt or Slp) is required to specify *odd* expression. Second, Odd is an indirect autorepressor over the course of segmentation as a whole: during cellularisation, Odd represses *runt* and Runt represses *slp;* then during gastrulation Slp represses *odd.* However, this regulatory chain is only completed in the posteriors of the Odd stripes (where Odd expression is older) explaining why these stripes narrow from the posterior rather than being lost outright.

### 5. Patterning dynamics explain the severity of the *eve* mutant phenotype

#### Eve is responsible for patterning most of the expression boundaries in the final segment pattern

In summary, under the dynamic model, most of the expression boundaries in the final segment pattern eventually trace back to Eve expression boundaries (Figure 9). Via direct regulation of *prd* and *slp,* the anterior borders of the Eve stripes specify both anterior and posterior borders of the odd-numbered *en* stripes, defining the transitions between three segment polarity states: Slp; En; Odd. Via direct regulation of *ftz* and *odd,* and hence indirect regulation of *runt* and *slp,* the posterior borders of the Eve stripes specify both the anterior and posterior borders of the even-numbered *en* stripes, and also the anterior borders of the *slp* primary stripes, defining the transitions between four segment polarity states: Slp; En; Odd; Slp.

**Figure 9:**
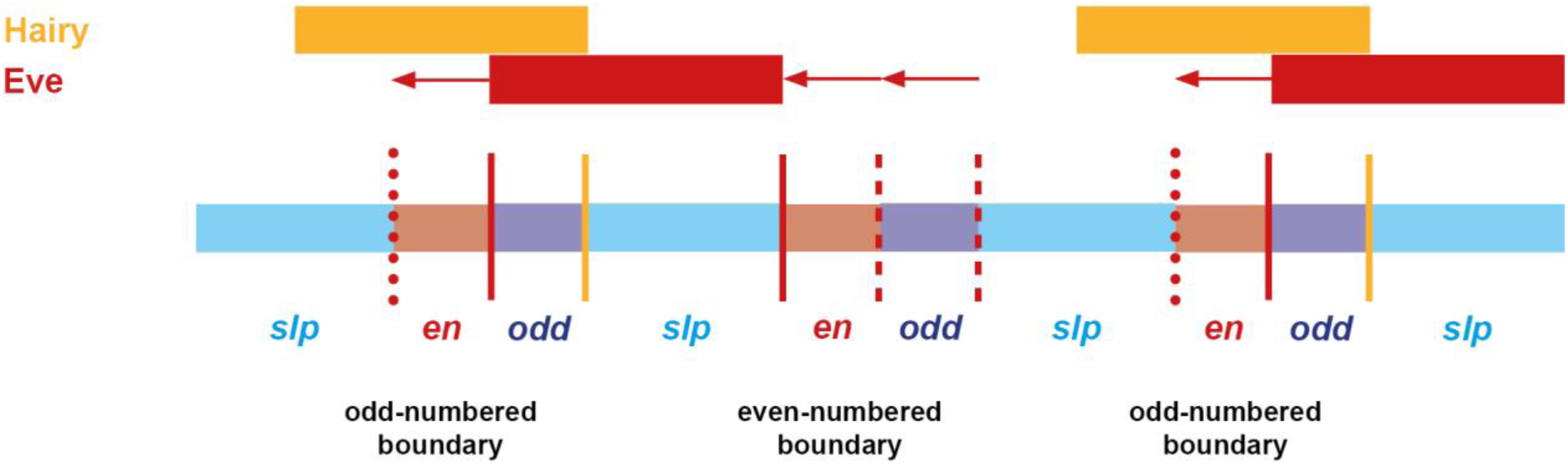
Most of the expression boundaries in the final pattern trace back to the Eve stripes. Schematic showing the relationship between the final pattern and the original signals from Hairy and Eve, according to the model. At the top of the schematic is shown the relative expression of Hairy and Eve domains at the end of the “early network” phase of patterning (i.e. equivalent to T32 in Figure 5B, or late cellularisation in a real embryo). At the bottom of the schematic is shown the final output generated by the “late network” phase of patterning (i.e. equivalent to T56 in Figure 5B, or early germband extension in a real embryo – see Figure 3A). Red vertical lines indicate expression boundaries in the final pattern that trace back to Eve, while yellow lines indicate those that trace back to Hairy. Within a double segment repeat, only the anterior boundaries of the *slp* secondary stripes trace back to Hairy (Hairy represses Runt, Runt represses late Eve, Eve represses Slp). The remaining five boundaries within a double segment repeat trace back to Eve: two are specified by the Eve anterior boundaries (see Figure 7) while three are specified by the Eve posterior boundaries (see Figure 8). Solid lines indicate boundaries that map directly to the Hairy/Eve domains shown above. Dashed red lines refer to past locations of the Eve posterior boundaries. Dotted red lines refer to the future locations of the Eve anterior boundaries. Red horizontal arrows indicate regions where dynamic Eve expression is important for patterning.

The remaining expression boundary in the final pattern – the anterior border of the *slp* secondary stripes – is the only one that traces back to Hairy. Repression from Hairy defines the anterior boundary of the *runt* primary stripes, which overlap Eve expression (see Figure 4C). At gastrulation, Runt represses the posteriors of the Eve stripes, permitting the expression of *slp,* which is repressed by Eve more anteriorly (see Figure 5B).

In previous models of pair-rule patterning, one of the main motivations for positing spatially graded activity of the Eve stripes (Figure 3C) was the realisation that many different expression boundaries (including both parasegment boundaries) within a double-segment repeat are reliant on Eve. However, as described above, the dynamic model can explain the same findings invoking only Boolean Eve activity, indicating that quantitative effects may not be essential to the patterning mechanisms operating within the blastoderm.

#### The dynamic model faithfully recapitulates the eve mutant phenotype

An informative additional test of the dynamic model is to see how well it predicts the expression dynamics and resulting cuticle phenotype of *eve* mutant embryos, given that these have been hard to explain using traditional patterning models.

Although *eve* was originally identified as a pair-rule gene on the basis of a pair-rule cuticle phenotype [1], it turned out that this particular mutant allele was an *eve* hypomorph, while *eve* null mutants yield an aperiodic denticle lawn phenotype instead [112]. Both odd-numbered and even-numbered *en* stripes are absent from *eve* null mutant embryos [113], indicating severe mispatterning of upstream pair-rule gene expression.

I carried out a number of double fluorescent in situs to characterise the development of pair-rule gene expression patterns in *eve* mutant embryos (Figures 10; Figure 11; see also Supplement 1). The results are largely in accordance with earlier, more fragmentary characterisations of gene expression in these mutants [33,34,58,61,106,114].

**Figure 10:**
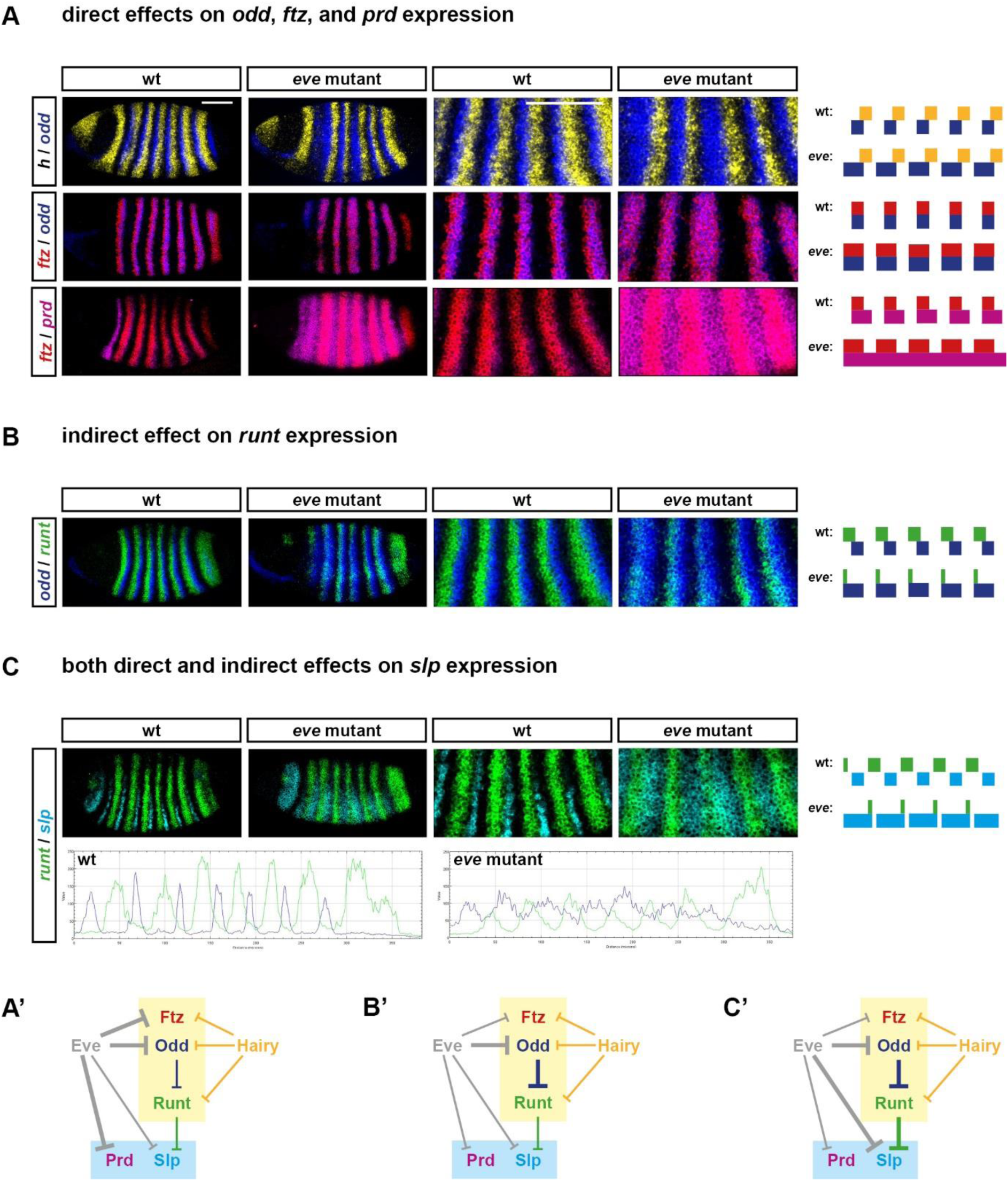
Altered pair-rule gene expression in *eve* mutant embryos during cellularisation. **A**-**C**: *In situ* images of pair-rule gene expression in cellularisation stage wild-type and *eve* mutant embryos (left), compared to simulated transcriptional output from “wild-type” and “*eve* mutant” pair-rule networks (right). Both whole embryos and enlarged views of stripes 2-6 are shown. The simulated expression patterns shown are equivalent to T32 in Figure 12. Transcripts of each pair-rule gene are shown in a different colour (yellow=*hairy*, green=*runt*, red=*ftz*, blue=*odd*, magenta=*prd*, cyan=*slp*). Scale bars 100 μm. **A’**-**C’**: Regulatory interactions relevant to the aberrant expression patterns in *eve* mutants are highlighted on the early pair-rule network (bold arrows). Eve and its regulatory effects, which are absent from the mutant embryos, are shown in grey on the network diagrams. **A**: Eve normally represses *ftz, odd,* and *prd* (**A’**). In *eve* mutant embryos, all three genes are ectopically expressed: the *ftz* and *odd* stripes expand anteriorly, and *prd* is expressed ubiquitously, rather than in stripes. These expression changes are recapitulated by the simulation. **B**: Eve normally indirectly regulates *runt* expression, by repressing its repressor, Odd (**B’**). In *eve* mutant embryos, *odd* expression expands anteriorly (see **A**), resulting in a downregulation of the *runt* stripes, except at their anterior margins (see also the intensity profiles in **C**). This effect is recapitulated in a discrete manner by the simulation. **C**: Eve normally regulates Slp in two ways (**C’**): 1) by repressing it directly, and 2) by repressing it indirectly, via indirectly maintaining the expression of its repressor, Runt (see **B**), via direct repression of *odd* (see **A**). (See also Figures 7 and 8.) In *eve* mutant embryos, *slp* is expressed fairly ubiquitously, rather than in narrow stripes. This expansion is evident in the simulated expression, although *slp* remains periodic overall. The difference between the observed and simulated expression can be explained by quantitative effects, which are not captured by the qualitative nature of the model. The plots below the *in situ* images show the AP intensity profiles of *runt* (green) and *slp* (blue) along a narrow ventral strip of the trunk of the two embryos pictured. In the wild-type embryo, the *runt* stripes are all (except stripe 7) roughly symmetrical and strongly expressed. In the *eve* mutant embryo, *runt* stripes 1-6 (which overlap with *odd* expression, see **B**) have much lower intensity than *runt* stripe 7 (which doesn’t overlap with odd expression, see **B**) and exhibit a sawtooth pattern, in which expression intensity decreases from anterior to posterior. The *slp* expression in the *eve* mutant embryo, while broad, does display a pair-rule modulation, which is in opposite phase to the downregulated *runt* stripes. Therefore, the same two regulatory interactions (repression of *runt* by Odd, and of *slp* by Runt) are evident in both the *in situ* data and the simulated data, but lead to slightly different expression patterns in each case, one quantitative and one qualitative.

**Figure 11:**
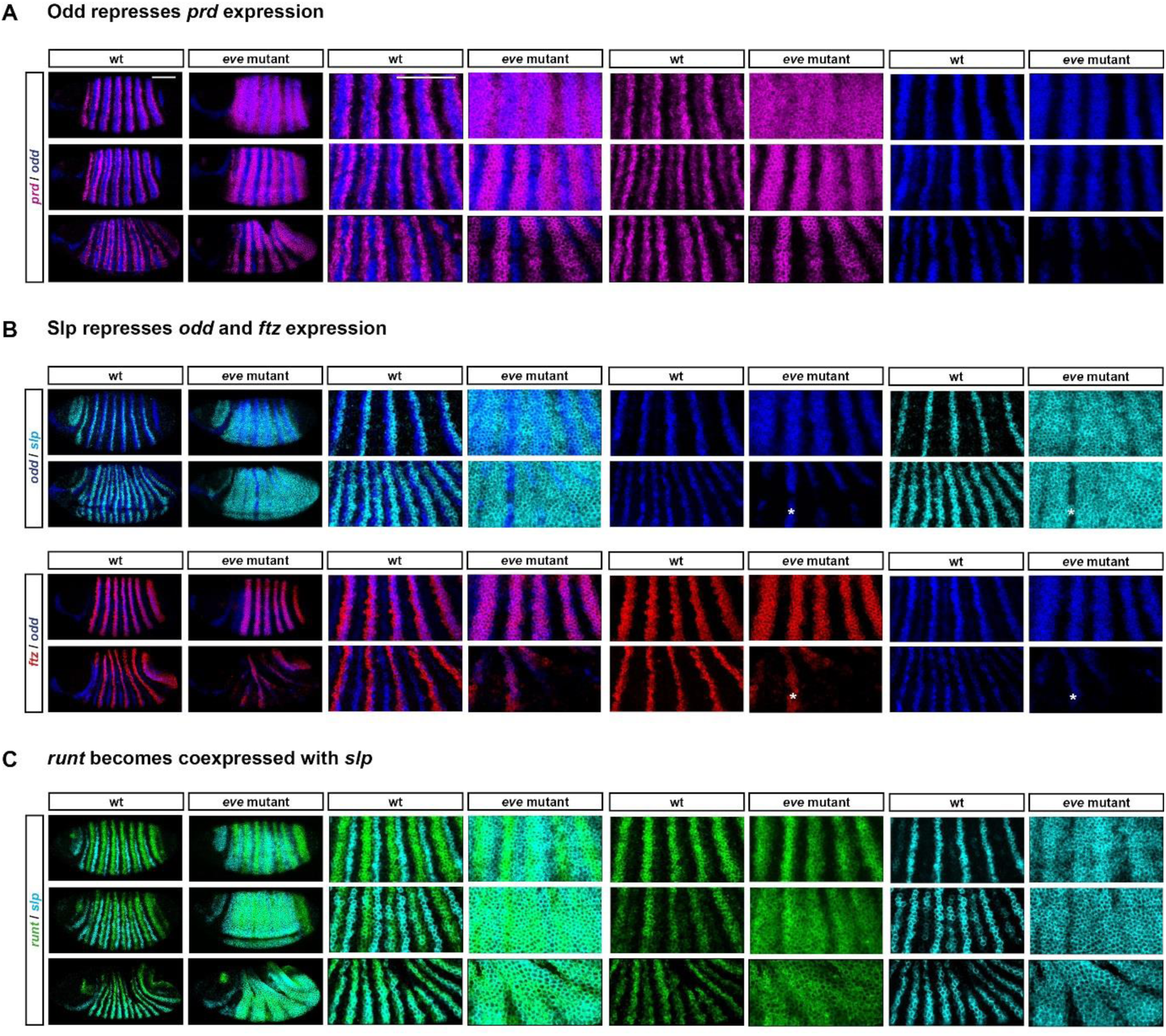
Altered pair-rule gene expression in *eve* mutant embryos during gastrulation. *In situ* images of pair-rule gene expression in wild-type and *eve* mutant embryos over the course of gastrulation. For each set of *in situs,* each row compares a wild-type and a mutant embryo of roughly equal age (age increases from top to bottom). From left to right, the individual panels show whole embryos (anterior left, dorsal top); enlarged views of stripes 2-6; and individual channels for the enlarged regions. Transcripts of different genes are shown in different colours (green=*runt*, red=*ftz*, blue=*odd*, magenta=*prd*, cyan=*slp*). In each case, pattern evolution occurs in a predictable manner over the course of gastrulation, given the structure of the late pair-rule network (Figure 1B) combined with the different starting conditions of each genotype. Scale bars 100 μm. **A**: Odd starts to repress *prd* at gastrulation, and so *prd* expression is lost from cells in which *odd* and *prd* expression initially overlap. In wild-type embryos, the *odd* primary stripes overlap the centres of the *prd* pair-rule stripes, which therefore split in two. In *eve* mutant embryos, broad *odd* stripes are overlain on initially aperiodic *prd* expression, which consequently resolves into a pair-rule pattern. **B**: Slp protein appears at the beginning of gastrulation, and represses both *odd* and *ftz.* In wild-type embryos, this causes the primary stripes of both *odd* and *ftz* to narrow from the posterior (where they overlap the *slp* primary stripes). In *eve* mutant embryos, *slp* is broadly expressed, causing general repression of *odd* and *ftz.* Note that both *odd* and *ftz* expression persists in stripe 3 (asterisks), corresponding with a gap in the *slp* expression domain. **C**: During gastrulation, both *slp* and *runt* are regulated similarly, and Slp represses all of the repressors of *runt* (see Figure 1B). Consequently, *runt* and *slp* take on almost identical expression patterns. In wild-type embryos, the two genes become expressed in coincident segmental stripes. In *eve* mutant embryos, early broad expression of *slp* allows *runt* to also become ubiquitously expressed. Note that the *slp* domain later resolves into a pair-rule pattern (likely due to repression from residual Ftz protein).

When compared with wild-type embryos of corresponding ages, a number of significant changes are obvious in the *eve* mutants. During cellularisation, the *odd* primary stripes are broader than normal (Figure 10A). When the secondary pair-rule genes first turn on, their expression is largely aperiodic, rather than pair-rule (Figure 10A,C). Finally, single-segmental patterns do not emerge at gastrulation: instead, *prd* resolves into broad pair-rule stripes (Figure 11A), *runt* and *slp* become expressed fairly ubiquitously (Figure 11B,C), and *ftz* and *odd* expression largely disappears (Figure 11B).

I then simulated *eve* “mutants” by starting with the dynamic model and simply setting *eve* transcription to always remain off (Figure 12B; Movie 11). Encouragingly, this simulation recapitulates the same changes to pair-rule gene expression as seen in the real embryos, indicating that the model should shed light on the aetiology of the mutant phenotype. Indeed, as described below, the observed expression changes follow logically from the structure of the pair-rule network, and the myriad direct and indirect roles of the Eve stripes.

**Figure 12:**
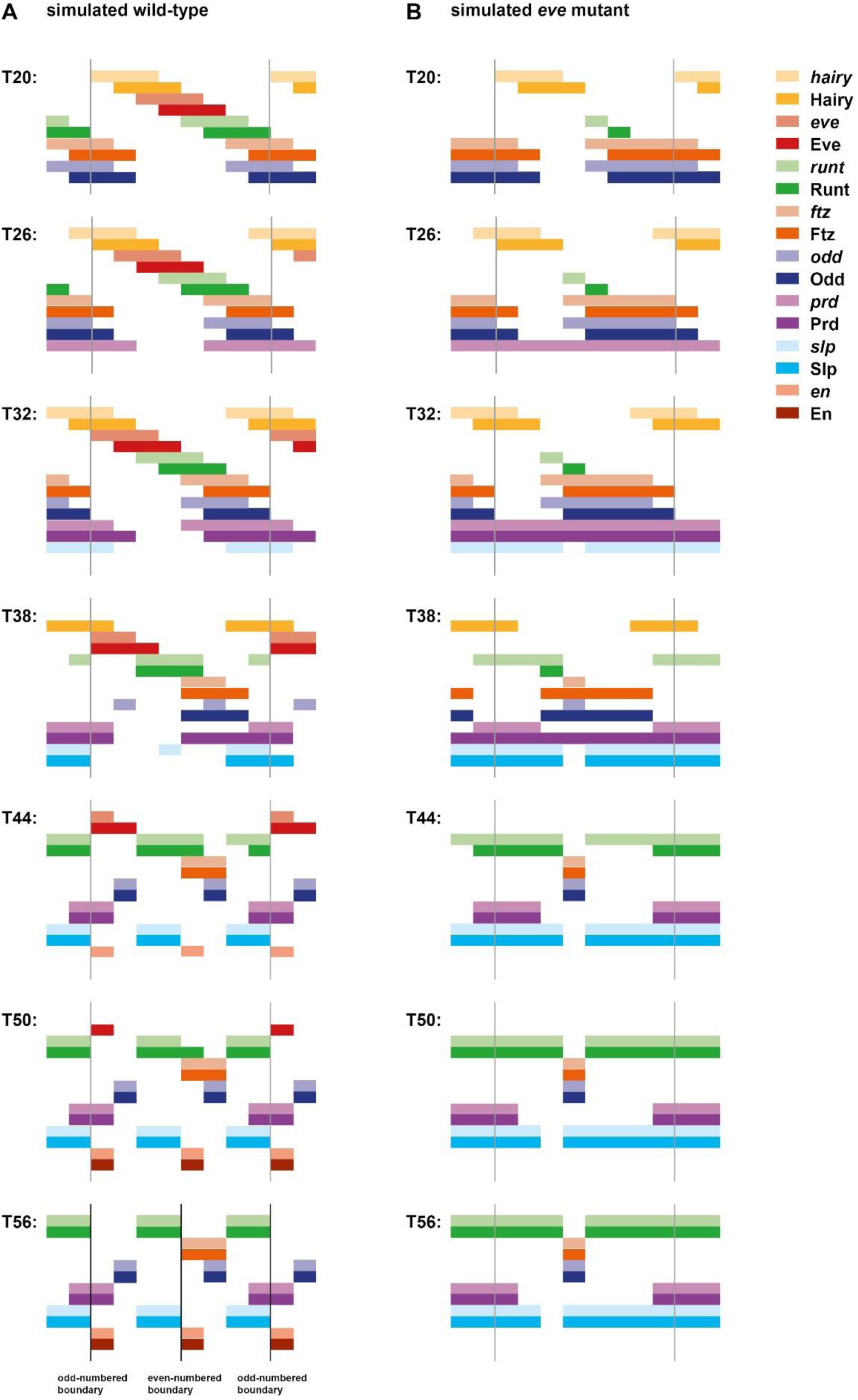
Simulated *eve* mutants recapitulate the *eve* mutant phenotype. Pair-rule gene expression patterns generated by simulating either the “wild-type” model of the pair-rule network (**A**, identical to Figure 5B) or an *“eve* mutant” network in which *eve* transcription is absent (**B**). See Movies 10 and 11 for the original simulations. In each panel, the horizontal axis represents the AP axis (anterior left), while the vertical axis represents the different gene products that might be expressed in a given “cell” (column). Pale colours represent active transcription; dark colours represent active protein (see colour key at top right for details). Grey vertical lines indicate the span of an idealised double segment repeat of 8 “cells”. The same seven timepoints are shown for each simulation (and match those in Figure 5). In the wild-type simulation (**A**), an appropriate segment-polarity pattern is produced by the end of the simulation, including segmental stripes of En expression, and two prospective parasegment boundaries (black vertical lines) within each double-segment repeat. In the *eve* mutant simulation (**B**), aberrant expression patterns are present at every timepoint. En expression and single-segment periodicity are absent from the final output, and no parasegment boundaries are patterned.

#### The aetiology of the eve mutant phenotype

Eve expression plays a relatively minor role in late patterning, and most of the expression changes seen in *eve* mutant embryos result from the loss of Eve activity during the early stages of patterning.

First, during mid-cellularisation, there are direct effects on *odd, ftz* and *prd,* all of which become ectopically expressed in regions where they would normally be repressed by Eve (Figure 10A). The *odd* and *ftz* primary stripes expand anteriorly, while the *prd* interstripes (i.e. the gaps between the early broad stripes) are completely absent.

The ectopic Odd expression has a knock-on effect on *runt* expression, which is largely repressed in the simulation, and generally down-regulated in the real embryos (Figure 10B). This loss of Runt activity contributes to the ectopic expression of *slp* that appears at the end of cellularisation: in the absence of its two repressors, Eve and Runt, *slp* becomes expressed almost ubiquitously throughout the trunk (Figure 10C).

Finally, the downstream effects of the aberrant patterns established during cellularisation play out over the course of gastrulation, after Opa-dependent regulatory interactions kick in. Repression from the broad pair-rule stripes of Odd and Ftz (respectively) cause the aperiodic domains of *prd* and *slp* to resolve into pair-rule patterns (Figure 11A,C). At the same time, *odd* and *ftz* are themselves repressed by the ectopic Slp expression in the embryo (Figure 11B). In the absence of its repressors Eve and Odd, *runt* expression becomes broadly expressed by the beginning of germband extension (Figure 11C).

As a consequence of all this mispatterning, *en* expression is completely repressed, and parasegment boundaries never form (Figure 12). The odd-numbered *en* stripes are blocked by the ectopic Slp expression that replaces the Eve stripes. In wild-type embryos, these *en* domains are specified by the short regulatory chain [Eve ---| Slp ---| En]. Therefore, in *eve*, *slp* double mutants they should reappear, as indeed they do [36,114]. On the other hand, the even-numbered *en* stripes are redundantly repressed in *eve* mutant embryos by both ectopic Odd and ectopic Slp, as a result of the regulatory chains [Eve ---| Odd ---| En] and [Eve ---| Odd ---| Runt ---| Slp ---| En]. Accordingly, they reappear in *eve, odd* double mutants [115], but not in *eve*, *slp* double mutants [36].

### 6. The Drosophila pair-rule network is compatible with both simultaneous and sequential segmentation

#### A single spatial input is sufficient to recapitulate Drosophila pair-rule patterning dynamics

By simulating the *Drosophila* early pair-rule network, I have shown how dynamic inputs from just two factors, Hairy and Eve, are sufficient to organise the expression of the system as a whole. In the *Drosophila* blastoderm, these inputs are driven by the dynamic output of the posterior gap system. However, elaborate control of pair-rule gene expression by gap factors is likely a relatively recent novelty in arthropod segment patterning, originating during the evolutionary transition from short-germ to long-germ embryogenesis. I therefore explored ways in which the patterning potential of the *Drosophila* pair-rule network might be preserved in the absence of explicit gap gene inputs.

The relationship between expression dynamics and regulatory control logic is one-to-many: it is possible for distinct regulatory networks to produce identical output. (Indeed, resultant “developmental systems drift” is likely to be a widespread phenomenon, [116]). Therefore, any given control logic that preserves the wild-type dynamics of the Hairy and Eve stripes should theoretically be able to substitute for their actual regulation by gap factors, without perturbing downstream pair-rule gene expression.

A plausible example of an alternative network is shown in Figure 13–figure supplement 1. In this scenario, gap inputs into Eve are replaced by repression from Runt and Odd, both regulatory interactions that occur in the late network (see Figure 1B). In contrast, Hairy remains an “input-only” factor in the pair-rule network, but instead of being regulated by gap factors, it now represses itself, leading to cyclic expression. Autorepression is common for *her/hes* family genes, and notably is important for vertebrate somitogenesis [117,118]. I further simplify the system by tying the onset of Opa expression to the decay of signal X, by making X repress Opa (see Supplement 2). Everything else, including the late network, is unchanged.

If the synthesis and decay rates of Hairy expression are tuned in the model in such a way that the Hairy oscillation cycle matches the shift rate in the original model, and the simulation is initialised with a repeating posterior-to-anterior phase gradient of Hairy expression across the tissue (Figure 13A; Movie 12), the alternative network generates exactly the same behaviour as the original model, with the exceptions that 1) *hairy* transcripts are now completely out of phase with Hairy protein rather than broadly in phase, and 2) the initial pair-rule pattern takes longer to establish during the early phase of the simulation (compare with Figure 5B and Movie 9).

**Figure 13:**
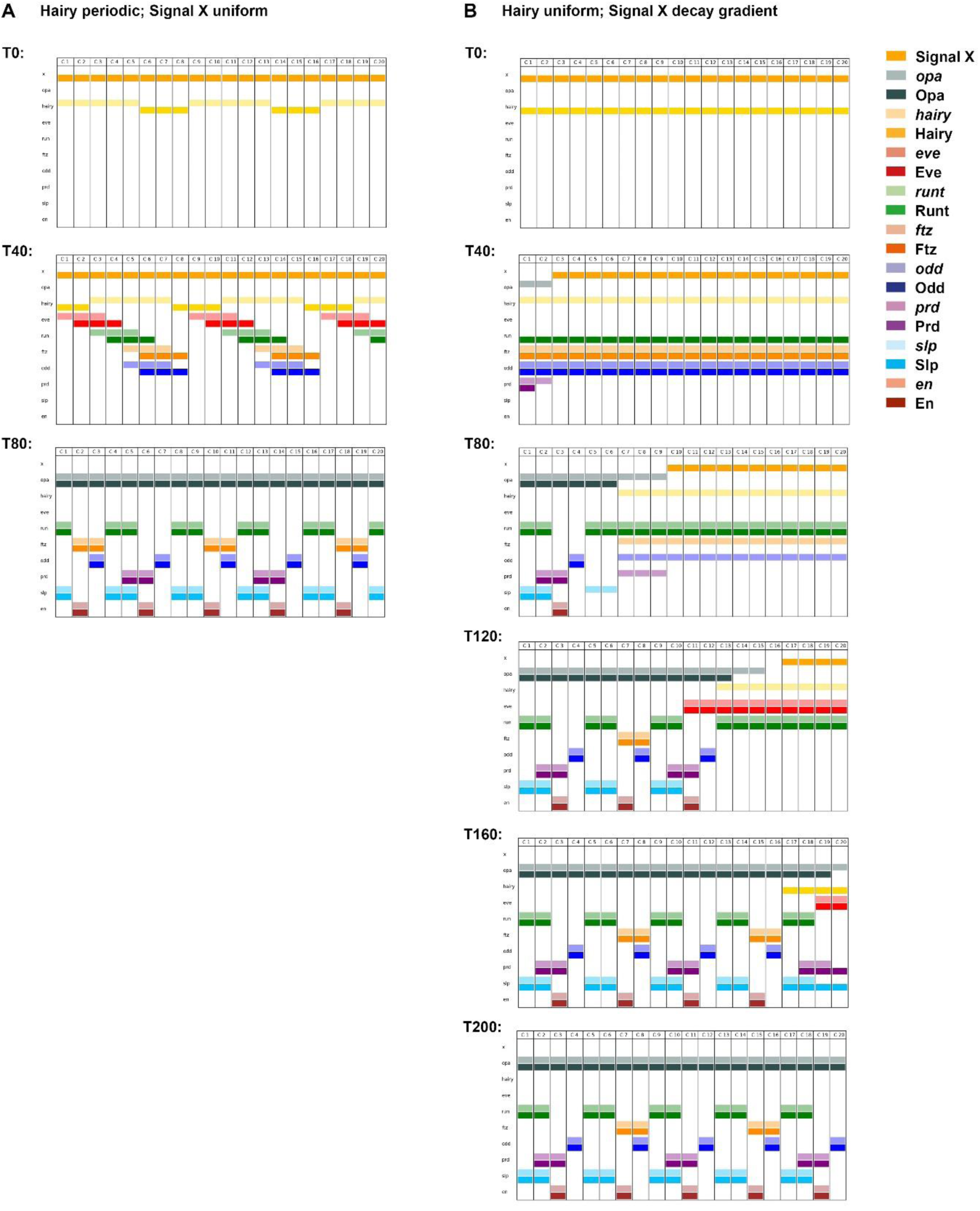
A slightly modified pair-rule network can pattern segments in both simultaneous and sequential modes. **A**,**B**: Simulation output for a slightly modified version of the pair-rule network, in which gap regulation of *hairy* and *eve* is replaced by additional cross-regulatory interactions within the early network (see Figure 13–figure supplement 1), and Opa, which controls the time of transition to the late network, is regulated by Signal X. Each panel shows the system state at a specific timepoint between TO and T200 (see Movies 12 and 13 for complete output). In each panel, the horizontal axis represents a region of AP axis (anterior left) and the vertical axis represents the different gene products that might be expressed in a given “cell” (individual columns C1 to C20). Pale colours represent active transcription; dark colours represent protein activity (see colour key for details). In **A**, the system is initialised with uniform expression of Signal X, and a periodic phase gradient of Hairy expression (protein or transcript “age” increases from posterior-to-anterior, and the pattern repeats every 8 cells – see Supplement 2 for details). Given these starting conditions, the dynamical behaviour of the system is almost identical to the unmodified network (compare Figure 5B or Movie 10). Note however that *hairy* transcript is always out of phase with Hairy protein. In **B**, the system is initialised with uniform expression of Hairy, but a decay gradient of Signal X (protein age increases from posterior-to-anterior). Given these altered starting conditions, the system behaves differently and patterning takes longer, but the same stable segmental pattern eventually emerges. Note that the primary pair-rule genes continuously oscillate within cells that do not express Opa, and segmental stripes emerge progressively from anterior to posterior as Opa turns on along the AP axis.

Inspection of the alternative network topology (Figure 13–figure supplement 1) reveals why inputs from Hairy alone are sufficient to organise the rest of the system. If Hairy is on, then *runt, ftz,* and *odd* will all be repressed, permitting the expression of *eve*. If Hairy then turns off, *runt* will be derepressed, soon resulting in the repression of *eve*. As with the “wild-type” network, this will then result in the derepression of *ftz/odd* and the consequent repression of *runt.* If Hairy then turned on once again, *ftz* and *odd* would be repressed, and the cell state would return to the beginning.

Thus, periodic bursts of Hairy expression are sufficient to organise repeating expression of the temporal sequence Hairy **⤑** Eve **⤑** Runt **⤑** Ftz/Odd within each nucleus. If appropriately spaced kinematic waves of Hairy expression travel from posterior to anterior across an array of cells, a correctly-phased pair-rule pattern will emerge, and downstream patterning will proceed as normal. Thus, only minor modifications to the *Drosophila* pair-rule network are required for it to acquire the capacity of autonomously carrying out pair-rule patterning, independent of continuing gap inputs.

#### Swapping temporal and spatial starting conditions transforms simultaneous patterning into a clock and wavefront-like system

The patterning potential of the alternative network topology described above demonstrates clearly that pair-rule patterning is a fundamentally temporal process, with appropriate spatiotemporal expression emerging from the internal dynamics of the regulatory system. During the “early” phase of the simulation, each nucleus cycles through the expression of the various primary pair-rule genes. Later, when Opa turns on and changes the regulatory logic, the expression within each nucleus stabilises, with the final segment-polarity state determined by the particular combination of pair-rule factors inherited from the dynamic, early phase. Cells further anterior or further posterior have reached different phases of the early pattern when the Opa signal is received, and therefore adopt different fates.

Thus, when simulating this modified version of the *Drosophila* network, cell fate determination results simply from the intersection of two temporal signals: 1) the periodic bursts of Hairy expression that organise primary pair-rule gene expression, and 2) a broad contextual signal, Opa, that influences the network as a whole. Positional information cannot be conveyed by either of these signals in isolation: for example, if Opa were to turn on slightly earlier or slightly later, the assignment of segment polarity fates to particular cells would be completely different.

In mimicking the patterning of the *Drosophila* blastoderm (Figure 13A; Movie 12), I have initialised the simulated tissue so that one of these two signals (Hairy) is spatially patterned, while the other (Opa) is not. However, this need not be the case. Because it is the combination of the two signals that specifies positional information, the particular distribution of spatial patterning between the two signals is theoretically irrelevant to the final pattern. If the tissue is instead initialised with ubiquitous Hairy expression, the loss of this spatial patterning can be compensated for by patterning signal X – which originally conveyed only strictly temporal information – with an anterior-to-posterior decay gradient. As X decays and disappears from progressively more posterior cells over time, Opa will turn on in an anterior-to-posterior wave.

Strikingly, but in line with the theory, a simulation set up this way yields the same final segment pattern as the previous simulation (Figure 13B; Movie 13). However, the spatiotemporal patterns of pair-rule gene expression observed over the course of the two simulations are very different. In the simulation where Hairy is spatially patterned (Figure 13A; Movie 12), pair-rule stripes emerge very early and all segments are patterned simultaneously. However, in the simulation where X is spatially patterned (Figure 13B; Movie 13), a regular pair-rule pattern does not emerge. Instead, the part of the tissue expressing X continuously undergoes synchronised expression oscillations, and segmental stripes emerge sequentially in an anterior-to-posterior sequence, as the anterior limit of X expression retracts across the tissue.

In other words, the first simulation resembles pair-rule gene expression during long-germ segmentation, while the second simulation resembles pair-rule gene expression during short-germ segmentation, simply as a result of altered starting conditions. The fact that a simple model based closely on the *Drosophila* pair-rule network is able to recapitulate both sets of segmentation dynamics strongly hints that the genetic networks operating in long-germ and short-germ arthropods may share much in common.

## DISCUSSION

### Pair-rule patterning is a fundamentally temporal process

In this manuscript I have analysed the structure and dynamics of the *Drosophila* pair-rule network, using a combination of simulated and experimental data to reveal how segment patterning is achieved. I have discovered a functional role for dynamic gap inputs, and propose revised mechanisms for the patterning of the odd-numbered and even-numbered parasegment boundaries. In contrast to previous models based around the principle of static morphogen gradients, these mechanisms involve a coordinated interplay between intrinsic network dynamics and extrinsic spatial and temporal signals, and do not necessarily require graded pair-rule activity.

These findings contribute to the evolving view of the role of Even-skipped, perhaps the best known of the pair-rule factors. Eve has long been known to be required for the expression of both sets of *en* stripes, and hence both sets of parasegment boundaries [112,113]. Originally, Eve was thought to achieve this directly, by activating *en.* Later, it was recognised that Eve does not regulate *en* directly, but instead represses several other pair-rule factors that themselves repress *en* [59,60]; however, quantitative information inherent within the Eve stripes was still believed to directly pattern the *en* stripes [36,61]. This conclusion is challenged by the new model of the pair-rule system presented here, which suggests that static domains of Eve expression would cause a similar degree of pattern loss to that seen in *eve* mutant embryos (compare Figure 5A and 12B).

Instead, I propose that the central role of Eve in pair-rule patterning results from a combination of the specific structure of the early pair-rule network and the specific dynamics of the posterior gap system. Of particular note, because the Eve stripes are dynamically expressed and Eve is also at the top of the pair-rule network hierarchy, the influence of Eve activity on gene expression is not restricted to the cells currently within each Eve stripe, but also extends to the cells posterior to the stripes, like wakes stretching out behind moving ships.

While this work clarifies many aspects of segment patterning, it also highlights certain temporal regulatory phenomena that appear to be crucial to the whole process, but which are currently much less well understood than the more overtly spatial aspects of gene regulation. Chief among these is the activation of the secondary pair-rule genes, *prd* and *slp,* at specific times during cellularisation. It seems likely that there are several temporal regulators of *Drosophila* segmentation that have not as yet been identified.

### Magnitudes of stripe shifts in the *Drosophila* blastoderm

I have presented evidence that posterior-to-anterior shifts of pair-rule stripes, driven by dynamic gap inputs, play a functional role in segment patterning. However, it is not straightforward to calculate the magnitude of the expression shifts exhibited by pair-rule stripes in real embryos. To date they have only been measured by comparing absolute stripe position (in % egg length) in fixed material of different ages [74,79]. These are of necessity population estimates, complicated by embryo-to-embryo variability. Interpreting these measurements is further complicated by the observation that nuclei migrate away from the poles of the embryo over the course of cycle 14, therefore raw measurements will overestimate shift magnitudes of posterior stripes. Allowing for this effect, published measurements suggests shifts of 2-3 nuclei for *eve* stripes 3-7, 1 nucleus for *eve* stripe 2, and 0 nuclei for *eve* stripe 1 [79].

These numbers suggest that the shifts in stripes 3-7 are large enough to support the dynamic patterning mechanisms proposed here, although it would be good to be able to confirm the measurements through live imaging of endogenous pair-rule gene expression using a method such as the MS2-MCP system [119–121]. Quantitative characterisation of stripe dynamics within individual embryos will help determine the relative contribution of shift-based and gradient-based mechanisms to segment patterning. Note, though, that these mechanisms are not mutually exclusive: indeed, the existence of shifts might well contribute to concentration-dependent spatial patterning of target genes, by helping shape the contours of the pair-rule stripes at the protein level.

The situation is less clear cut for the stripe 2 region (patterning for the stripe 1 region is already known to be different from the rest of the trunk, influenced strongly by anterior gap gene expression [122]). Any shifts that occur here are likely to be fairly insignificant; live imaging of an *eve* stripe 2 reporter element reveals no anterior shift at all [123], so the estimated 1 nucleus shift of the endogenous stripe 2, if accurate, would have to be mediated by the *eve* late element [23]. It may therefore be the case that independent patterning of the various pair-rule genes by stripe-specific elements has more importance in this anterior region of the embryo, with dynamic patterning taking over further posterior. This idea is supported by the fact that *prd* expression is independently regulated in this region (the early *prd* anterior domain gives rise to *prd* pair-rule stripes 1 and 2), rather than being patterned entirely by pair-rule inputs as in the rest of the trunk [26,111].

### *Drosophila* zebra elements: relictual components of an ancestral segmentation clock?

Recently, Michael Akam and I proposed that the topology of the pair-rule gene regulatory network is temporally regulated, with the early network involved in regularising the initial “pair-rule” patterns of gene expression during cellularisation, and the late network mediating the transition to patterns of single-segment periodicity during gastrulation [6]. In this paper, I have explored how the sparse early network relies on dynamic inputs from the gap system to correctly phase the pair-rule stripes, while the late network is independent of the gap system, and acts like a multi-stable switch between different segment-polarity fates.

I have also shown that it is fairly straightforward to convert the overall network into a “clock and wavefront” system, with the early network constituting the “clock” that generates periodicity, and the late network stabilising the outputs of the early network into segmental patterns (Figure 13B; Movie 13). Given that we know that long-germ insects such as *Drosophila* are derived from short-germ ancestors, it seems unlikely that this fluidity between simultaneous and sequential patterning modes is a mere coincidence. I therefore propose that output from the *Drosophila* early pair-rule network is developmentally homologous to the oscillatory pair-rule gene expression seen in the growth zone of short-germ insects such as *Tribolium castaneum,* while the output from the late pair-rule network is developmentally homologous with the refining stripes seen anterior to the growth zone (Figure 14).

**Figure 14:**
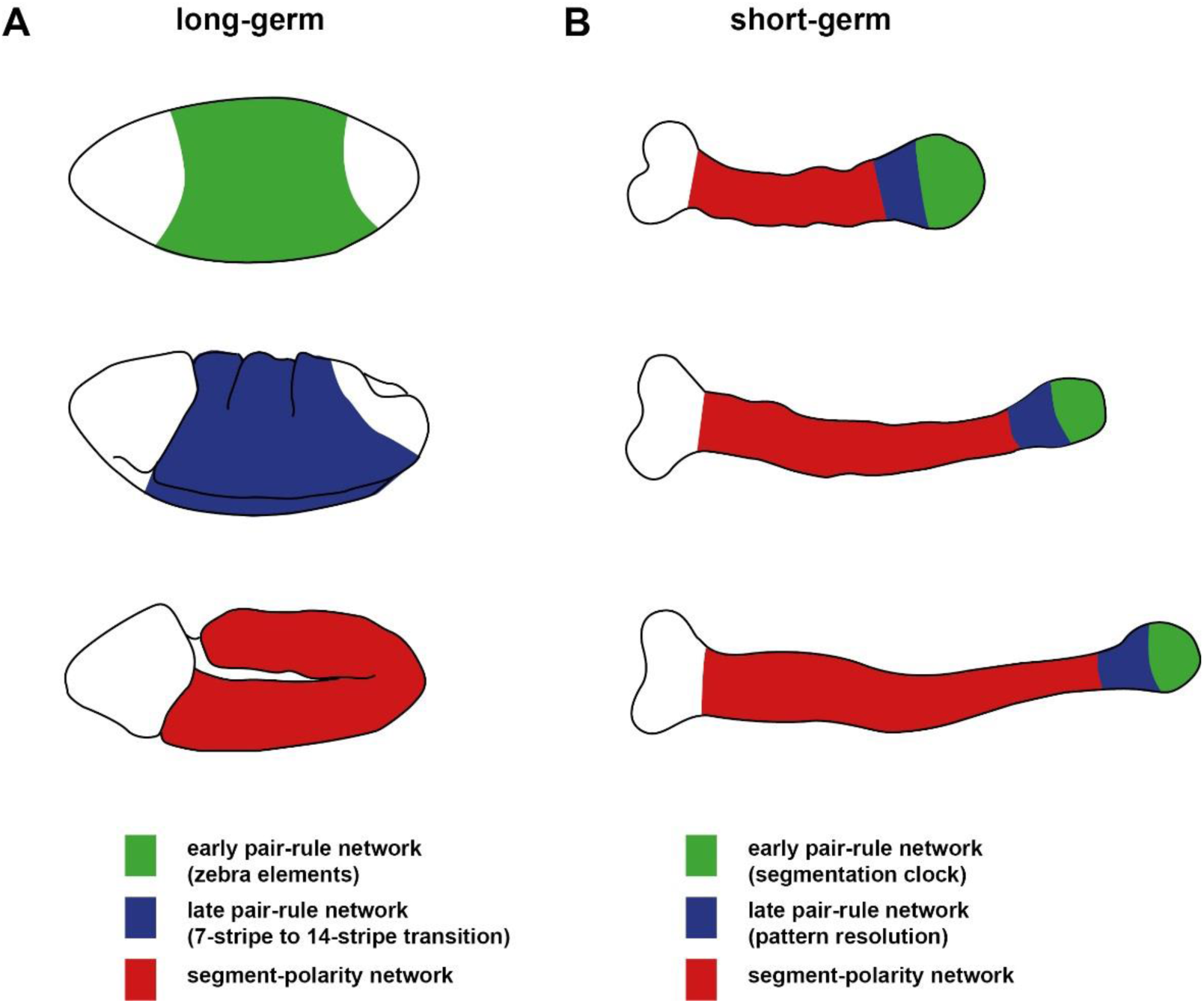
proposed regulatory homology between phases of segmentation gene expression in long-germ and short-germ embryos. **A**: In *Drosophila,* the quintessential long-germ insect, there are three main phases of segment patterning during embryogenesis. During cellularisation (top), most pair-rule genes are regulated via zebra elements, and become expressed in periodic patterns throughout the trunk of the embryo (green region). During gastrulation (middle), segmental patterns of pair-rule genes and segment-polarity genes emerge throughout the trunk (blue region) fairly simultaneously. Finally, during germband extension (bottom), the segment-polarity network maintains the segment pattern via intercellular signalling throughout the trunk (red region). **B**: In short-germ embryos, for example those of the red flour beetle, *Tribolium castaneum,* segment patterning occurs continuously throughout germband extension. Three germbands of increasing age are depicted: the trunk elongates from the posterior as embryogenesis proceeds. Throughout the whole process, primary pair-rule genes exhibit oscillatory expression in the posterior growth zone (green regions). I propose that this phase of expression is developmentally homologous to the pair-rule gene expression regulated by the early network in *Drosphila* (i.e. zebra elements may be derived from clock enhancers). Pair-rule stripes then resolve and (may undergo frequency doubling) in the anterior growth zone (blue region). I propose that this phase of expression is developmentally homologous to the pair-rule gene expression regulated by the late network in *Drosphila*. Finally, segment-polarity genes are expressed in segmental stripes starting just anterior to the growth zone (red regions). This phase of expression is thought to be regulated by a conserved segment-polarity network [141,142].

Aside from the motivating evidence from the dynamics of the segmentation process, this hypothesis is consistent with several other lines of comparative evidence gathered from various arthropod model species. For example, it fits with the strikingly conserved temporal sequence of pair-rule gene expression observed from both short germ and long-germ embryos [68]. It is also supported by the deep similarity between the morphogenetic processes involved in *Drosophila* germband extension and those driving convergent extension anterior to the *Tribolium* growth zone [124,125].

Note, however, that the “clock and wavefront” system that emerges from the modified pair-rule network (Figure 13B; Movie 13) is reliant on the regular, deterministic framework of the simulation, whereas the segmentation mechanisms operating in real short-germ embryos presumably involve additional elements. For example, synchronising intercellular interactions would be required to counteract disruptive effects such as transcriptional noise and cell rearrangements. (These signals could simply feed into the intracellular transcriptional network of the basic system, perhaps by influencing *hairy* expression, as seen in the cockroach *Periplaneta americana* [126].) In addition, in order to recover the slowing, narrowing kinematic waves of pair-rule gene expression seen in real embryos, the unrealistic instantaneous transition from oscillatory to stabilising regimes used in the model could be replaced by a frequency gradient of some sort [127].

### The evolution of long-germ segmentation

If true, the hypothesis proposed above suggests that the evolutionary transition from sequential to simultaneous patterning would require two important changes, but that the bulk of the pair-rule network could persist unaltered.

The first change is that the regulation of *hairy* and *eve*, which is presumably governed by “clock enhancers” in short-germ species, would have to be taken over by gap-regulated stripe-specific elements. However, in order for these gap-regulated stripes to maintain the same patterning output from the pair-rule network as a whole, they would have to be expressed with the same dynamics as if they were produced by a segmentation clock, thus strongly constraining the dynamics of the gap system. This prediction is strikingly concordant with recent findings that the *Drosophila* gap system exhibits oscillatory dynamics in the posterior half of the blastoderm [46].

The second change is that the spatiotemporal patterning of extrinsic signals (such as Opa) that control the timing of segmentation would have to be modified from some kind of wavefront-like expression coordinated with posterior elongation, to simultaneous expression within the blastoderm. This would allow pattern resolution to occur early and all at once, rather than gradually over the course of germband extension.

While gap-based patterning of individual pair-rule stripes is much more complicated than the algorithmic generation of periodicity by a segmentation clock, there is an obvious payoff in terms of the speed of development. A segmentation clock can only tick so fast, and therefore represents a tight production bottleneck when generating segments (Figure 14–figure supplement 1). (Indeed, selection for quicker development may underlie the evolutionary success of pair-rule patterning, which effectively halves the number of clock cycles required to produce a body of a given length.) In contrast, as demonstrated by the *Drosophila* embryo, segmentation can be accomplished remarkably quickly when a periodic spatial pattern is present from the beginning.

Still, it has been hard to imagine how pair-rule gene expression could easily transition from being controlled by a segmentation clock to being controlled by gap inputs, while all the while producing a strongly conserved segment polarity pattern. The proposal a little over a decade ago that the takeover by the gap system occurred progressively along the AP axis [128] mitigates this problem significantly. Complementing this idea, the findings in this paper suggest additional ways in which this transition could occur gradually and seamlessly.

First, because it seems likely that the role of the gap system is effectively to mimic the output and dynamics of a segmentation clock, there is no reason why gap-based patterning and clock-based patterning shouldn’t work partially redundantly to pattern a given set of stripes during the transition from one mechanism to the other. Indeed, it is possible that the two mechanisms coexist today in insects such as *Nasonia* and *Bombyx,* which exhibit some evidence of both patterning modes [129–131].

Second, because the early pair-rule network has the capacity to organise the expression of the whole system given minimal inputs, gap control of all the pair-rule genes would not have to evolve all at once. Gap control of just one pair-rule gene (e.g. *hairy)* could theoretically be sufficient to produce the correct final output (Figure 13A), while stripe-specific elements for the remaining pair-rule genes could then evolve later, offering additional boosts to patterning speed and reducing the magnitude of required gap gene shifts (Figure 14–figure supplement 1). Indeed, a recent *in silico* evolution study [132] concluded that it would be difficult to significantly alter the gap system without disturbing the overall pattern of pair-rule gene expression from the stripe-specific elements, and so made the hypothetical suggestion that Eve stripe shifts combined with appropriate pair-rule cross-regulation might offer a solution to this problem – dovetailing nicely with the findings in this study.

It is also possible that sufficiently robust gap patterning of pair-rule stripes could supplant the need for expression shifts entirely: this might well apply to the *Drosophila* stripe 2 region (see above), in which case it offers a downstream functional reason for why the transition between static and dynamic regimes of gap gene expression occurs further posterior in *Drosophila* than in other dipterans such as the scuttle fly *Megaselia abdita* [133,134].

## CONCLUDING REMARKS

In the introduction to this paper, I highlighted three recent findings related to pair-rule patterning: 1) that pair-rule orthologs exhibit oscillating expression in the growth zones of short-germ arthropods, 2) that pair-rule gene expression in long-germ insects is patterned by dynamic gap gene expression, and 3) that the pair-rule network in *Drosophila* undergoes an extensive topology change at gastrulation. I then presented evidence that dynamic Eve expression is crucial for segment patterning, and also that the *Drosophila* pair-rule network is broadly compatible with both simultaneous and sequential modes of segmentation.

I therefore propose the following evolutionary hypothesis, which ties together and potentially explains the three original findings. 1) Long-germ and short-germ segmentation are not dichotomous modes of patterning, but rather alternative behaviours of a largely conserved pair-rule network. 2) In long-germ insects, *ad hoc* patterning by gap inputs stands in for the expression of particular pair-rule “clock enhancers”, meaning that gap gene networks are under strong selection to preserve the original oscillatory expression dynamics of the pair-rule genes. 3) In *Drosophila,* the “early” pair-rule network is derived from the “clock” part of an ancestral short-germ segmentation mechanism, while the “late” network is derived from the stabilising genetic interactions that would have originally established parasegment boundaries anterior to the growth zone.

These predictions can be tested in the future by comparative studies in emerging arthropod model species.

## MATERIALS AND METHODS

Wild-type in situ images are from a previously published dataset [135]. The *eve* mutation used was *eve^3^* (gift of Bénédicte Sanson) and was balanced over *CyO hb::lacZ* (Boomington stock no. 6650) in order to easily distinguish homozygous mutant embryos. Whole mount double fluorescent in situ hybridisation, microscopy, and image analysis were carried out as described previously [6]. The intensity profiles in Figure 10C were produced using the “multi plot” function in Fiji [136]. Simulations were coded in Python (www.python.org), using the libraries NumPy [137] and Matplotlib [138]. All models and simulations are described in detail in Supplement 2.

## ACKNOWLEDGEMENTS

I am very grateful to Michael Akam, who advised and encouraged me throughout the course of this project and edited the manuscript, and to Tim Weil, who contributed lab space and taught me to work with flies. Michalis Averof and Ezzat El-Sherif commented on a draft of the manuscript. I thank Andrew Peel, Andy Oates, Johannes Jaeger, Berta Verd, Siegfried Roth, Bénédicte Sanson, James Briscoe, and Alfonso Martinez-Arias for useful discussions.

## FUNDING

This project was funded by a PhD studentship from the Biotechnology and Biological Sciences Research Council “Genes to Organisms” doctoral training program, and a research grant from the Isaac Newton Trust.

## COMPETING INTERESTS STATEMENT

I declare that no competing interests exist.

## SUPPLEMENTARY MATERIAL

### Appendices

Appenidx 1: Genetic interactions between the primary pair-rule genes during cellularisation

Appendix 2: Details of models and simulations

### Supplementary files

Supplementary File 1: Simulation software

Supplementary File 2: Example simulation script

### List of Movie files

**Movie 1:** Simulated expression of the primary pair-rule genes, assuming static gap inputs

**Movie 2:** Simulated expression of the primary pair-rule genes, assuming dynamic gap inputs

**Movie 3:** Simulated expression of the primary pair-rule genes, assuming slow gap shifts

**Movie 4:** Simulated expression of the primary pair-rule genes, assuming fast gap shifts

**Movie 5:** Simulated expression of the primary pair-rule genes, assuming very fast gap shifts

**Movie 6:** Pattern establishment when *hairy, eve,* and *runt* take gap inputs

**Movie 7:** Pattern establishment when *hairy, eve,* and *ftz/odd* take gap inputs

**Movie 8:** Pattern establishment when all primary pair-rule genes take gap inputs

**Movie 9:** Simulated behaviour of the whole pair-rule system, assuming static gap inputs

**Movie 10:** Simulated behaviour of the whole pair-rule system, assuming dynamic gap inputs

**Movie 11:** Pair-rule gene expression in a simulated *eve* mutant **Movie 12:** Simultaneous patterning by the modified pair-rule network

**Movie 13:** Sequential patterning by the modified pair-rule network

Each simulation has two alternate versions, **A** and **B**. The former are straightforward visualisations of expression state at each time point: pale colours represent active transcription, and dark colours represent protein activity, consistent with the figures in the manuscript. The alternate versions additionally visualise time delay information, giving a greater insight into the logic of the simulations. Solid colours represent protein activity, while hashed colours represent active transcription. Time delays are conveyed by a transparency effect: after a gene turns on in a given cell, the transcriptional output will gradually gain intensity until the protein turns on, while after a gene turns off in a given cell, the protein output will gradually lose intensity until the protein activity turns off. (Note that although this effect can give the impression of changing gene product concentrations, the logic of the system remains entirely Boolean.)

---

**Figure 4-figure supplement 1:**
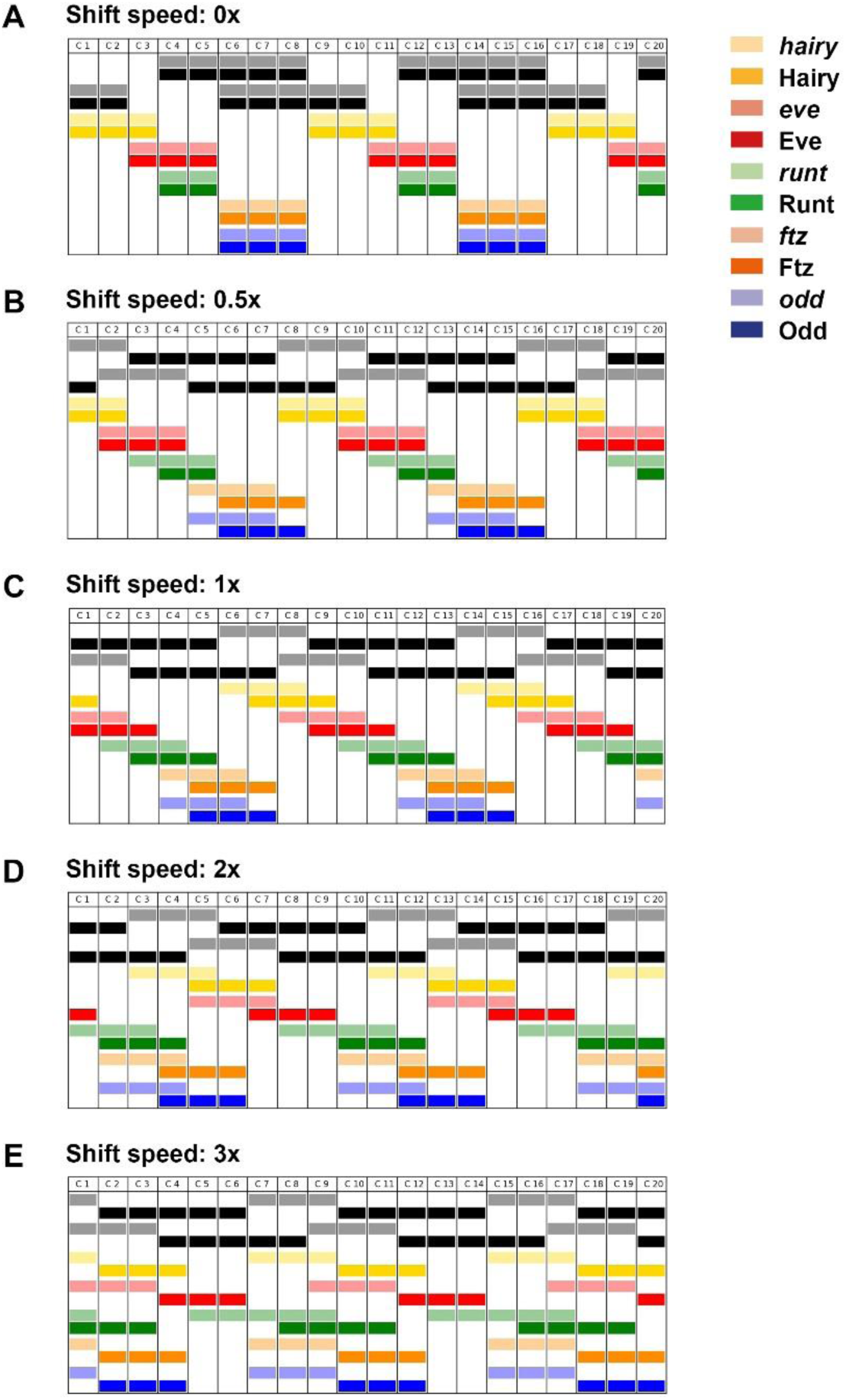
Stripe phasing depends on the speed of the gap shifts. Simulation output showing the expression patterns generated by the early pair-rule network, assuming various anterior-to-posterior shifts speeds of the gap inputs (shown in black). See Movies 1-5 for full simulation output (Movie 1 = 0x; Movie 2 = 1x; Movie 3 = 0.5x; Movie 4 = 2x; Movie 5 = 3x). Gap domain shift speeds are relative to the time delay for pair-rule protein synthesis / decay: a speed of 1x leads to a one nucleus offset between the anterior borders of transcript and protein domains, a speed of 2x leads to a two nucleus offset, and so on. See Supplement 2 for further details about the simulations.

**Figure 6-figure supplement 1:**
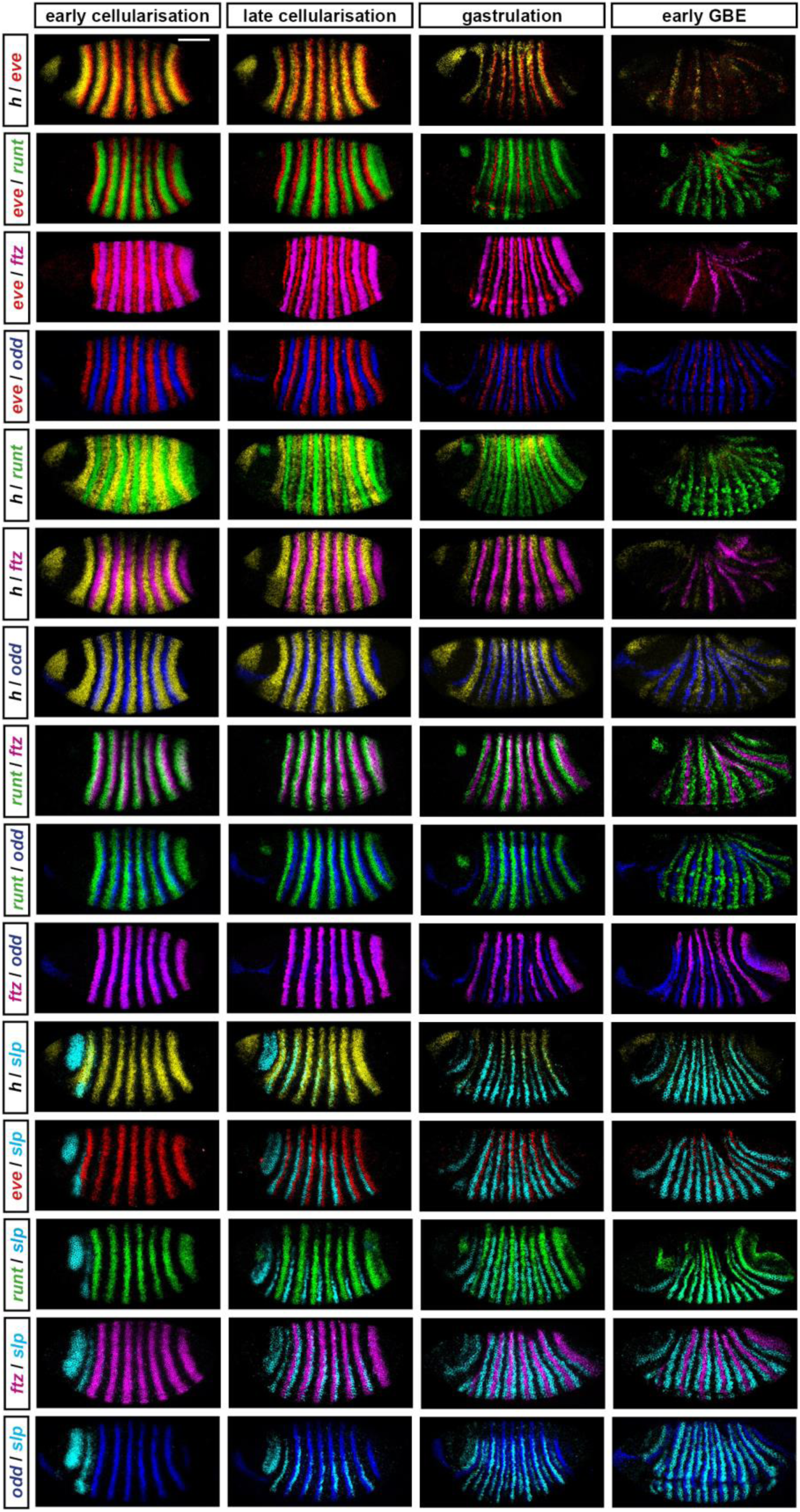
Uncropped views of the embryos shown in Figure 6. Each panel shows an uncropped version of the corresponding double fluorescent *in situ* image in Figure 6. All panels show a whole embryo lateral view, anterior left, dorsal top. Embryos within each column are of approximately equal age. Scale bar = 100 μm.

**Figure 6-figure supplement 2:**
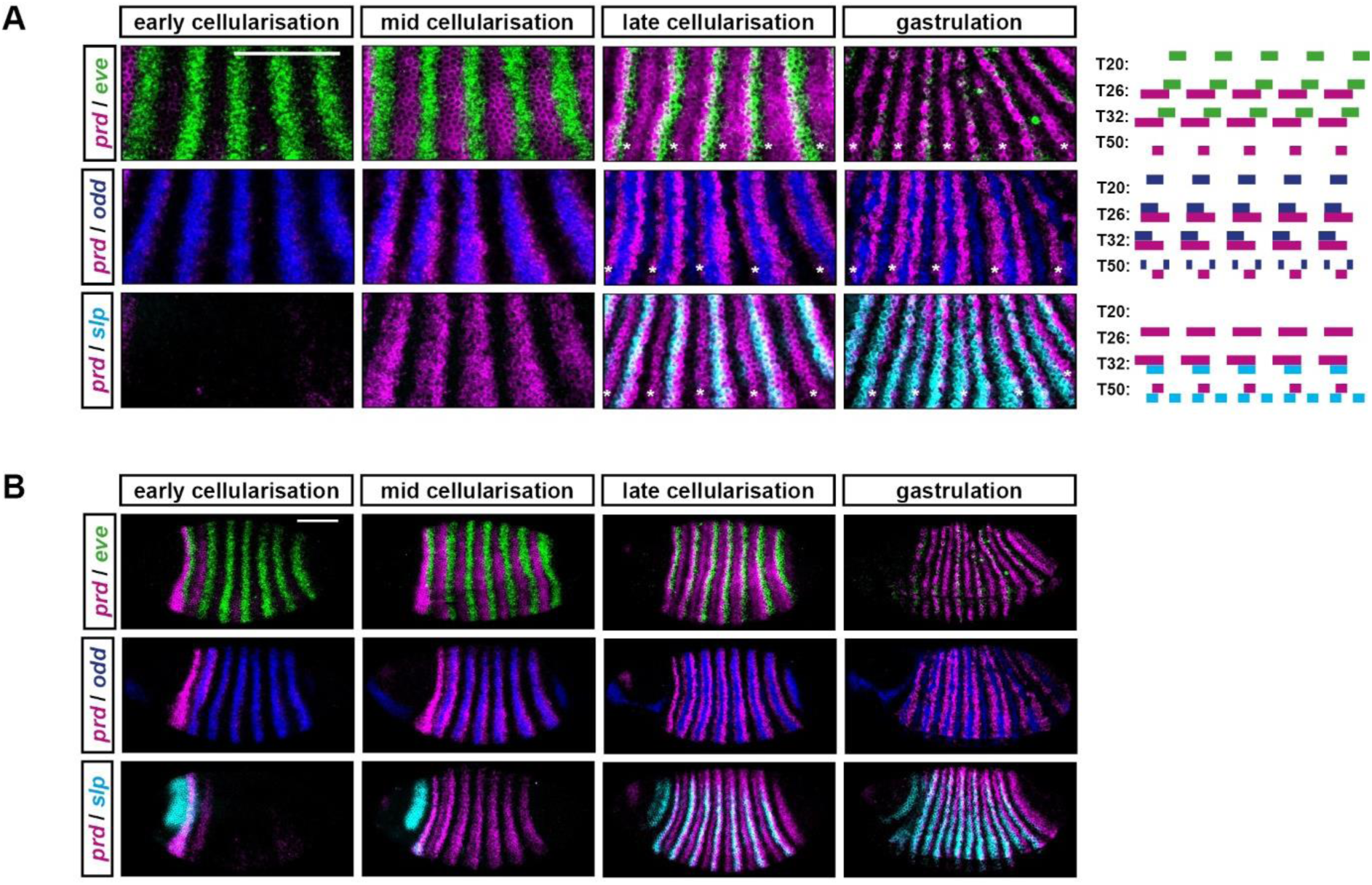
Some aspects of *prd* expression are not recovered by the simulation. This figure compares double fluorescent *in situ* data of pair-rule gene expression with simulated transcript expression, as in Figure 6. **A**: *prd* transcripts (magenta) are shown relative to *eve* (green), *odd* (blue) or *slp* (cyan) transcripts, at four different embryo ages (left) or simulation timepoints (right). Note that the figure presents a different set of developmental ages / simulated timepoints from those in Figure 6, in order to show cellularisation in greater temporal detail. At “early cellularisation”, the primary pair-rule genes are expressed but the secondary pair-rule genes *prd* and *slp* are not; at “mid cellularisation”, *prd* is expressed but *slp* is not; at “late cellularisation”, *slp* is additionally expressed. While the simulated expression of *prd* is broadly appropriate early on (e.g. compare *prd* and *odd* expression at mid-cellularisation / T26), several aspects of *prd* expression are not recapitulated by the model. First, the changing phasing of *prd* posterior borders and *eve* anterior borders between mid-cellularisation and late cellularisation (see also Figure 8). Second, the splitting of the *prd* stripes at late cellularisation, slightly prior to the appearance of the secondary stripes of *odd* and *slp* [6]. Third, the patterning of the *prd* “A” stripes (i.e. the narrow stripes formed from anterior portions of the early broad stripes, asterisks in the *in situ* images) and the consequent emergence of single-segment periodicity. **B**: Uncropped views of the embryos shown in **A**. Scale bars 100 μm.

**Figure 7–supplementary figure 1:**
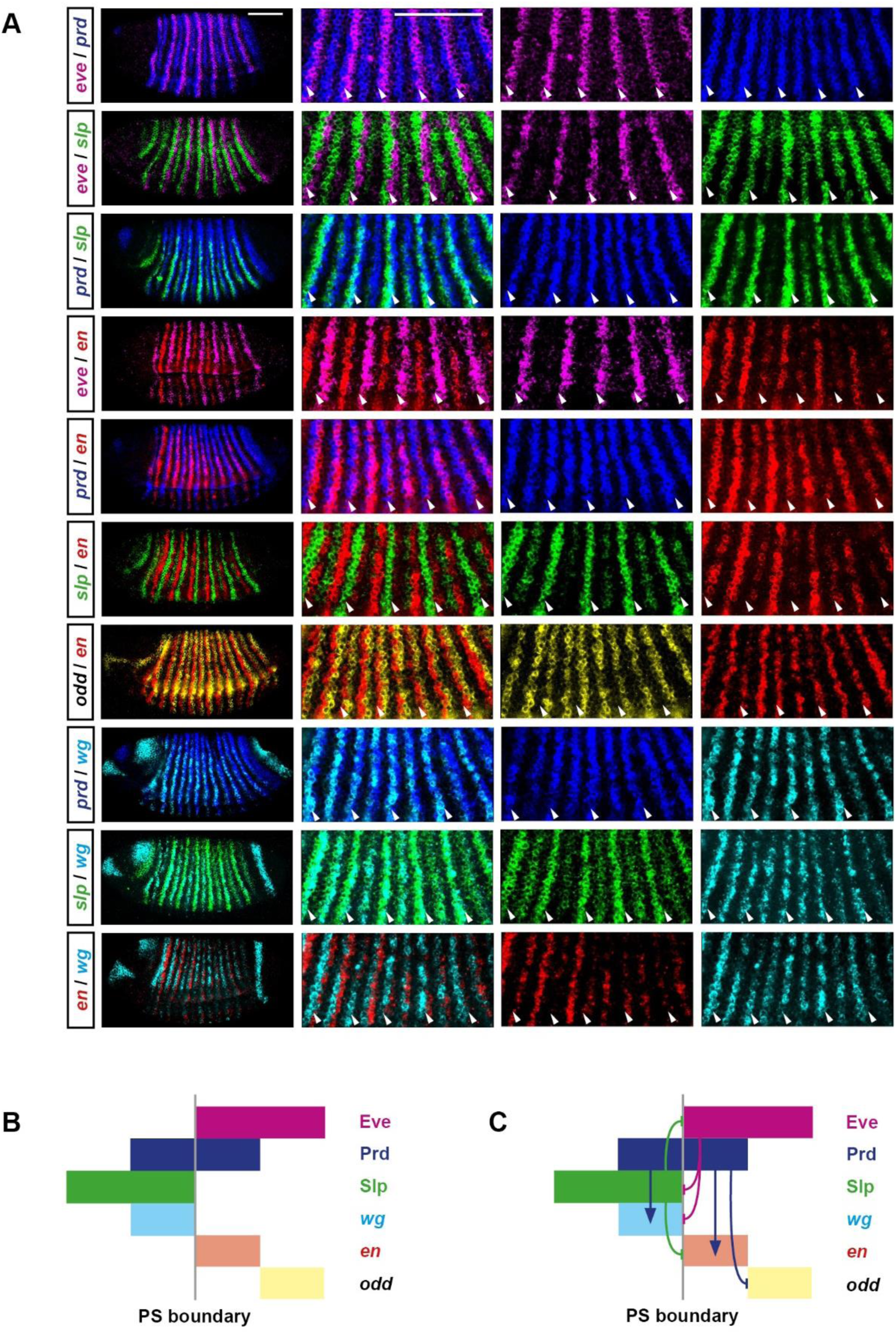
Gene expression at the odd-numbered parasegment boundaries. **A**: Double fluorescent *in situ* images of segmentation gene expression in gastrulation stage embryos (lateral view, anterior left, dorsal top). Each row shows a different pairwise comparison of the transcripts of the genes *eve* (magenta), *prd* (blue), *slp* (green), *en* (red), *odd* (yellow), and *wg* (cyan). From left to right, the panels in each row show: 1) a whole embryo view; 2) an enlarged view of pair-rule repeats 2-6; 3) the individual channel for the first gene listed in the row label; 4) the individual channel for the second gene listed in the row label. Arrowheads mark the locations of prospective odd-numbered parasegment boundaries. Scale bars 100 μm. **B**: Summary schematic showing the relative expression of the Eve, Prd, and Slp protein domains, and the *en, wg,* and *odd* transcript domains, at the odd-numbered parasegment boundaries. **C**: The same schematic as in **B**, with the addition of the regulatory interactions between Eve, Prd, and Slp and their target genes. Aside from the colours, which are changed to match the *in situ* images, this schematic is identical to Figure 7A.

**Figure 13-figure supplement 1:**
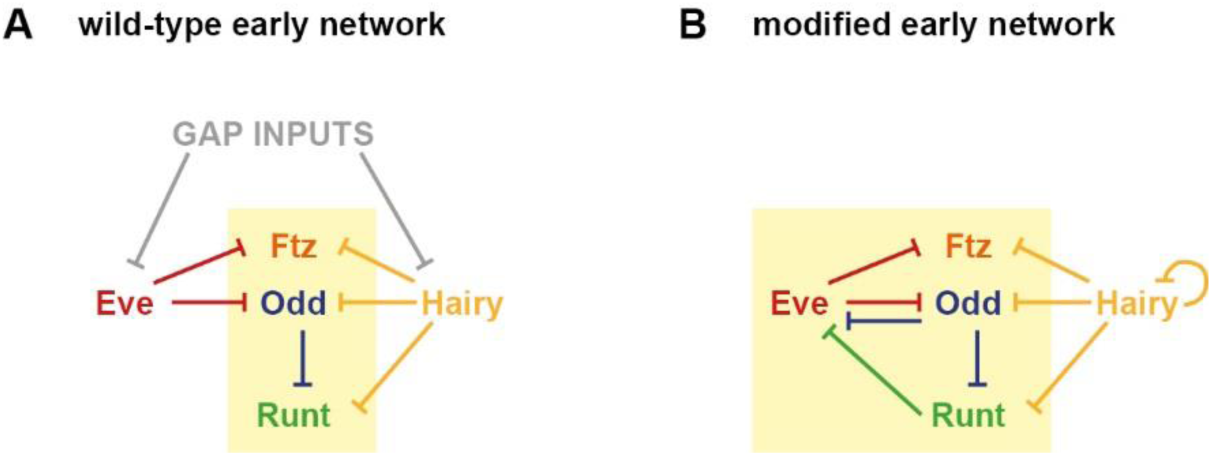
A modified early network in which gap inputs are replaced by cross-regulatory interactions. **A**: The wild-type “early” network (only the primary pair-rule genes are shown, for simplicity). **B**: A modified early network, in which the regulation of Eve and Hairy by gap inputs is replaced by additional cross-regulatory interactions. Eve is repressed by Runt and Odd, while Hairy is repressed by itself. This hypothetical network topology is used for the simulations in Figure 13.

**Figure 14–figure supplement 1:**
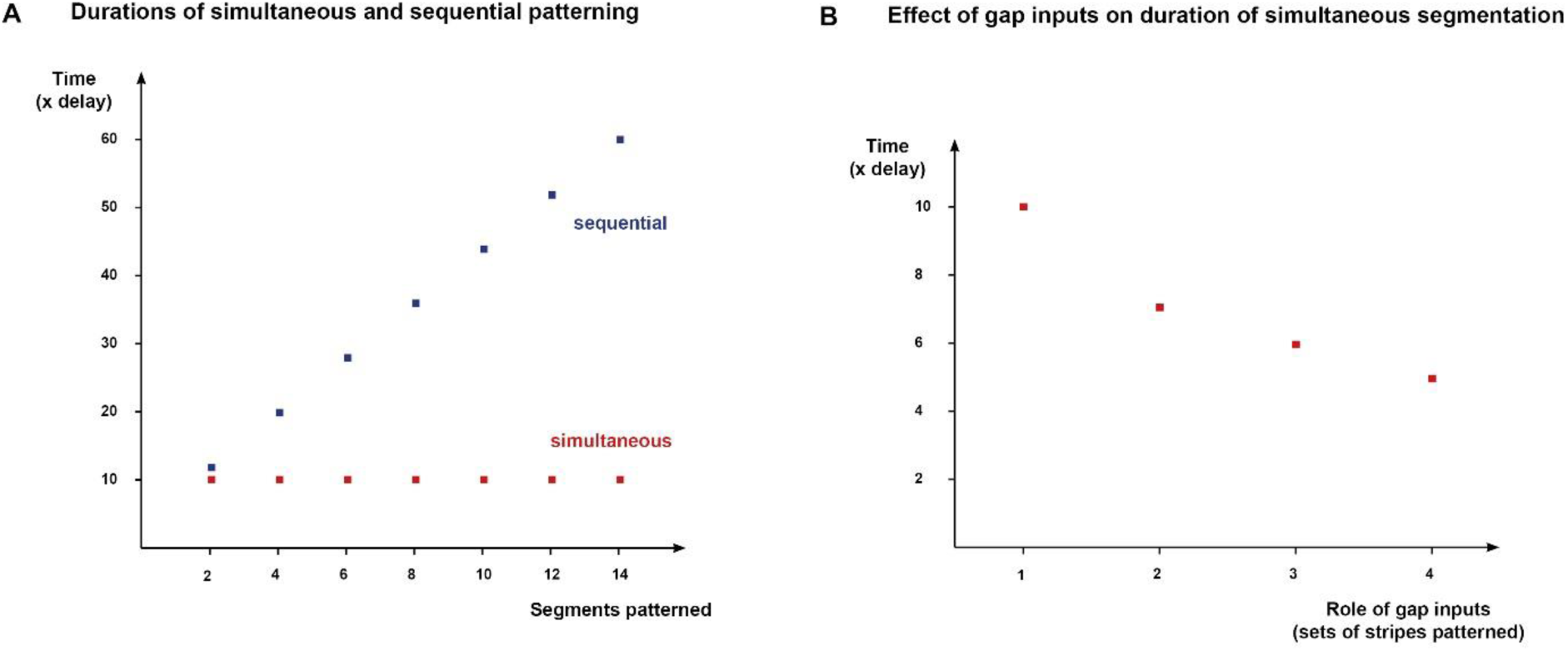
Selection for patterning speed could lead to increasingly elaborate regulation of pair-rule genes by the gap system. **A**: Time required to pattern a given number of segments by either simultaneous or sequential patterning. Patterning duration is given as a multiple of the synthesis / decay time delay of the pair-rule gene products. Times shown are minimums for the modified pair-rule network (Figure 13). Sequential patterning requires 12x the delay for the initial two segments, plus 8x the delay for every two additional segments. Simultaneous patterning requires a constant minimum time of 10x the delay, no matter the number of segments. **B**: Minimum time (in multiples of the synthesis / decay delay) required for simultaneous patterning, for systems with increasingly elaborate gap patterning. If only *hairy* receives gap inputs (1), a minimum of time 10x the delay is required. If both *hairy* and *eve* receive gap inputs (2), the minimum is 7x the delay. If *runt* or *ftz/odd* additionally receive gap inputs (3), the minimum is 6x the delay. Finally, if all the primary pair-rule genes receive gap inputs (4), the minimum is 5x the delay. Note that the stripes of *ftz* and *odd* are considered a single pattern, because of their identical regulation in the simulations.

## Appendix 1

### Regulatory interactions between the primary pair-rule genes during cellularisation

This appendix concerns the cross-regulatory interactions between the five primary pair-rule genes *(hairy, eve, runt, ftz,* and *odd)* during cellularisation. Looking at each gene in turn, I examine the evidence for its expression being directly regulated by the other primary pair-rule factors. The conclusions form the basis for the topology of the “early” pair-rule network presented in Figure 1A.

Gene regulatory network models have been characterised as “intellectual syntheses” of the combined evidence (typically expression data) from a large number of diverse experiments [143]. In order to analyse the control logic of the *Drosophila* primary pair-rule genes, the main sources of evidence I consider are wild-type stripe phasings, expression patterns in mutant and transgenic embryos, and regulatory element reporter studies. I have collated relevant observations from the literature, and complement these with new double fluorescent *in situs* from *hairy, eve* and *runt* mutant embryos.

An important aspect of this analysis is determining the timing of particular expression changes, in order to disentangle regulatory interactions that form part of the early network from those that only relate to the late network. Based on this analysis, I conclude that Hairy and Eve act largely as “input-only” factors in the early pair-rule network, and organise the expression of the remaining pair-rule genes around themselves.

Note that figures in this Appendix may refer to embryos as being at “phase 1”, “phase 2”, or “phase 3”. As defined in Clark & Akam 2016b, “phase 1” refers to early cellularisation, when most pair-rule gene expression is controlled in an ad hoc manner by stripe-specific elements, “phase 2” refers to mid-cellularisation, characterised by regular periodic patterns usually driven by zebra elements, and “phase 3” refers to late cellularisation and gastrulation, when the transition to single segment periodicity occurs. The early network operates during phase 2, while the late network operates during phase 3.

#### Regulation of *hairy*

##### Regulatory elements

*hairy* possesses a full set of stripe-specific elements (1+5, 2+6, 3+4, 7, reviewed in Schroeder et al. 2011). However, *hairy* is not known to possess any kind of periodically expressed element. This suggests that the majority of its regulation comes directly from the gap system.

##### No evidence for regulation by Eve, Ftz, or Odd

In wild-type embryos, expression of *hairy* overlaps with Eve, Ftz and Odd (inferred from Pisarev et al. 2009 and Clark & Akam 2016a), indicating that it is not repressed by any of them. In agreement with this, the *hairy* expression pattern is not significantly altered by ectopic expression of Eve, Ftz or Odd, aside from repression of *hairy* stripe 1 in HS-Odd embryos [59,86,145]. *hairy* expression is also largely normal in *ftz* and *odd* mutant embryos [26,30,145].

In contrast, *hairy* expression is rather abnormal in *eve* mutant embryos: *hairy* stripe 2 becomes repressed, while the remaining stripes exhibit abnormal widths and spacings (Ingham & Gergen 1988; Hooper et al. 1989; Vavra & Carroll 1989; Appendix 1-figure 1). However, these changes are unlikely to reflect direct regulation of *hairy* by Eve. *hairy* stripe 2, which is sensitive to Slp, is presumably repressed by the ectopic Slp expression that occurs in *eve* mutant embryos [26,122]. The subtler effects on the remaining stripes are as yet unexplained, but it has been suggested that they reflect a patterning role of the early broad expression of Eve during cycles 12 and 13 [32,34]. If so, the effects on the *hairy* stripes are likely indirect, and mediated via subtle changes to gap gene expression.

**Appendix 1–figure 1:**
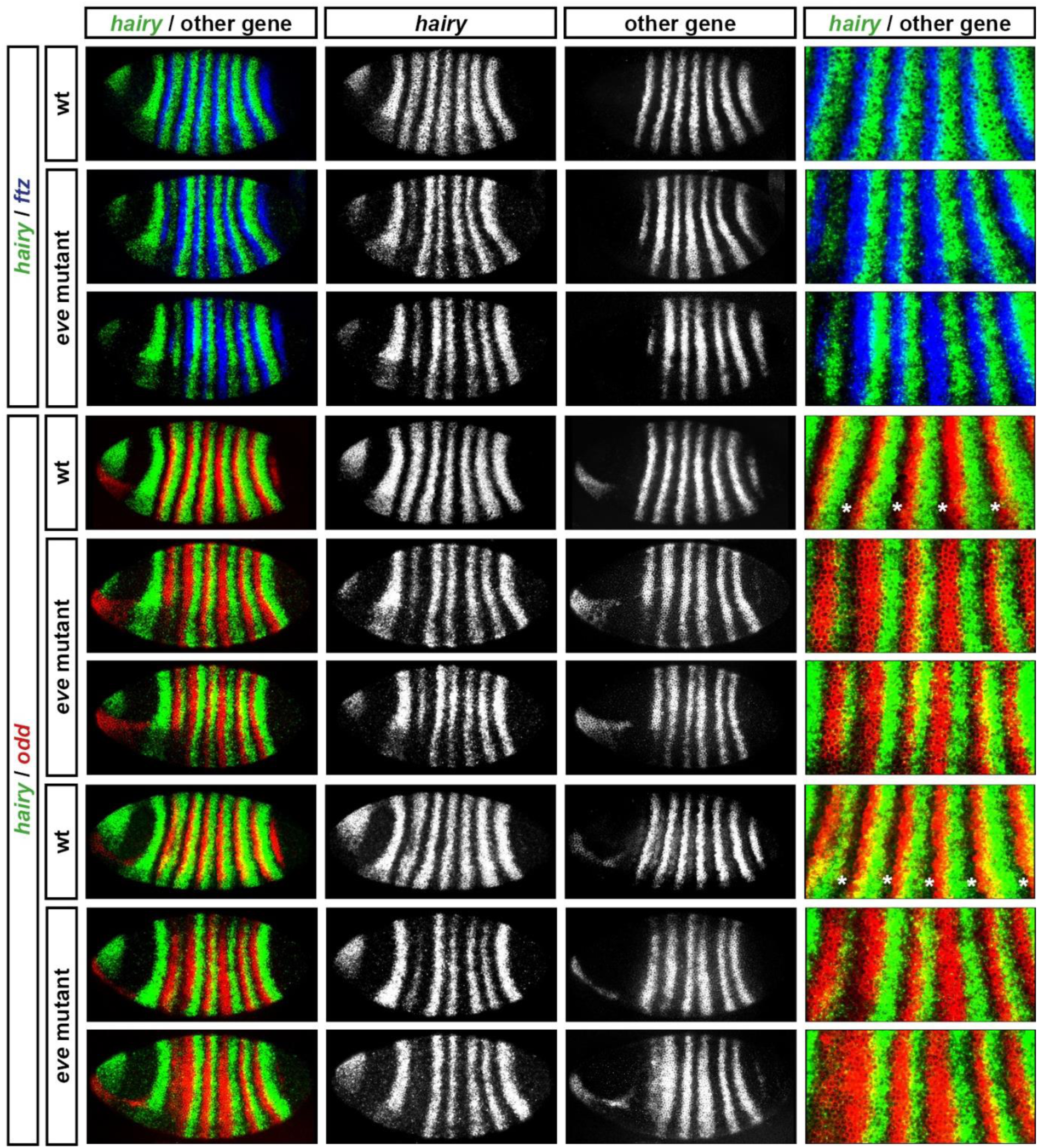
*ftz* and *odd* are patterned by Hairy in *eve* mutant embryos. Expression of *ftz* and *odd* relative to *hairy* in wild-type and *eve* mutant embryos. For the *hairy/odd in situs,* the upper three panels show embryos at mid cellularisation while the lower three panels show embryos at late cellularisation. Two different mutant embryos are shown for each time point. Note loss of *hairy* stripe 2 in *eve* mutant embryos, and corresponding anterior expansion of *odd* stripe 2. Note also the broadened stripes 2 and 4 of both *ftz* and *odd,* and the reduction of the clear gaps between the posteriors of the *hairy* stripes and the anteriors of the *odd* stripes (asterisks in wild-type embryos). In addition to the repression of *hairy* stripe 2, *hairy* stripes 3-6 exhibit abnormal widths and spacing.

##### Little evidence for regulation by Runt

It has been previously proposed that *hairy* is directly repressed by Runt. In wild-type embryos the *hairy* stripes are out of phase with the *runt* stripes (Figure 2E), and *runt* mutant embryos exhibit ectopic expression of *hairy* [33,34,146,147]. Direct repression by Runt is thought to cause the splitting of the *hairy* stripe 3+4 element into distinct stripes [148,149], and is also thought to be involved in proper separation of *hairy* stripes 6 and 7.

However, this evidence is not clear cut. Runt cannot be absolutely required for the splitting of *hairy* 3+4, as this splitting still occurs, at least temporarily, in *runt* mutant embryos (Vavra & Carroll 1989; Appendix 1–figure 2). In addition, *hairy* expression is not significantly affected by heatshock-mediated misexpression of Runt in blastoderm stage embryos [60,150]. At 30 minutes after heatshock (when direct effects would be expected to be evident), only *hairy* stripe 1 is repressed by ectopic Runt. Later weakening of stripes 2, 5, and 6 in these experiments could be indirect effects, perhaps via documented effects of Runt on gap gene expression [150,151]. Notably, *hairy* stripes 3 and 4 do not appear at all repressed in the HS-Runt embryos, indicating either that the splitting of stripes 3 and 4 in wild-type embryos is not mediated via direct repression of the 3+4 element by Runt, or that this element is sensitive to Runt only temporarily.

**Appendix 1–figure 2:**
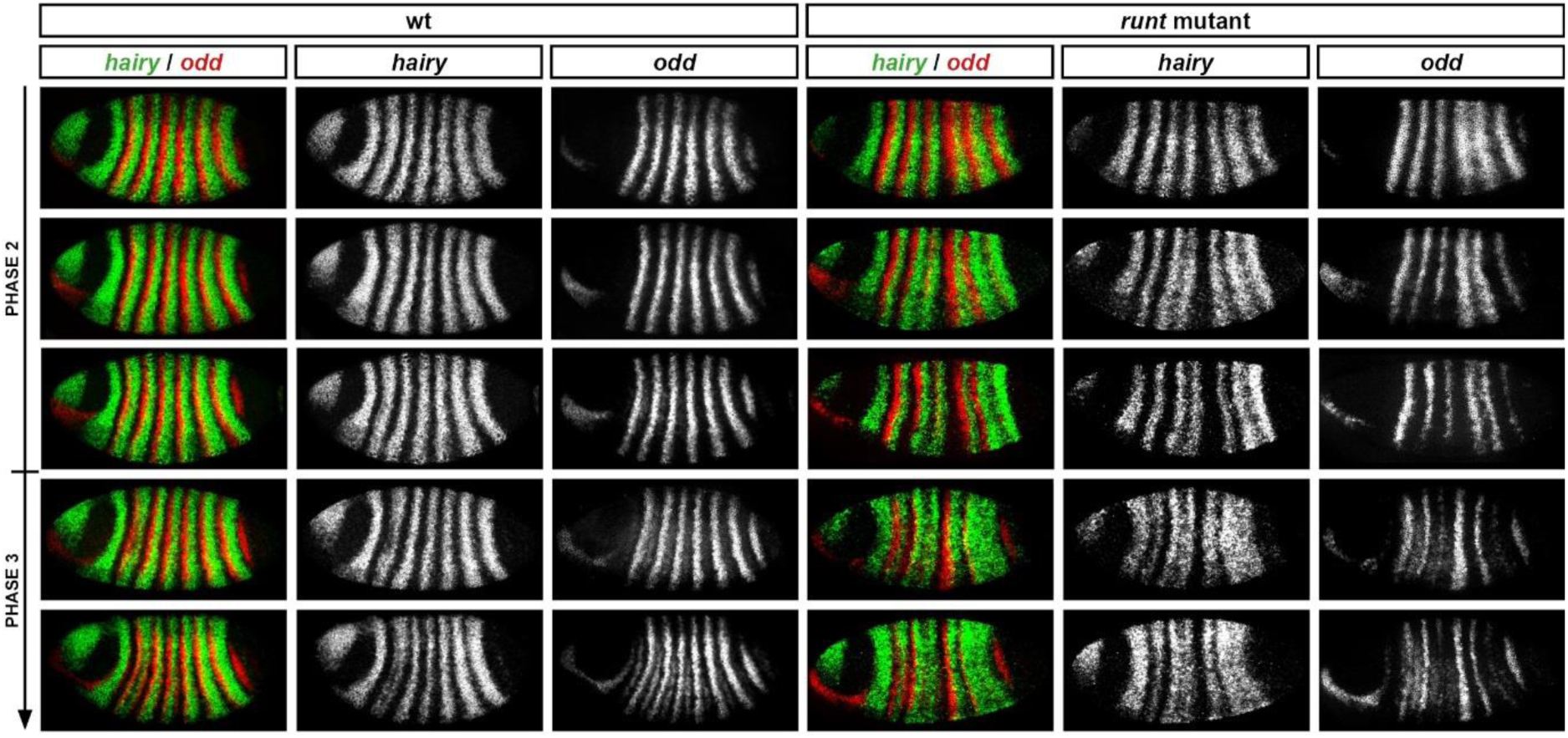
*hairy* stripes do not fuse until late cellularisation in *runt* mutant embryos. Relative expression of *hairy* and *odd* in wild-type and *runt* mutant embryos. The *hairy* stripes establish fairly normally (row 1), but the gap between stripes 4 and 5 widens during mid-cellularisation (rows 2-3), then fusions between stripes 3-4 and 6-7 occur at late cellularisation (rows 4 and 5). *odd* expression correlates negatively with *hairy* expression at all stages. Arrow indicates increasing developmental age. Single channel images are shown in greyscale to the right of the double channel images.

The *hairy* stripes seem to establish fairly normally in *runt* mutant embryos, with the fusions of stripes 3/4 and stripes 6/7 not occurring until late cellularisation (Appendix 1–figure 2). This casts further doubt on a direct role for Runt in specifying the *hairy* pair-rule pattern. First, spatial inputs from Runt are not required for the initial emergence of seven *hairy* stripes (cf. Hartmann et al. 1994). Second, the late appearance of ectopic *hairy* expression indicates that any direct regulation of *hairy* by Runt may be specific to the late network. Suggestively, similar fusions of *hairy* stripes 3/4 and 6/7 occur at gastrulation in *opa* mutant embryos [6], indicating that Runt and Opa might cooperate to repress Hairy, in the same way that they cooperate to repress Odd.

##### Conclusion

In summary, there is little convincing evidence for direct regulation of *hairy* by other primary pair-rule factors during cellularisation. Eve and Runt activity certainly influence *hairy* expression, but their effects seem to be either indirect, or restricted to later phases of patterning. Clear evidence of direct repression during cellularisation is limited to specific anterior stripes (e.g. stripe 1 is sensitive to Runt and Odd, and stripe 2 is sensitive to Slp). It therefore appears that *hairy* stripes 3-7 are a direct output of the gap system.

#### Regulation of *even-skipped*

##### Regulatory elements

Like *hairy, eve* also possesses a full set of stripe-specific elements (1, 2, 3+7, 4+6, 5; reviewed in Schroeder et al. 2011). It also possesses a “late” element generating strong expression in seven narrow stripes [23,24]. However the expression of the late element does not kick in until the end of cellularisation, after the primary stripes of the secondary pair-rule genes *prd* and *slp* have already emerged [26]. The switchover from the stripe-specific elements to the late element appears to be regulated by Opa [6]. The *eve* pattern is therefore likely to be specified by gap inputs during cellularisation, with pair-rule inputs taking control at gastrulation.

##### No evidence for regulation by Hairy or Ftz

In wild-type embryos, *eve* and *hairy* expression overlaps throughout segmentation (Carroll et al. 1988; Hooper et al. 1989; see top row of Figure 6), so *eve* is evidently not repressed by Hairy. Consistent with this interpretation, *eve* expression is not directly affected by expression of Hairy fused to an activator domain [149], and eve expression is normal until late cellularisation in hairy mutant embryos (Appendix 1–figure 3).

**Appendix 1–figure 3:**
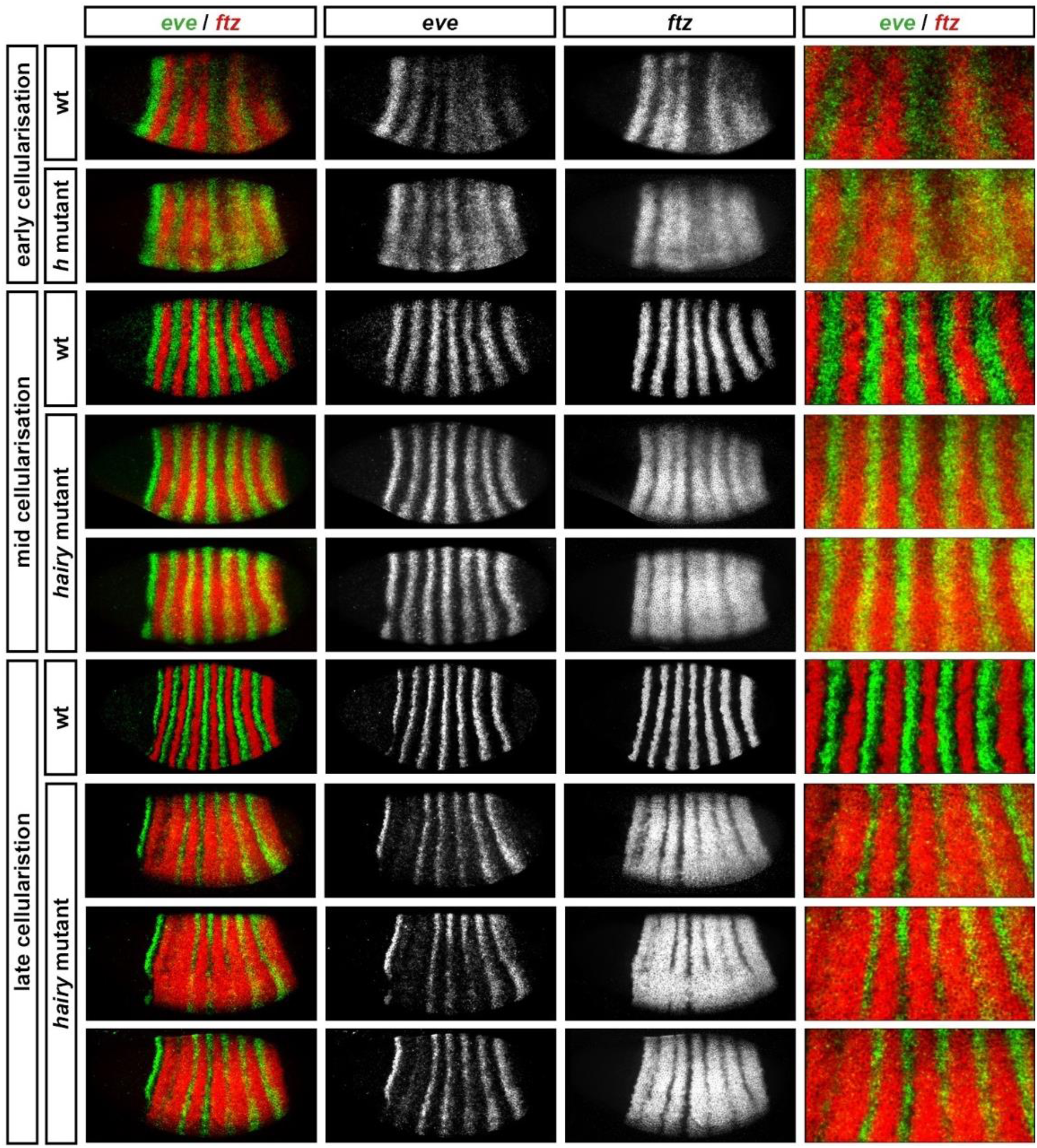
*ftz* is regulated by Hairy but *eve* is not. Relative expression of *ftz* and *eve* in wild-type and *hairy* mutant embryos at early, mid-, and late cellularisation. *ftz* expression expands dramatically in the mutant embryos, while *eve* expression is normal until late cellularisation. Two different mutant embryos are shown at mid-cellularisation, and three at late cellularisation. Single channel images are shown in greyscale in the central panels, and enlarged images of stripes 2-6 are shown at the right.

*eve* is also not repressed by Ftz: *eve* expression is not repressed by ectopic Ftz at any stage of segmentation [86], nor activated by ectopic Ftz fused to an activation domain [153], and *eve* expression does not change in *ftz* mutants [33]. Therefore, there is clear cut evidence that neither Hairy nor Ftz directly regulate *eve*.

##### Little evidence in favour of regulation by Runt or Odd

In contrast, mutant and misexpression studies indicate that both Runt and Odd can repress *eve* expression [34,60,145]. However, in order to determine whether these regulatory interactions are relevant to the early pair-rule network, it is important to analyse the timing of any changes to *eve* expression.

All *eve* stripes are effectively repressed by ectopic Odd or Runt in late cellularisation stage embryos [60,145]. However, during mid-cellularisation only *eve* stripe 1 is repressed by HS-Odd [145,154], and only *eve* stripe 2 is significantly repressed by HS-Runt [150]. Minor changes to some of the other *eve* stripes also occur in cellularisation stage HS-Runt embryos [150], but the time at which these changes were observed (30-40 minutes after the end of a 20 minute heatshock) is consistent with them being indirect responses to Runt. Runt activity is known to affect the gap system [150,151], therefore it is possible that the observed changes to *eve* expression are mediated by misexpressed gap factors.

The evidence from misexpression experiments therefore suggests that *eve* expression is not directly regulated by Runt or Odd until late cellularisation, aside from stripe-specific effects on stripes 1 and 2. This conclusion is also supported by analysis of the evidence from mutant embryos. *eve* expression is normal in *odd* mutant embryos [32] and also in embryos deficient for the entire *odd, sob, drm* cluster of *odd-skipped* paralogs (my data, not shown). In *runt* mutant embryos, the *eve* stripes show abnormal spacing (Frasch & Levine 1987; Ingham & Gergen 1988; Vavra & Carroll 1989; Warrior & Levine 1990; Jaynes & Fujioka 2004), but this is likely to be an indirect effect, resulting from regulatory effects of the early broad Runt domain on the gap system (see above). Fairly regular *eve* stripes are maintained until late cellularisation, when *eve* expression expands markedly (Appendix 1-figure 4). This delay is further evidence that Runt is not important for patterning *eve* until late cellularisation.

**Appendix 1–figure 4:**
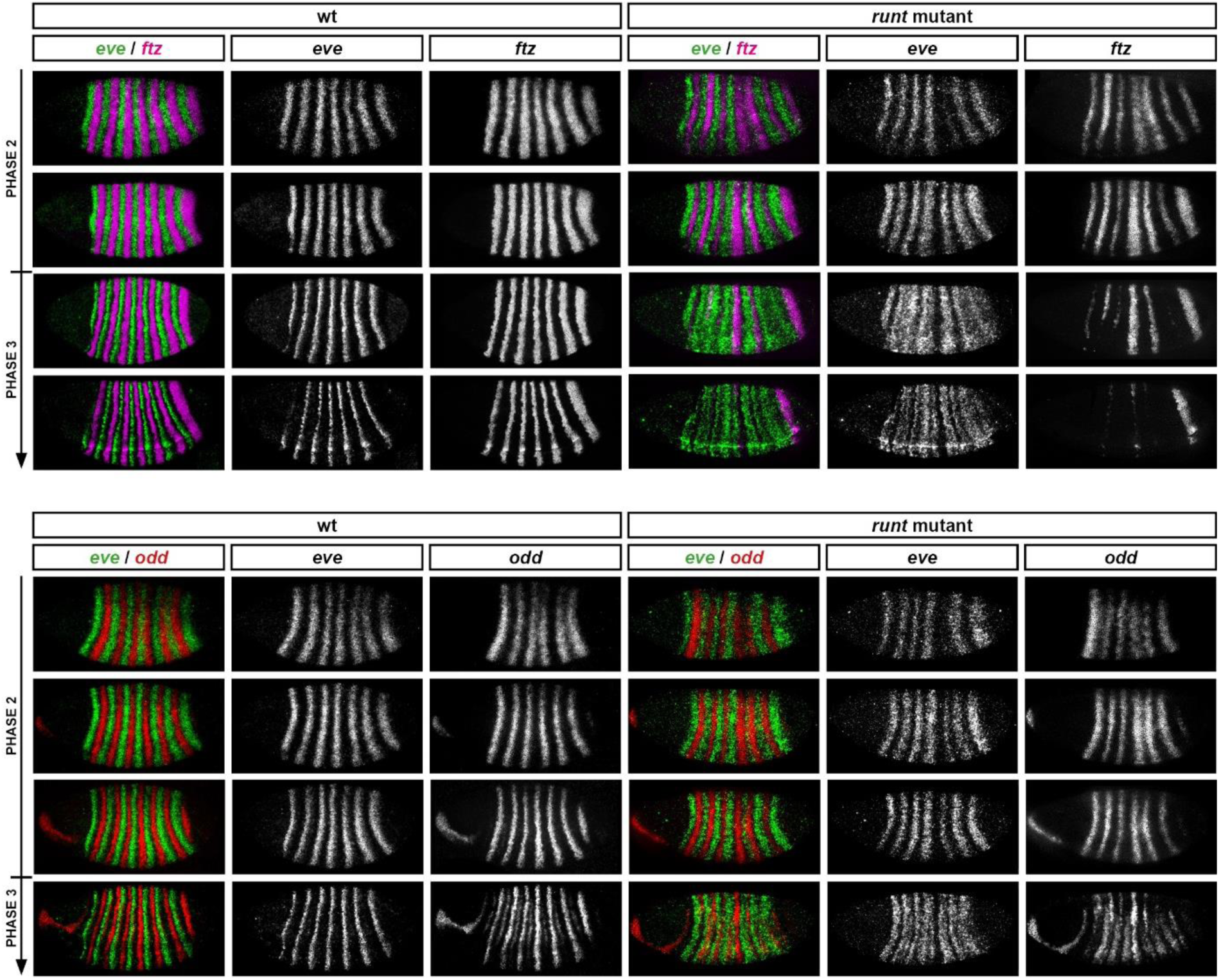
*ftz* and *odd* expression remains out of phase with the *eve* stripes in *runt* mutant embryos, despite irregularities in stripe spacing. Expression of *ftz* and *odd* relative to *eve* in wild-type and *runt* mutant embryos. Note that strong expression of *ftz* and *odd* stripes 4 and 5 in *runt* mutant embryos corresponds to an absence of *eve* and *hairy* expression in these regions (compare Appendix 1-figure 2). (These stripes fade only at gastrulation, presumably due to repression from newly synthesised Slp protein.) Single channel images are shown in greyscale to the right of the double channel images. Arrows indicate increasing developmental age.

This conclusion is also supported by observations from *hairy* mutant embryos, which exhibit significant coexpression of *eve* and *runt* (Appendix 1–figure 5). Anterior expansion of the *runt* stripes in these embyos means that *runt* is expressed throughout the *eve* stripes for most of cellularisation, however, aside from in stripe 2, *eve* expression is not significantly repressed until late cellularisation. *eve* transcript expression is also likely to overlap with Runt protein expression during cellularisation in wild-type embryos, although not so extensively. (Note that while overlaps are obvious between Eve protein and Runt protein [156] and between *eve* transcript and *runt* transcript (Appendix 1–figure 5), an *eve* RNA/Runt protein double would be required for explicit confirmation that *eve* is expressed in Runt-positive cells in wild-type embryos.)

**Appendix 1–figure 5:**
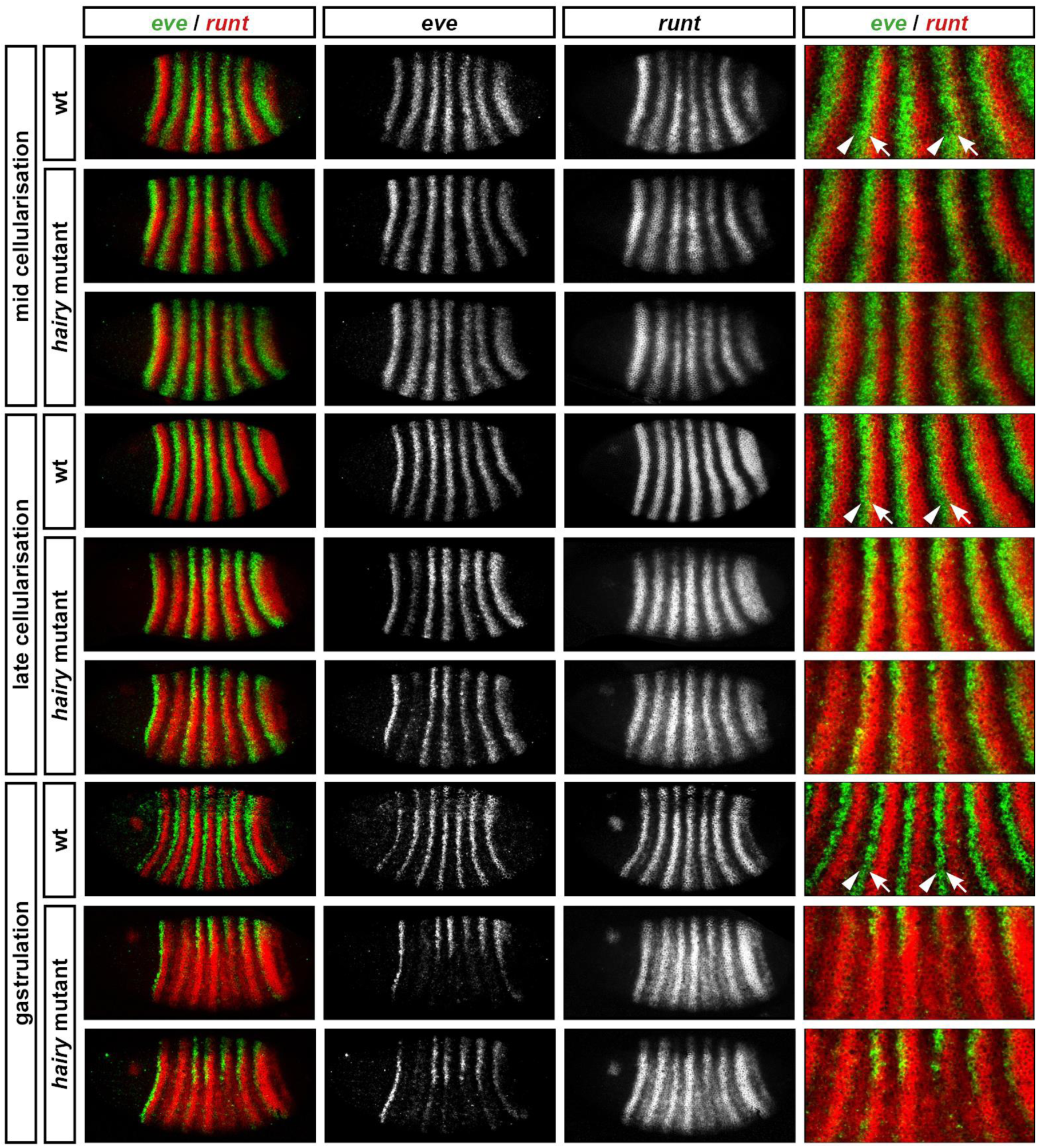
*runt* stripes are shifted anteriorly in *hairy* mutant embryos compared to wild-type. Relative expression of *eve* and *runt* in wild-type and *hairy* mutants, at mid-cellularisation, late cellularisation, and gastrulation. Two different mutant embryos are shown for each time point. Arrowheads in wild-type embryos indicate the anterior border of an *eve* stripe; arrows indicate the anterior border of a *runt* stripe. In the mutant embryos the *runt* stripes shift anteriorly and eventually entirely overlap the whole width of the *eve* stripes. In the mutant embryos, *eve* stripe 2 is repressed at late-cellularisation, however *eve* stripes 3-7 are not lost until gastrulation. Note the low level *runt* expression in between the stripes, which appears from late cellularisation in the mutant embryos. Single channel images are shown in greyscale in the central panels, and enlarged views of stripes 2-6 are shown at the right.

##### Conclusion

In summary, perturbing the expression of other pair-rule genes does not cause widespread gain or loss of *eve* expression until late cellularisation, suggesting that they do not directly regulate *eve* prior to this. Absence of Runt activity perturbs the spacing of the *eve* stripes, but, as discussed above, this effect is likely to be indirect (although I would not rule out a subtle role for Runt in quantitatively regulating/refining the *eve* stripes in wild-type). The precise, regularly spaced *eve* stripes in cellularising embryos therefore appear to be largely a direct output of the gap system. Note however that, as seen for *hairy* stripes 1 and 2, pair-rule cross-regulation does seem to be important for *eve* stripes 1 and 2 (which are sensitive to Odd and Runt, respectively). These effects might be mediated via the *eve* stripe 1 and *eve* stripe 2 elements, which would therefore take both gap and pair-rule inputs. Obvious pair-rule control of *eve* expression in stripes 3-7 does not become evident until late cellularisation, and is presumably mediated by the *eve* late element.

#### Regulation of *runt*

##### Regulatory elements

*runt* has both a full set of stripe-specific elements and a zebra element (reviewed in Schroeder et al. 2011). There are individual elements for all seven stripes, although the elements for stripes 1 and 2 also drive some expression in stripe 7 [26]. The zebra element is expressed during both cellularisation and gastrulation [82]. The boundaries of the wild-type *runt* stripes could therefore plausibly come from either the gap system or the pair-rule system, depending on how these various elements interact.

##### No evidence for regulation by Ftz

In cellularising wild-type embryos, *runt* expression overlaps both *eve* and *ftz* expression (Figure 2B,C), suggesting it is not repressed by either Eve or Ftz. This conclusion is largely supported by the evidence from mutant and misexpression studies. Consistent with Ftz not regulating *runt,* HS-Ftz has no direct effect on *runt* expression at any stage of segmentation [86], and *runt* expression is normal during cellularisation in *ftz* mutant embryos (Klingler & Gergen 1993).

##### No clear-cut evidence for regulation by Eve

The evidence relating to Eve is more complicated. In *eve* mutant embryos a fairly normal pair-rule pattern of *runt* forms initially, although several of the stripes are subsequently downregulated (Appendix 1–figure 6; Figure 10). As argued below and in the main text, this repression is likely indirect, mediated by ectopic Odd. *runt* expression is also affected by ectopic Eve, although the effect is variable depending on the stage at which Eve is misexpressed [59]. HS-Eve represses *runt* stripes 1-6 in gastrulating embryos, as expected from the sharp boundaries between *eve* and *runt* expression that form at late cellularisation in wild-type (see Appendix 1-figure 5). However, heatshocks in younger embryos can cause a dramatic broadening of the *runt* stripes, implicating Eve as an activator of *runt.* It is not clear whether this latter effect is direct or indirect, nor at which point exactly the switch from activation to repression occurs. Note though that the Eve protein is not known to act as a transcriptional activator [59,100,157–159].

**Appendix 1–figure 6:**
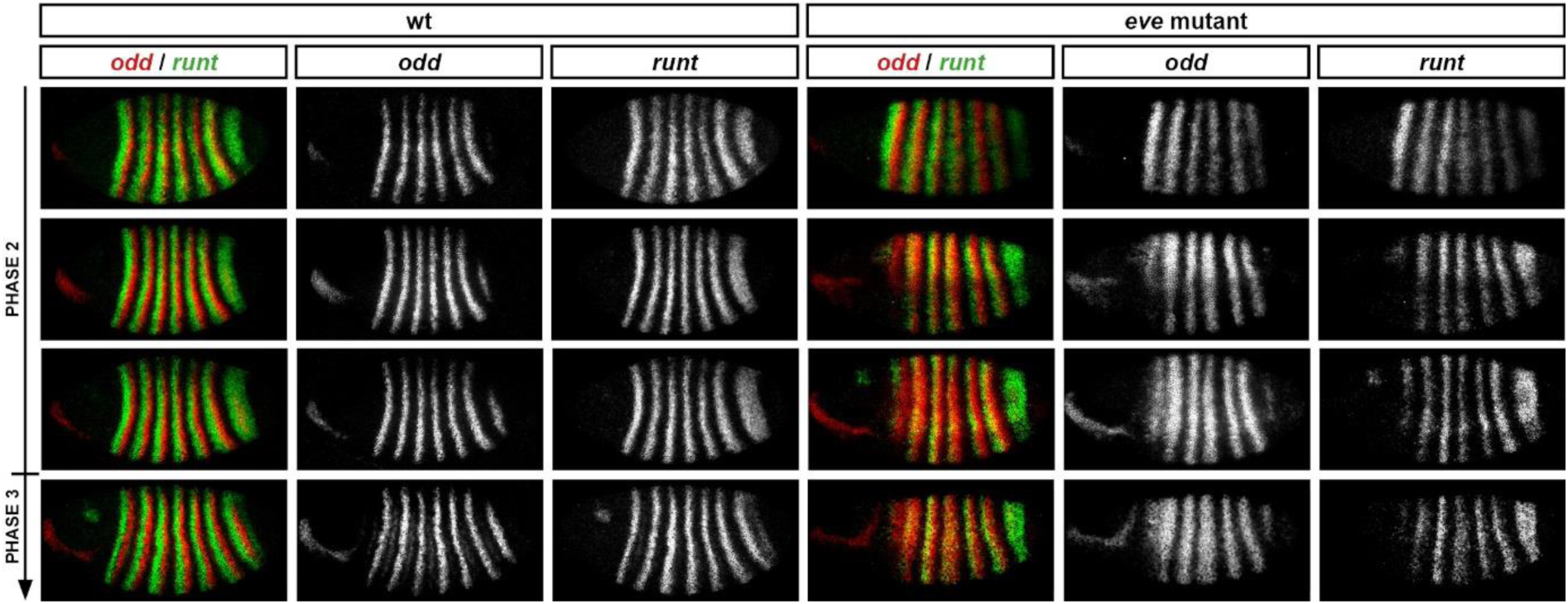
Relative expression of *runt* and *odd* in wild-type and *eve* mutant embryos during phase 2. *odd* stripes are expanded anteriorly in *eve* mutant embryos, overlapping the whole width of the runt stripes, rather than just their posteriors as in wild-type. *runt* expression in stripes 1-6 is downregulated in *eve* mutant embryos compared to wild-type embryos, due to repression from the ectopic Odd. Single channel images are shown in greyscale to the right of the double channel images. Arrow indicates increasing developmental age.

##### Good evidence for regulation by Hairy and Odd

In cellularising wild-type embryos the anterior and posterior borders of the *runt* stripes correspond closely to borders of *hairy* and *odd* expression, respectively (Figure 2E,H). Consistent with this stripe phasing, I find good evidence that both Hairy and Odd pattern *runt* expression during cellularisation, although the conclusions I draw are somewhat different than previous analyses.

In *hairy* mutant embryos, *runt* is expressed in a fairly normal seven stripe pattern, with weak expression in between the stripes (Ingham & Gergen 1988; Klingler & Gergen 1993; Appendix 1-figure 5). Because this pattern still contains seven well-defined stripes, it has been previously interpreted as representing the normal expression from the stripe-specific elements overlain on a background of low-level ectopic expression from a derepressed zebra element [26,106]. Under this view, the spatial pattern of *runt* expression in wild-type embryos would be determined mainly by the gap system, while the zebra element would play only a minor, redundant role.

However, I interpret this pattern of *runt* expression in *hairy* mutant embryos differently. Direct comparison with the *eve* stripes indicates that the strong stripes of *runt* shift anteriorly relative to their normal positions by around 1-2 nuclei, and are therefore not equivalent to the stripes observed in wild-type (Appendix 1–figure 5). This indicates that repression from Hairy normally specifies the anterior boundaries of the *runt* stripes in wild-type embryos, presumably through the *runt* zebra element. Protein fusion misexpression experiments indicate that this regulatory interaction is direct [149].

The evidence in favour of repression by Odd is fairly straightforward. *runt* expression is partially repressed by HS-Odd during cellularisation, while in *odd* mutant embryos the *runt* stripes broaden slightly [145]. This broadening presumably occurs at the posteriors of the *runt* stripes and reflects activation of *runt* expression in nuclei which are Odd positive but Hairy negative in wild-type, and therefore free of both Odd and Hairy in the mutant embryos. As discussed in the main text, derepression of *odd* expression in *eve* mutant embryos leads to a subsequent downregulation of the *runt* stripes, although this repression of *runt* is not total (Appendix 1–figure 6; Figure 10). Repression by Odd is also likely to be responsible for much of the residual periodicity of *runt* expression seen in *hairy* mutant embryos (Appendix 1–figure 5).

##### Conclusion

In summary, although *runt* possesses a full set of stripe-specific elements, the precise spatial regulation of its expression at mid cellularisation seems to be determined mainly by pair-rule inputs, specifically repression from Hairy and Odd. Positional information from these pair-rule factors seems to largely override spatial cues from the gap system in determining stripe boundaries, as demonstrated by expanded *runt* expression in *hairy* and *odd* mutants. The *runt* zebra element is therefore probably more important for patterning than are the stripe-specific elements, and indeed it is sufficient for fairly normal segmentation in their absence (Butler et al. 1992). However, it is clear that the stripe-specific elements do exert some influence on *runt* expression throughout cellularisation, as the control of Hairy and Odd over the *runt* expression pattern is not total. For example, the *runt* stripes are only downregulated in HS-Odd and *eve* mutant embryos rather than completely lost, while in wild-type embryos *runt* stripe 3 emerges from within a domain of Hairy expression. Therefore, while accurate to a first approximation, the model of early *runt* regulation depicted in Figure 1A is clearly an oversimplification of the more elaborate control logic seen in the embryo.

#### Regulation of *ftz* and *odd*

I analyse the regulation of *ftz* and *odd* simultaneously, because they exhibit very similar expression during cellularisation in a variety of genetic and experimental backgrounds. Any patterning differences between the two genes are noted and discussed.

##### Regulatory elements

In contrast to *hairy, eve* and *runt,* the genes *ftz* and *odd* have traditionally been considered secondary pair-rule genes, regulated by other pair-rule factors rather than by the gap system [17,30,58]. However, more recent analyses have revealed that their early expression is regulated by stripe-specific elements, and they are now classified as primary pair-rule genes [26,81]. Despite this status upgrade, they are evidently not as extensively regulated by the gap system as are the other three primary pair-rule genes. Neither *ftz* nor *odd* possesses a full set of stripe-specific regulatory elements: *ftz* has 1+5, 2+7 and 3+6, and so lacks an element for stripe 4, while *odd* has 1+5 and 3+6, and so lacks elements for stripes 2, 4 and 7 [26]. Both genes also possess a zebra element expressed throughout cellularisation [26,27]. The zebra elements are solely responsible for patterning the stripes that do not have their own stripe-specific elements. However, the boundaries of the remaining stripes could be plausibly specified by either gap factors or pair-rule factors, depending on how the elements interact.

##### Strong evidence for regulation by Eve and Hairy

Cross-regulatory interactions with other pair-rule genes have long been recognised to play an important role in determining the expression of *ftz* and *odd* during cellularisation, as their expression tends to be strongly perturbed in pair-rule mutant embryos (for example, for *ftz,* see Howard & Ingham 1986; Carroll & Scott 1986).

In wild-type embryos, the anterior borders of the *ftz* and *odd* stripes are closely associated with the posterior borders of the *eve* stripes, while the posterior borders of the *ftz* and *odd* stripes are closely associated with the anterior borders of the *hairy* stripes (Figure 2F,G,I,J). These patterns suggest that *ftz* and *odd* are repressed by both Eve and Hairy. This interpretation is supported by the expression of *ftz* and *odd* in mutant embryos.

In *eve* mutant embryos, *ftz* and *odd* are expressed in periodic patterns that are fairly complementary with the *hairy* stripes (Appendix 1-figure 1). Notably, fusions of *odd* stripes 1+2 correlate with the loss of *hairy* stripe 2 discussed above, indicating that the periodicity of *odd* expression in these embryos relies on repression by Hairy. In addition, the clear gaps between the posteriors of the *hairy* stripes and the anteriors of the *odd* stripes that are seen in wild-type embryos (asterisks in Appendix 1-figure 1) are lost, indicating that these are usually established in response to repression by Eve.

Expression changes between wild-type and *eve* mutant embryos are not so obvious for *ftz,* consisting of slight broadening of certain stripes (particularly 2 and 4), plus almost complete loss of stripe 1 *(odd* stripe 1 is also lost ventrally). However, the fact that stripes 2-6 of *ftz* and *odd* are expressed in extremely similar patterns to each other in *eve* mutant embryos (Appendix 1–figure 7) indicates that both genes are subject to the same patterning by Hairy. (The stripes of *odd* are consistently slightly broader than those of *ftz,* suggesting that *odd* is repressed slightly less effectively by Hairy.) I have not investigated the the differential expression of *ftz* stripe 1 and *odd* stripe 1 in *eve* mutant embryos, but this phenomenon indicates that their stripe 1 elements are each subject to their own bespoke regulation.

**Appendix 1–figure 7:**
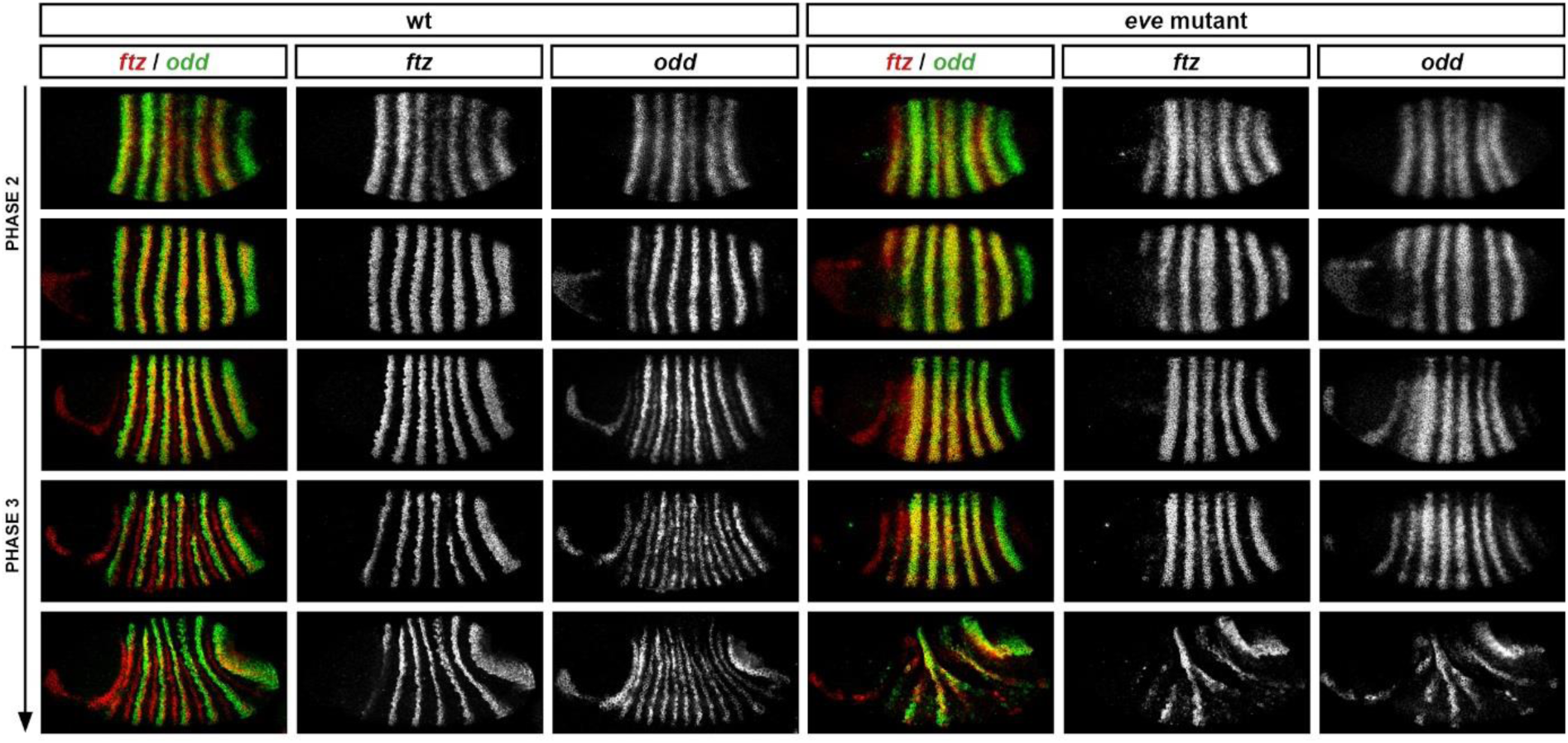
*ftz* and *odd* are expressed very similarly to each other in *eve* mutant embryos. Relative expression of *ftz* and *odd* in wild-type and *eve* mutant embryos. Stripes 2-6 of the two genes coincide exactly from phase 2 onwards. Single channel images are shown in greyscale to the right of the double channel images. Arrow indicates increasing developmental age.

The broadening of *ftz* stripe 4 (which lacks a stripe-specific element) in *eve* mutants is consistent with the anterior boundary of this stripe normally being patterned by repression from Eve. However, it is likely that the anterior boundaries of the remaining *ftz* stripes are initially positioned by gap factors so as to slightly overlap with Eve expression in wild-type embryos (see discussion of this topic in Clark & Akam 2016b). This would explain why they are located slightly anterior to the *odd* anterior boundaries in wild-type embryos, and why they do not significantly expand in *eve* mutant embryos

The evidence from *hairy* mutants is more dramatic. In these embryos, *ftz* and *odd* expression initially expands throughout almost the entire trunk (Appendix 1–figure 8), indicating that general repression by Hairy is crucial for their patterning during early cellularisation. As previously noted [152], this derepression is more extensive than would be predicted based on the spatial pattern of Hairy expression in fixed embryos, and therefore likely contributes to the severe and variable cuticle phenotypes of *hairy* null mutants [15,99].

**Appendix 1–figure 8:**
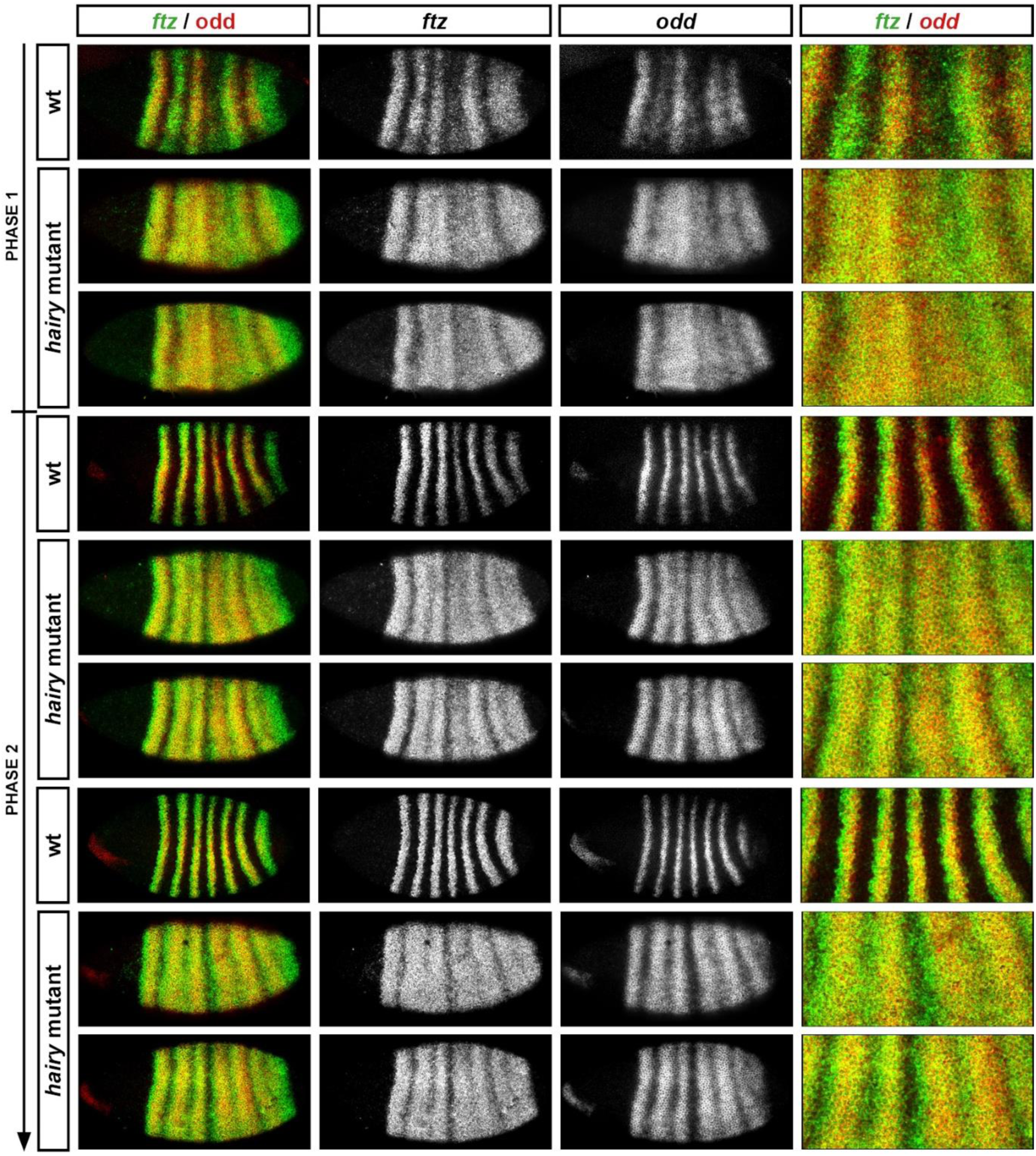
*ftz* and odd are ectopically expressed throughout the trunk in *hairy* mutant embryos. Relative expression of *ftz* and *odd* in wild-type and *hairy* mutant embryos during the first half of cellularisation. Broad ectopic expression of both genes appears early. Single channel images are shown in greyscale to the right of the double channel images. Arrow represents increasing developmental age (phase 1 until mid phase 2).

Direct repression of *ftz* and *odd* by Hairy and Eve is also supported by evidence from heatshock-mediated misexpression: both genes are repressed by HS-Eve (Manoukian and Krause 1992), and by HS-Hairy [89,90]. In addition, both genes are ectopically expressed in response to expression of Hairy fused to an activator domain [149]. Notably, odd is more effectively repressed by HS-Eve than is ftz, a difference that has been suggested to stem from different inherent sensitivities of *ftz* and *odd* to Eve activity [59]. However, this phenomenon could equally stem from Ftz autoactivation [87,88], and a resulting difficulty in turning *ftz* expression off once it has already been established.

##### No evidence for regulation by Runt

In wild-type embryos, the stripes of *ftz* and *odd* overlap the posteriors of the *runt* stripes during cellularisation (Figure 2C,H), suggesting that *ftz* and *odd* are not repressed by Runt. Consistent with this conclusion, *ftz* is not repressed after Runt misexpression, nor activated by Runt fused to an activator domain (Manoukian & Krause 1993; Jiménez et al. 1996; effects on *odd* not reported).

However, *ftz* and *odd* do exhibit altered expression in *runt* mutants during cellularisation, notably a weakening of stripes 3 and 6 (Carroll & Scott 1986; Ingham & Gergen 1988; Lawrence & Johnston 1989; Vavra & Carroll 1989; Jaynes & Fujioka 2004; Appendix 1-figure 4). Again, as discussed for the other pair-rule genes, this effect of Runt on stripe width and spacing appears to be indirect. In the mutant embryos, the patterns of *ftz* and *odd* still correspond negatively with those of *eve* and *hairy* (Appendix 1–figure 2; Appendix 1–figure 4), with the effects on stripes 3 and 6 apparently reflecting the partial fusion of *hairy* stripes 3-4 and 6-7, as well as more subtle changes to the relative phasings of the Hairy and Eve stripes (see Warrior & Levine 1990). It thus seems clear that *ftz* and *odd* are directly repressed by Hairy and Eve, but not by Runt.

##### Regulation by each other

Interestingly, Ftz and Odd appear to directly activate each other during early cellularisation: stripes of *ftz* broaden shortly after Odd misexpression, and vice versa [86,145]. However, all seven stripes of *ftz* or *odd* still appear (albeit weakened slightly) in embryos mutant for the other gene, indicating that this activation is not necessary for their expression [26,62,86,145].

##### Conclusion

In conclusion, the stripes of *ftz* and *odd* are largely defined by pair-rule cross-repression, presumably via their zebra elements. The posterior boundaries of the stripes of both genes are defined by repression from Hairy, while the anterior boundaries of the *odd* stripes are defined by repression by Eve. The anterior boundaries of the *ftz* stripes seem to be defined by Eve in certain cases, but by gap inputs in others.

The significant role of the zebra elements explains why *ftz* and *odd* need not possess a full set of stripe-specific elements: the necessary spatial information for patterning their stripes can be provided instead via Eve and Hairy. However, it is clear that certain stripe-specific elements do play non-redundant roles in patterning: for example, establishing *ftz* and *odd* stripe 3 expression despite the late-resolving Hairy pattern in this region, or helping to differentially position the anterior boundaries of the *ftz* and *odd* stripes.

It is also clear that there are still questions to be answered about the regulation of *ftz* and *odd* (particularly of *ftz)* during cellularisation. How do the stripe-specific and zebra elements interact, what is the role of Ftz autoactivation in patterning the *ftz* stripes, and what is the explanation for the surprisingly crucial role for Hairy in generating a periodic output pattern?

### Appendix 2: Details of models and simulations

This appendix provides details of the gene regulatory network models and simulation algorithm used to generate the simulations shown in Movies 1-13. Output data from these simulations are also presented in Figures 2, 4–6, 10, 12, and 13.

I first provide overviews of the general modelling framework (theory), and its specific implementation (software). I then list the details (control logic, time delay values, and initial conditions) of each individual simulation. Where necessary, I discuss the way in which I have translated inferred genetic interactions into a formalised model, pointing out any ad hoc or uncertain assumptions. I also highlight situations where the regulation in the embryo is clearly more nuanced than can be captured by the simple models I use.

#### General modelling framework

I use a Boolean network approach [143, 161–164] to model the interactions between a number of transcription factors in a given cell. In order to capture some of the essential “biology” of gene expression, cell state is separated into two distinct layers, roughly equivalent to transcript and protein respectively (see Thomas 1991). The current state of the “protein” layer determines the transcriptional output from the cell at the next timepoint, based on the control logic of the network. The state of the “transcript” layer is in turn converted into protein state at some later timepoint. Each component of the network is associated with specific time delays that govern the conversion process between transcript state and protein state, allowing different gene products to exhibit different “synthesis speeds” and/or “protein stability”. Given some set of initial conditions, cell state will evolve deterministically over time, eventually arriving at a stable state or a limit cycle.

Each component in the network (i.e. each transcription factor) is represented by a Boolean variable. If the variable equals 0, the component is “off’, while if the variable equals 1, the protein is “on”. Given *n* components in the network, there are therefore *2^n^* possible network states. (If helpful) these possible states can be thought of as the vertices of an *n*-dimensional hypercube. A particular network state (protein state) can be specified by a one-dimensional array of length n, where the value (0 or 1) of index *i* gives the expression state of the *i*th component in the network. For example, for a network of three gene products A,B, and C, the array [1,0,1] would represent the state where A and C are on, and B is off.

The transcriptional output of each particular component in the network can be calculated from the protein state array by a specific logical function (the “control logic” of the gene). Each network component will have its own logical function, which may take into account the protein states of all, some, or none of the components in the network, including itself. There is no constraint on the form of the logical functions, so long as they map each of the 2^n^ possible network states to a specified level of transcription (0 or 1). Combinatorial regulatory interactions are therefore easily represented, although quantitative regulatory effects are ruled out by definition.

Stable states of the network can be easily calculated from the overall set of protein state → transcriptional output mappings; they are those where the output state for each component is unchanged from its input state. These stable attractor states can be thought of as representing particular cell “fates”.

The network is updated synchronously, by discrete timesteps. The transcriptional output of the network at timepoint *t*+1 is simply calculated from the network protein state at time t. However, calculating the new protein state is more complicated, because of the time delays that need to be accounted for.

Each component in the network is associated with two positive integer parameter values, a “synthesis delay” and a “decay delay”. The synthesis delay determines how long a component must be transcribed before the component turns on at the protein level, while the decay delay determines how long a component has to be transcriptionally silenced before the component turns off. Each component is also associated with two non-negative integer variables, “transcript age” and “protein age” respectively, that provide some “memory” to the system.

The particular rules governing the conversion process for a given network component are as follows:

~~~
if (*transcript=*0 and *old_protein*=0) then *new_protein* → 0, *transcript_age* → 0
if (*transcript*=1 and *old_protein*=0) then increment *transcript_age:*
       if (*transcript_age* < *synthesis_delay*) then *new_protein* → 0; else *new_protein* → 1
if (*transcript=*1 and *old_protein=*1) then *new_protein* → 1, *protein_age* → 0
if (*transcript=*0 and *old_protein=*1) then *transcript_age* → 0, increment *protein_age:*
       if (*protein_age* < *decay_delay*) then *new_protein* → 1; else *new_protein* → 0, protein_age → 0
~~~

Note: *“transcript”* = transcriptional output at timepoint *t*+1; *“old_protein”* = protein state at timepoint *t*; *“new_protein”* = protein state at timepoint *t*+1.

The simulation algorithm is thus set up so that a protein will only turn on if there is continuous transcription for the entire duration of the synthesis delay, and will only turn off is transcription is continuously absent for the entire duration of the decay delay, effectively ignoring very brief changes to transcriptional output.

#### Implementation

The simulation software is written in object-oriented Python (www.python.org), using the additional libraries NumPy and Matplotlib [137,138]. All the functions necessary for producing and visualising a simulation are contained in one .py file (Supplementary File 1). The user simply specifies a network model and simulation parameter values in a separate script (see Supplementary File 2 for a template), and can thereby easily carry out bespoke simulations. Simulations can be run simultaneously for an arbitrary number of “cells”, each of which behaves independently. Each cell has the same network model and parameter values, but can be assigned different initial conditions.

The things the user must specify are:

1) The network components (i.e. the various transcription factors in the system)
2) The control logic (i.e. regulation) of each component. This is done simply by defining a function that takes the current state of the network as its argument and returns either 0 or 1 as appropriate.
3) The synthesis and decay delays associated with each component. There are also functions to set default values for all components, if desired.
4) The number of “cells” to be simulated.
5) The initial conditions of each cell. Protein state, protein age, transcript state, and transcript age may all be specified. Values are set to zero by default.
6) The number of timepoints to simulate.
7) If visualising the output with an animation, the colour associated with each gene product, and other aesthetic preferences.

The user then simply has to call the simulation function to calculate the system state at each timepoint, given the chosen initial conditions, control logic, and parameter values. The user may then export or visualise the output data.

Visualisations represent individual cells as different columns (named C1, C2, C3…) and different gene products as different named rows. Both “transcript” and “protein” outputs are shown for each row. Depending on user preference, transcript and protein ages can also be represented in the visualisation, with “decaying” proteins shown as increasingly transparent before they turn off, and “nascent” transcripts shown as increasingly opaque until protein appears.

Visualisations can be shown as an animation, or the individual frames may be saved as image files. The frames can then be stitched together to produce video files, for example using ffmpeg (www.ffmpeg.org).

#### Pair-rule model details

Each simulation referenced in the main text models pair-rule gene expression along an idealised anteroposterior axis. The simulations show an array of 20 “cells”, each of which evolves independently. This cellular autonomy is justified for the model by the apical localisation of real pair-rule transcripts, which precludes diffusion between nuclei during cellularisation of the blastoderm [104,105]. (Note also that all the pair-rule genes code for transcription factors, rather than e.g. components of signalling pathways.) Initial conditions for each simulation consist of a pattern that repeats every 8 cells, with each group of 8 cells representing an idealised double segment repeat.

“Early network” simulations (Movies 1–8) are the simplest, restricted to modelling the establishment and maintenance of the original pair-rule pattern observed at mid-cellularisation. They model the expression of the five primary pair-rule genes *(hairy, eve, runt,* ftz, and *odd),* as patterned ultimately by abstracted gap inputs (e.g. “G1”, “G2”).

“Whole system” simulations (Movies 9-11) are more complicated, additionally including the patterning of the secondary pair-rule genes, and the transition to single-segment periodicity. As well as the primary pair-rule genes and the gap inputs, they model the secondary pair-rule genes *prd, slp,* and the segment-polarity gene *en.* They also utilise two temporal signals, “Signal X” and Opa. Signal X is a hypothetical signal controlling the onset of secondary pair-rule gene expression, while Opa controls the switch between the early and the late pair-rule networks.

“Modified network” simulations (Movies 12 and 13) are based on the “whole system” simulations, but tweak the inferred *Drosophila* network so as to produce a more autonomous patterning system that can operate in the absence explicit gap inputs.

In all simulations, the seven pair-rule factors and En are all assigned identical synthesis and decay delays, both equal to 6 timesteps. Studies of pair-rule gene expression kinetics indicate that protein synthesis and turnover rates in the embryo are equally rapid, and are both processes occurring on the order of 5-10 minutes [83,86]. The time delays associated with the remaining system components (gap inputs, Signal X, and Opa) are chosen in an ad hoc manner so as to provide appropriate spatiotemporal signals to the pair-rule genes.

The control logic, time delays, and initial conditions for each simulation are described below. Note that any unspecified initial conditions can be assumed to be set to zero.

#### “Early network”simulations (Movies 1-8)

The core control logic of these simulations is the “early network” shown in Figure 1A. It is formulated as below:

~~~
*hairy:* if (G1=ON) then *hairy* → OFF; else *hairy* → ON
*eve:*   if (G2=ON) then *eve* → OFF; else *eve* → ON
*runt:*  if (Hairy=ON or Odd=ON) then *runt* → OFF, else *runt* → ON
*ftz:*   if (Hairy=ON or Eve=ON) then *ftz* → OFF, else *ftz* → ON
*odd:*   if (Hairy=ON or Eve=ON) then *odd* → OFF, else *odd* → ON
~~~

In each simulation, no initial pair-rule gene expression is specified (i.e. in each cell, for each of the five variables, the initial conditions are transcript=0, protein=0, transcript age=0, protein age=0).

##### Effects of shifting gap inputs (simulations 1-5)

Simulations 1 to 5 explore the effects of dynamic gap inputs on pair-rule gene expression. The only differences between them relate to the “gap inputs” G1 and G2.

###### Control logic

G1 and G2 autoactivate in simulation 1 so as to produce stable domains, and autorepress in simulations 2-5 so as to produce dynamic domains.

Simulation 1 (static gap inputs):

~~~
*g1:* if (G1=ON) then *g1* → ON; else *g1* → OFF
*g2:* if (G2=ON) then *g2* → ON; else *g1* → OFF
~~~

Simulations 2-5 (dynamic gap inputs):

~~~
*g1:* if (G1=ON) then *g1* → OFF; else *g1* → ON
*g2:* if (G2=ON) then *g2* → OFF; else *g1* → ON
~~~

###### Time delays

In each of the simulations 2-5, the time delays and initial conditions of G1 and G2 are specifically chosen so as to produce a particular gap domain shift rate while preserving an 8 cell pattern repeat. Simulation 2 produces a shift rate of one cell every 6 timesteps (equal to the synthesis/decay delay of the pair-rule factors), which requires G1/G2 expression to go through a full oscillation every 48 timesteps. Simulations 3-5 have time delays and initial conditions equivalent to those of simulation 2, except multiplied by a scaling factor to give slower or faster oscillations of G1/G2 in each cell and hence slower or faster shifts. Simulation 3 is scaled by 2, Simulation 4 is scaled by 1/2, and Simulation 5 is scaled by 1/3.

|  | Sim 1 | Sim 2 | Sim 3 | Sim 4 | Sim 5 |
| --- | --- | --- | --- | --- | --- |
| **G1/G2 synthesis delay** | 6 | 18 | 36 | 9 | 6 |
| **G1/G2 decay delay** | 6 | 30 | 60 | 15 | 10 |

###### Intitial conditions

Common to all simulations:

*Initial conditions:* Common to all simulations:
|  | C1 | C2 | C3 | C4 | C5 | C6 | C7 | C8 |
| --- | --- | --- | --- | --- | --- | --- | --- | --- |
| **G1 value** | 0 | 0 | 0 | 1 | 1 | 1 | 1 | 1 |
| **G2 value** | 1 | 1 | 0 | 0 | 0 | 1 | 1 | 1 |

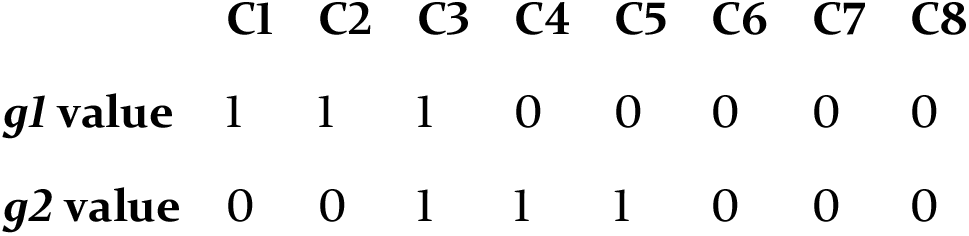
Common to simulations 2-4:

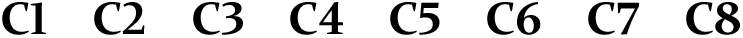
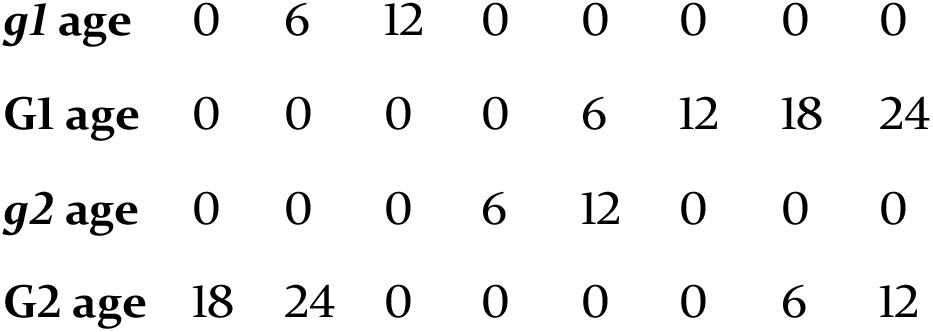
Specific to simulation 2:

Specific to simulation 2:
| C1 | C2 | C3 | C4 | C5 | C6 | C7 | C8 |
| --- | --- | --- | --- | --- | --- | --- | --- |

|  |  |  |  |  |  |  |  |  |
| --- | --- | --- | --- | --- | --- | --- | --- | --- |
| ***g1 age*** | 0 | 6 | 12 | 0 | 0 | 0 | 0 | 0 |
| **G1 age** | 0 | 0 | 0 | 0 | 6 | 12 | 18 | 24 |
| ***g2 age*** | 0 | 0 | 0 | 6 | 12 | 0 | 0 | 0 |
| **G2 age** | 18 | 24 | 0 | 0 | 0 | 0 | 6 | 12 |

Specific to simulation 3:

Specific to simulation 3:
|  | **C1** | **C2** | **C3** | **C4** | **C5** | **C6** | **C7** | **C8** |
| --- | --- | --- | --- | --- | --- | --- | --- | --- |
| ***g1 age*** | 0 | 12 | 24 | 0 | 0 | 0 | 0 | 0 |
| **G1 age** | 0 | 0 | 0 | 0 | 12 | 24 | 36 | 48 |
| ***g2 age*** | 0 | 0 | 0 | 12 | 24 | 0 | 0 | 0 |
| **G2 age** | 36 | 48 | 0 | 0 | 0 | 0 | 12 | 24 |

Specific to simulation 4:

Specific to simulation 4:
|  | **C1** | **C2** | **C3** | **C4** | **C5** | **C6** | **C7** | **C8** |
| --- | --- | --- | --- | --- | --- | --- | --- | --- |
| ***g1 age*** | 0 | 3 | 6 | 0 | 0 | 0 | 0 | 0 |
| **G1 age** | 0 | 0 | 0 | 0 | 3 | 6 | 9 | 12 |
| ***g2 age*** | 0 | 0 | 0 | 3 | 6 | 0 | 0 | 0 |
| **G2 age** | 9 | 12 | 0 | 0 | 0 | 0 | 3 | 6 |

Specific to simulation 5:

Specific to simulation 5:
|  | **C1** | **C2** | **C3** | **C4** | **C5** | **C6** | **C7** | **C8** |
| --- | --- | --- | --- | --- | --- | --- | --- | --- |
| ***g1 age*** | 0 | 2 | 4 | 0 | 0 | 0 | 0 | 0 |
| **G1 age** | 0 | 0 | 0 | 0 | 2 | 4 | 6 | 8 |
| ***g2 age*** | 0 | 0 | 0 | 2 | 4 | 0 | 0 | 0 |
| **G2 age** | 6 | 8 | 0 | 0 | 0 | 0 | 2 | 4 |

##### Effects of additional gap inputs (simulations 6-8)

These three simulations model situations where *runt* and/or *ftz/odd* are initially controlled by gap inputs, before being taken over the pair-rule network later on. These simulations involve some extra components. Simulation 6 has an additional gap input, G3, which regulates *runt,* while simulation 7 has an analogous gap input, G4, which regulates *ftz* and *odd.* Both G3 and G4 are present in simulation 8. All three simulations additionally have a temporal signal, “G” which controls whether *runt* and/or *ftz/odd* are controlled by the gap inputs G3/G4, or alternatively by pair-rule inputs as normal. Details of any new control logic and initial conditions are listed below. Anything not explicitly detailed is the same as simulation 2.

###### Control logic

Simulations 6-8

~~~
*g:   g* → OFF
~~~

Simulations 6 and 8

~~~
*g3:*    if (G3=ON) then *g3* → OFF; else *g3* → ON
*runt:*  if (G=ON):
              if (G3=ON) then *runt* → OFF; else *runt* → ON
       if (G=OFF):
              if (Hairy=ON or Odd=ON) then *runt* → OFF, else *runt* → ON
~~~

Simulations 7 and 8

~~~
*g4:*    if (G4=ON) then *g4* → OFF; else *g4* → ON
*ftz:*   if (G=ON):
              if (G4=ON) then ftz → OFF; else *ftz* → ON
       if (G=OFF):
              if (Hairy=ON or Eve=ON) then *ftz* → OFF, else *ftz* → ON
*odd:*   if (G=ON):
              if (G4=ON) then *odd* → OFF; else *odd* → ON
       if (G=OFF):
              if (Hairy=ON or Eve=ON) then *odd* → OFF, else *odd* → ON
~~~

Time delays.

|  | **Sim 6** | **Sim 7** | **Sim 8** |
| --- | --- | --- | --- |
| **G decay delay** | 12 | 6 | 6 |
| **G3 synthesis delay** | 18 | - | 18 |
| **G4 synthesis delay** | - | 18 | 18 |
| **G3 decay delay** | 30 | - | 30 |
| **G3 decay delay** | - | 30 | 30 |

Note: the G decay delay effectively specifies how long *runt* and/or *ftz/odd* are controlled by gap inputs rather than the pair-rule network.

###### Initial conditions

Common to simulations 6-8:

*Initial conditions:* Common to simulations 6-8:
|  | C1 | C2 | C3 | C4 | C5 | C6 | C7 | C8 |
| --- | --- | --- | --- | --- | --- | --- | --- | --- |
| **G value** | 1 | 1 | 1 | 1 | 1 | 1 | 1 | 1 |

Specific to simulations 6 and 8:

Specific to simulations 6 and 8:
|  | C1 | C2 | C3 | C4 | C5 | C6 | C7 | C8 |
| --- | --- | --- | --- | --- | --- | --- | --- | --- |
| **g3 value** | 0 | 0 | 0 | 0 | 1 | 1 | 1 | 0 |
| **g3 age** | 0 | 0 | 0 | 0 | 0 | 6 | 12 | 0 |
| **G3 value** | 1 | 1 | 1 | 1 | 0 | 0 | 0 | 1 |
| **G3 age** | 6 | 12 | 18 | 24 | 0 | 0 | 0 | 0 |

Specific to simulations 7 and 8:

Specific to simulations 7 and 8:
|  | C1 | C2 | C3 | C4 | C5 | C6 | C7 | C8 |
| --- | --- | --- | --- | --- | --- | --- | --- | --- |
| **g4 value** | 1 | 0 | 0 | 0 | 0 | 0 | 1 | 1 |
| **g4 age** | 12 | 0 | 0 | 0 | 0 | 0 | 0 | 6 |
| **G4 value** | 0 | 1 | 1 | 1 | 1 | 1 | 0 | 0 |
| **G4 age** | 0 | 0 | 6 | 12 | 18 | 24 | 0 | 0 |

##### Whole system simulations (Movies 9-11)

###### Control logic

The structure of the pair-rule network as a whole is based largely on evidence from Clark & Akam 2016, and summarised in Figure 1. The topology of the network is context dependent, with many genetic interactions affected by the transcriptional cofactor, Opa. Since Opa protein appears in the *Drosophila* embryo only at the end of cellularisation [80], segment patterning is sequentially directed by “early” and “late” incarnations of the overall network.

I therefore make the control logic of each segmentation gene in the network conditional on the presence or absence of Opa. If Opa is off, the control logic is a formulation of the early network (Figure 1A), while if Opa is on, the control logic is a formulation of the late network (Figure 1B). The specific regulatory rules for each pair-rule gene (plus en) are described below, supplemented with explanatory notes where appropriate.

~~~
*hairy:* if (Opa=OFF):
              if (G1=ON) then *hairy* → OFF; else *hairy* → ON
       if (Opa=ON):
              *hairy* → OFF
~~~

Notes: While Opa is off, the regulation of *hairy* is the same as in Simulations 1-8. As soon as Opa turns on, *hairy* expression turns off. Hairy protein plays no role in the late pair-rule network and is independent of pair-rule gene expression throughout the whole patterning process.

~~~
*eve*:   if (Opa=OFF):
              if (G2=ON) then *eve* → OFF; else *eve* → ON
       if Opa=ON:
              if (Runt=ON or Odd=ON or En=ON or (Slp=ON and Eve=OFF)) then *eve* → OFF; else *eve* → ON
~~~

Notes: While Opa is off, the regulation of *eve* is the same as in Simulations 1-8. As soon as Opa turns on, *eve* expression is instead patterned by various repressors (Runt, Odd, Slp, En). The qualification in the model that Slp represses *eve* only if Eve is not already turned on is required in order to resolve overlaps between *eve* and *slp* expression that form in the simulations just before the switch to the late network occurs. Specifying that Eve expression “wins” this contest ensures that the anterior boundaries of the Eve stripes define the parasegment boundaries, as occurs in the embryo [160]. However, the overlap situation is not a scenario that occurs in real embryos: the *eve* boundaries stabilise at roughly the same time *slp* turns on (rather than afterwards), and therefore the *eve* and *slp* domains are mutually exclusive throughout the segmentation process. A more realistic model that avoided this issue would need to include more sophisticated temporal control of *eve* and *slp* regulation, and/or time-dependent gap domain dynamics. Note that for simulation 11, which models an *eve* mutant embryo, the control logic of *eve* is replaced by unconditional repression (*eve* → OFF).

~~~
*runt:*   if (Opa=OFF):
               if (Hairy=ON or Odd=ON) then *runt* → OFF; else *runt* → ON
        if (Opa=ON):
               if ((Eve=ON and Runt=OFF) or (Odd=ON and Runt=OFF) or En=ON) then *runt* → OFF; else *runt* → ON
~~~

Notes: While Opa is off, the regulation of *runt* is the same as in Simulations 1-8. As soon as Opa turns on, *runt* expression is instead patterned by various repressors (Eve, Odd, En). However, I have specified that if Runt is coexpressed with either Eve or Odd, the Runt expression will “win” and *runt* transcription will not turn off. This is because Eve/Runt and Runt/Odd overlaps are both present during cellularisation (at the anteriors and the posteriors of the Runt stripes, respectively), and both resolve in favour of *runt* expression during gastrulation [6,144]. In contrast, En expression has the potential to supplant Runt expression, as occurs at the posterior borders of the *runt* primary stripes. Note that the inferred control logic is a phenomenological description of regulation within the embryo, and does not necessarily imply Runt autoregulation at a mechanistic level. Note also that the Opa-dependent control logic specified above reflects the late regulation of the *runt* “seven stripe element”, and I have not explicitly modelled the expression arising from the “six stripe element” [6,82].

~~~
*ftz:*   if (Opa=OFF):
              if (Hairy=ON or Eve=ON or Slp=ON) then *ftz* → OFF; else *ftz* → ON
       if (Opa=ON):
              if (Eve=ON or Slp=ON or Ftz=OFF) then *ftz* → OFF; else *ftz* → ON
~~~

Notes: While Opa is off, the regulation of *ftz* is the same as for Simulations 1-8, except with the addition of repression from Slp. (Note that this last interaction is not diagrammed in Figure 1A, since Slp protein is synthesised too late to have any practical effect on *ftz* expression while the early network is in operation.) When Opa turns on, *ftz* expression remains repressed by Eve and Slp, but is no longer regulated by Hairy. Instead, it requires autoactivation. Having this autoregulation in the model permits the anterior border of the Ftz domain to remain stable and define the even-numbered parasegment boundaries, as occurs in the embryo [160]. However, note that while direct Ftz autoactivation is well-documented [87,88], it remains to be tested whether Ftz activity is absolutely required for late Ftz expression.

~~~
*odd:*   if (Opa=OFF):
              if (Hairy=ON or Eve=ON or Slp=ON) then *odd* → OFF; else *odd* → ON
       if (Opa=ON):
              if (Runt=ON or En=ON or (Prd=ON and Odd=OFF)) then *odd* → OFF; else *odd* → ON
~~~

Notes: While Opa is off, the regulation of *odd* is the same as for Simulations 1-8, except with the addition of repression from Slp. (Note that this last interaction is not diagrammed in Figure 1A, since Slp protein is synthesised too late to have any practical effect on *odd* expression while the early network is in operation.) When Opa turns on, *odd* expression is no longer regulated by Hairy or Eve, and instead becomes repressed by Runt, En, and Prd. It remains repressed by Slp. I have specified that the repression by Prd only occurs if Odd expression is not already turned on – this reflects the observation that Prd activity patterns the nascent *odd* secondary stripes [62], but has no apparent effect on the *odd* primary stripes. This phenomenon – which does not necessarily imply Odd autoregulation at the mechanistic level – should be further investigated to test the inferred control logic.

~~~
*prd:*   if (Opa=OFF):
              if (Signal X=ON or Eve=ON) then *prd* → OFF; else *prd* → ON
       if (Opa=ON):
              if (Prd=ON and Odd=OFF) then *prd* → ON; else *prd* → OFF
~~~

Notes: While Opa is off, *prd* is repressed by Eve (which patterns where it is expressed) and by the hypothetical “Signal X” (which patterns *when* it is expressed). When Opa turns on, *prd* is no longer repressed by Eve, and instead repressed by Odd. It also requires autoactivation. The autoregulation is required in the model in order to maintain a stable posterior boundary within the odd-numbered parasegments. Note that while early repression by Eve and late repression from Odd are both strongly supported by experimental evidence, not much more is known about *prd* regulation. The control logic above is therefore largely speculative, and indeed does not perform particularly well in the simulations (see Figure 6–figure supplement 2). The regulation of *prd* therefore warrants further investigation.

~~~
*slp:*   if (Opa=OFF):
              if (Signal X=ON or Eve=ON or Runt=ON or Prd=OFF) then *slp* → OFF; else *slp* → ON
       if (Opa=ON):
              if (Eve=ON or En=ON or ((Ftz=ON or Odd=ON) and Slp=OFF)) then *slp* → OFF; else *slp* → ON
~~~

Notes: While Opa is off, *slp* is repressed by Eve, Runt, and Signal X, and requires activation by Prd. Slp expression is thus spatially patterned by Eve and Runt, and temporally patterned by Signal X and Prd. The requirement for activation by Prd means that *slp* expression follows *prd* expression after a short time lag, in agreement with real expression in the embryo. This stipulation in the model is not entirely ad hoc: *slp* expression is reduced in *prd* mutant embryos (data not shown), indicating that *prd* does contribute to *slp* activation. However, *slp* expression is not abolished in these embryos, indicating that other, as yet unidentified, temporal signals must also be involved in the process. When Opa turns on, *slp* is patterned by repression from Eve, En, Ftz, and Odd. The qualification that Ftz and Odd only repress *slp* if Slp expression is absent reflects the observation that expression overlaps between Slp and Ftz/Odd at the posteriors of the Ftz/Odd primary stripes are always resolved in favour of Slp. Again, note that this phenomenological description does not necessarily imply Slp autoregulation at the mechanistic level.

~~~
*en:*   if (Opa=OFF):
             *en* → OFF
      if (Opa=ON):
             if (Ftz=ON and Odd=OFF and Slp=OFF) or (Prd=ON and Odd=OFF and Slp=OFF and Runt=OFF)) then *en* → ON; else *en* → OFF
~~~

Notes: While Opa is off, *en* expression is repressed. When Opa turns on, *en* is spatially patterned by both activators and repressors. *en* requires activation from either Prd or Ftz, and is always repressed by Odd and Slp. The odd-numbered *en* stripes (those activated by Prd) are additionally repressed by Runt. In contrast, the even-numbered *en* stripes (those activated by Ftz) are insensitive to Runt activity.

The remaining components of the model (Signal X, Opa, G1/G2) provide broad or abstracted inputs into the periodically expressed components described above. (Of the four signals, only Opa is identified with a specific transcription factor.) Signal X and Opa are extrinsic inputs that are “off’ by default or “on” by default, respectively. G1 and G2 are abstracted, autoregulatory gap inputs, whose expression (and function) is restricted to early stages of patterning (i.e. when Opa is off). Note that G1 and G2 have different control logic in Simulation 9 (modelling static gap inputs) versus Simulations 10 and 11 (modelling dynamic gap inputa).

~~~
*signal x:      signal x* → OFF
*opa:           opa* → ON
*g1* (Sim 9):    if (Opa=OFF):
                      if (G1=ON) then *g1* → ON; else *g1* → OFF
               if (Opa=ON):
                      *g1* → OFF
*g1* (Sims 10,11): if (Opa=OFF):
                      if (G1=ON) then *g1* → OFF; else *g1* → ON
               if (Opa=ON):
                      *g1* → OFF
*g2* (Sim 9): if (Opa=OFF):
                      if (G2=ON) then *g2* → ON; else *g2* → OFF
               if (Opa=ON):
                      *g2* → OFF
*g2* (Sims 10,11): if (Opa=OFF):
                      if (G2=ON) then *g2* → OFF; else *g2* → ON
               if (Opa=ON):
                      *g2* → OFF
~~~

Time delays.

**Sims 9-11**
|  |  |
| --- | --- |
| **Signal X decay delay** | 24 |
| **Opa synthesis delay** | 36 |
| **G1/G2 synthesis delay** | 18 |
| **G1/G2 decay delay** | 30 |
| **All other delays** | 6 |

###### Initial conditions

Simulation 9.

*Initial conditions:* Simulation 9:
|  | **C1** | **C2** | **C3** | **C4** | **C5** | **C6** | **C7** | **C8** |
| --- | --- | --- | --- | --- | --- | --- | --- | --- |
| **Signal X value** | 1 | 1 | 1 | 1 | 1 | 1 | 1 | 1 |
| **G1 value** | 0 | 0 | 0 | 1 | 1 | 1 | 1 | 1 |
| **G2 value** | 1 | 1 | 0 | 0 | 0 | 1 | 1 | 1 |

Simulations 10 and 11.

Simulations 10 and 11:
|  | **C1** | **C2** | **C3** | **C4** | **C5** | **C6** | **C7** | **C8** |
| --- | --- | --- | --- | --- | --- | --- | --- | --- |
| **Signal X value** | 1 | 1 | 1 | 1 | 1 | 1 | 1 | 1 |
| **$g1$ value** | 1 | 1 | 1 | 0 | 0 | 0 | 0 | 0 |
| **$g1$ age** | 0 | 6 | 12 | 0 | 0 | 0 | 0 | 0 |
| **G1 value** | 0 | 0 | 0 | 1 | 1 | 1 | 1 | 1 |
| **G1 age** | 0 | 0 | 0 | 0 | 6 | 12 | 18 | 24 |
| **$g2$ value** | 0 | 0 | 1 | 1 | 1 | 0 | 0 | 0 |
| **$g2$ age** | 0 | 0 | 0 | 6 | 12 | 0 | 0 | 0 |
| **G2 value** | 1 | 1 | 0 | 0 | 0 | 1 | 1 | 1 |
| **G2 age** | 18 | 24 | 0 | 0 | 0 | 0 | 6 | 12 |

##### Modified network simulations (Movies 12-13)

These two simulations alter the control logic of *hairy* and *eve* and remove the gap inputs from the network. In addition, the control logic of *opa* is altered so as to be regulated by Signal X. The time delays associated with Hairy, Signal X, and Opa are also adjusted.

Under these modified conditions, the only initial conditions that need be specified are those of Hairy and Signal X. They are assigned very different starting patterns in the two simulations, however the two simulations eventually generate the same final output pattern.

###### Control logic

~~~
*hairy*  if (Opa=OFF):
              if (Hairy=ON) then *hairy* → OFF; else *hairy* → ON
       if (Opa=ON):
               *hairy* → OFF
*eve*:   if (Opa=OFF):
              if (Runt=ON or Odd=ON) then *eve* → OFF; else *eve* → ON
       if (Opa=ON):
              if (Runt=ON or Odd=ON or En=ON or (Slp=ON and Eve=OFF)) then *eve* → OFF; *else* eve → ON
*opa:*   if (Signal X=ON) then *opa* → OFF; else *opa* → ON
~~~

Time delays:

|  | **synthesis** | **decay** |
| --- | --- | --- |
| **Hairy** | 30 | 18 |
| **Signal X** | 6 | 42 / 144 |
| **Opa** | 18 | 6 |

Note that the decay delay for signal X is greatly increased for Simulation 13.

###### Initial conditions

Simulation 12.

*Initial conditions:* Simulation 12:
|  | C1 | C2 | C3 | C4 | C5 | C6 | C7 | C8 |
| --- | --- | --- | --- | --- | --- | --- | --- | --- |
| **Signal X** | 1 | 1 | 1 | 1 | 1 | 1 | 1 | 1 |
| *hairy value* | 1 | 1 | 1 | 1 | 1 | 0 | 0 | 0 |
| *hairy age* | 0 | 6 | 12 | 18 | 24 | 0 | 0 | 0 |
| **Hairy value** | 0 | 0 | 0 | 0 | 0 | 1 | 1 | 1 |
| **Hairy age** | 0 | 0 | 0 | 0 | 0 | 0 | 6 | 12 |

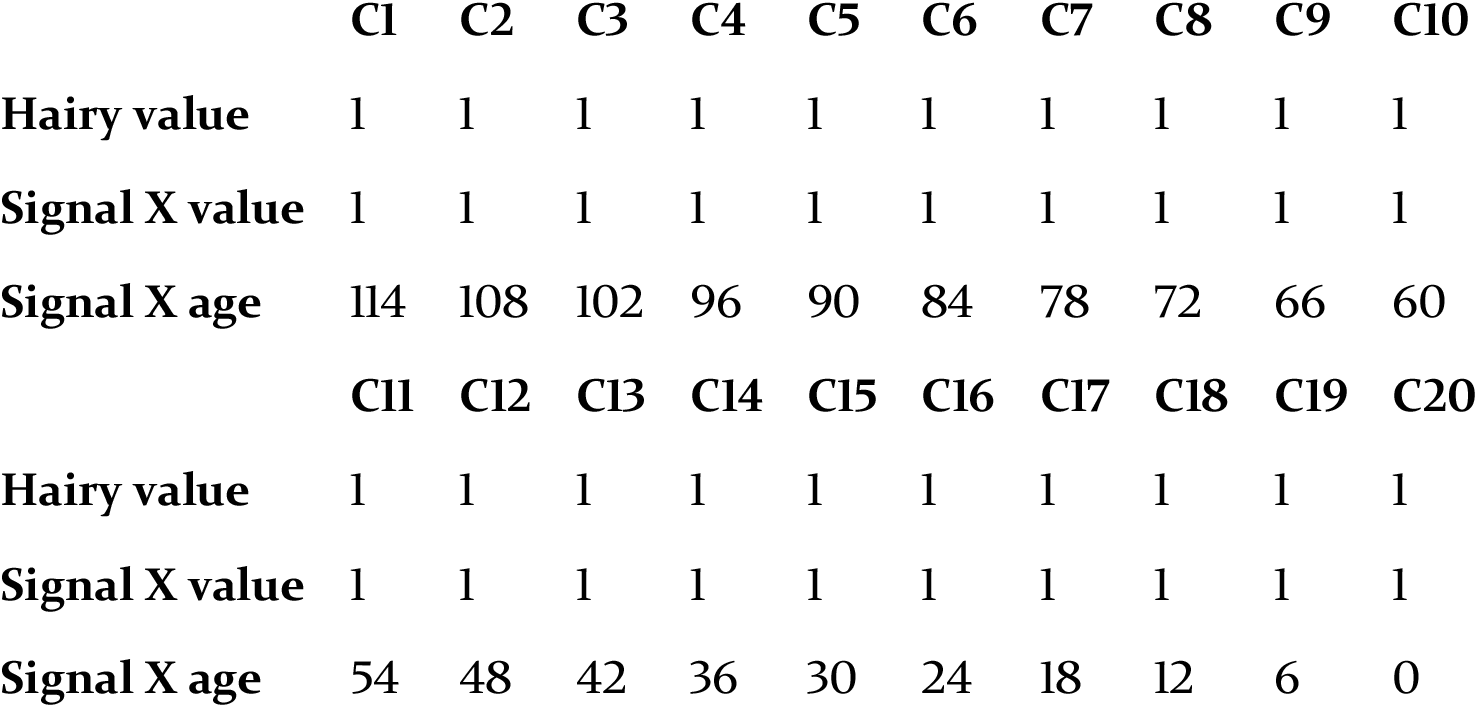
Simulation 13.

Simulation 13:
|  | C1 | C2 | C3 | C4 | C5 | C6 | C7 | C8 | C9 | C10 |
| --- | --- | --- | --- | --- | --- | --- | --- | --- | --- | --- |
| **Hairy value** | 1 | 1 | 1 | 1 | 1 | 1 | 1 | 1 | 1 | 1 |
| **Signal X value** | 1 | 1 | 1 | 1 | 1 | 1 | 1 | 1 | 1 | 1 |
| **Signal X age** | 114 | 108 | 102 | 96 | 90 | 84 | 78 | 72 | 66 | 60 |
|  | C11 | C12 | C13 | C14 | C15 | C16 | C17 | C18 | C19 | C20 |
| **Hairy value** | 1 | 1 | 1 | 1 | 1 | 1 | 1 | 1 | 1 | 1 |
| **Signal X value** | 1 | 1 | 1 | 1 | 1 | 1 | 1 | 1 | 1 | 1 |
| **Signal X age** | 54 | 48 | 42 | 36 | 30 | 24 | 18 | 12 | 6 | 0 |

## REFERENCES

1. Nüsslein-Volhard C, Wieschaus E. Mutations affecting segment number and polarity in Drosophila. Nature. 1980;287(5785):795–801.

2. Wieschaus E, Nüsslein-Volhard C. The Heidelberg Screen for Pattern Mutants of *Drosophila*: A Personal Account. Annu Rev Cell Dev Biol. 2016;32(1):annurev-cellbio-113015-023138.

3. Akam M. The molecular basis for metameric pattern in the Drosophila embryo. Development. 1987;101(1)1–22.

4. Nasiadka A, Dietrich BH, Krause HM. Anterior – posterior patterning in the Drosophila embryo. Adv Dev Biol Biochem. 2002;12:155–204.

5. Arnosti DN. ANALYSIS AND FUNCTION OF TRANSCRIPTIONAL REGULATORY ELEMENTS: Insights from Drosophila. Annu Rev Entomol. 2003;48(1):579–602.

6. Clark E, Akam M. Odd-paired controls frequency doubling in Drosophila segmentation by altering the pair-rule gene regulatory network. Elife. 2016;5:1–22.

7. Frohnhöfer HG, Nüsslein-Volhard C. Organization of anterior pattern in the Drosophila embryo by the maternal gene bicoid. Nature. 1986;324(6093)120–5.

8. Driever W, Nüsslein-Volhard C. A gradient of bicoid protein in Drosophila embryos. Cell. 1988;54(1):83–93.

9. Driever W, Nüsslein-Volhard C. The bicoid protein determines position in the Drosophila embryo in a concentration-dependent manner. Cell. 1988;54(1):95–104.

10. Tautz D, Lehmann R, Schnurch H, Schuh R, Seifert E, Kienlin A, et al. Finger protein of novel structure encoded by hunchback, a second member of the gap class of Drosophila segmentation genes. Nature. 1987;327(6121)383–9.

11. Rosenberg UB, Schroder C, Preiss A, Kienlin A, Cote S, Riede I, et al. Structural homology of the product of the Drosophila Kruppel gene with Xenopus transcription factor IIIA. Nature. 1986;319(6051)336–9.

12. Nauber U, Pankratz MJ, Kienlin A, Seifert E, Klemm U, Jäckle H. Abdominal segmentation of the Drosophila embryo requires a hormone receptor-like protein encoded by the gap gene knirps. Nature. 1988;336(6198)489–92.

13. Capovilla M, Eldon ED, Pirrotta V. The giant gene of Drosophila encodes a b-ZIP DNA-binding protein that regulates the expression of other segmentation gap genes. Development. 1992;114(1):99–112.

14. Jaeger J. The gap gene network. Cell Mol Life Sci. 2011;68(2):243–74.

15. Ingham P, Pinchin S. Genetic analysis of the hairy locus in Drosophila melanogaster. Genetics. 1985;463–86.

16. Macdonald PM, Ingham P, Struhl G. Isolation, structure, and expression of even-skipped: a second pair-rule gene of Drosophila containing a homeo box. Cell. 1986;47(5)721–34.

17. Gergen JP, Butler B a. Isolation of the Drosophila segmentation gene runt and analysis of its expression during embryogenesis. Genes Dev. 1988;2(9):1179–93.

18. Hafen E, Kuroiwa A, Gehring WJ. Spatial distribution of transcripts from the segmentation gene fushi tarazu during Drosophila embryonic development. Cell. 1984;37(3):833–41.

19. Coulter DE, Swaykus EA, Beran-Koehn MA, Goldberg D, Wieschaus E, Schedl P. Molecular analysis of odd-skipped, a zinc finger encoding segmentation gene with a novel pair-rule expression pattern. Embo J. 1990;8(12):3795–804.

20. Kilchherr F, Baumgartner S, Bopp D, Frei E, Noll M. Isolation of the paired gene of Drosophila and its spatial expression during early embryogenesis. 1986;321:493–9.

21. Grossniklaus U, Pearson RK, Gehring WJ. The Drosophila Sloppy Paired Locus Encodes Two Proteins Involved in Segmentation That Show Homology To Mammalian Transcription Factors. Genes Dev. 1992;6(6):1030–51.

22. Howard K, Ingham P, Rushlow C. Region-specific alleles of the Drosophila segmentation gene hairy. Genes Dev. 1988;2(8):1037–46.

23. Goto T, Macdonald P, Maniatis T. Early and late periodic patterns of even skipped expression are controlled by distinct regulatory elements that respond to different spatial cues. Cell. 1989;57(3):413–22.

24. Harding K, Hoey T, Warrior R, Levine M. Autoregulatory and gap gene response elements of the even-skipped promoter of Drosophila. EMBO J. 1989;8(4):1205–12.

25. Pankratz MJ, Jäckle H. Making stripes in the Drosophila embryo. Trends Genet. 1990;6(9):287–92.

26. Schroeder MD, Greer C, Gaul U. How to make stripes: deciphering the transition from non-periodic to periodic patterns in Drosophila segmentation. Development. 2011;138(14):3067–78.

27. Hiromi Y, Kuroiwa A, Gehring WJ. Control elements of the Drosophila segmentation gene fushi tarazu. Cell. 1985;43:603–13.

28. Dearolf CR, Topol J, Parker CS. Transcriptional control of Drosophila fushi tarazu zebra stripe expression. Genes Dev. 1989;3(3)384–98.

29. Butler B a, Soong J, Gergen JP. The Drosophila segmentation gene runt has an extended cis-regulatory region that is required for vital expression at other stages of development. Mech Dev. 1992;39(1–2):17–28.

30. Howard K, Ingham P. Regulatory interactions between the segmentation genes fushi tarazu, hairy, and engrailed in the Drosophila blastoderm. Cell. 1986;44(6):949–57.

31. Carroll SB, Scott MP. Zygotically active genes that affect the spatial expression of the fushi tarazu segmentation gene during early Drosophila embryogenesis. Cell. 1986;45(1):113–26.

32. Frasch M, Levine M. Complementary patterns of even-skipped and fushi tarazu expression involve their differential regulation by a common set of segmentation genes in Drosophila. 1987;0:981–95.

33. Ingham P, Gergen P. Interactions between the pair-rule genes runt, hairy, even-skipped and fushi tarazu and the establishment of periodic pattern in the Drosophila embryo. Development. 1988;104(Supplement):51–60.

34. Vavra SH, Carroll SB. The zygotic control of Drosophila pair-rule gene expression. II. Spatial repression by gap and pair-rule gene products. Development. 1989;107(3):663–72.

35. DiNardo S, O’Farrell PH. Establishment and refinement of segmental pattern in the Drosophila embryo: spatial control of engrailed expression by pair-rule genes. Genes Dev. 1987;1(10):1212–25.

36. Jaynes JB, Fujioka M. Drawing lines in the sand: even skipped et al. and parasegment boundaries. Dev Biol. 2004;269(2)609–22.

37. Martinez-Arias A, Lawrence PA. Parasegments and compartments in the Drosophila embryo. Nature. 1985;313(6004):639–42.

38. Ingham PW. The molecular genetics of embryonic pattern formation in Drosophila. Nature. 1988;335(6185):25–34.

39. DiNardo S, Heemskerk J, Dougan S, O’Farrell PH. The making of a maggot: patterning the Drosophila embryonic epidermis. Curr Opin Genet Dev. 1994;4(4):529–34.

40. Perrimon N. The genetic basis of patterned baldness in Drosophila. Cell. 1994;76(5)781–4.

41. Sanson B. Generating patterns from fields of cells: Examples from Drosophila segmentation. EMBO Rep. 2001;2(12):1083–8.

42. Grimm O, Coppey M, Wieschaus E. Modelling the Bicoid gradient. Development. 2010;137(14):2253–64.

43. Jaeger J, Blagov M, Kosman D, Kozlov KN, Manu, Myasnikova E, et al. Dynamical analysis of regulatory interactions in the gap gene system of Drosophila melanogaster. Genetics. 2004;167(4):1721–37.

44. Manu, Surkova S, Spirov A V., Gursky V V., Janssens H, Kim A-R, et al. Canalization of Gene Expression and Domain Shifts in the Drosophila Blastoderm by Dynamical Attractors. Shvartsman S, editor. PLoS Comput Biol. 2009 Mar 13;5(3):e1000303.

45. Gursky V V, Panok L, Myasnikova EM, Manu, Samsonova MG, Reinitz J,et al. Mechanisms of gap gene expression canalization in the Drosophila blastoderm. BMC Syst Biol. 2011;5(1):118.

46. Verd B, Clark E, Wotton KR, Janssens H, Jiménez-Guri E, Crombach A,et al. A damped oscillator imposes temporal order on posterior gap gene expression in Drosophila. bioRxiv. 2016;1–25.

47. Dubuis JO, Tkacik G, Wieschaus EF, Gregor T, Bialek W. Positional information, in bits. Proc Natl Acad Sci. 2013;110(41):16301–8.

48. Tkačik G, Dubuis JO, Petkova MD, Gregor T. Positional information, Positional error, and readout precision in morphogenesis: A mathematical framework. Genetics. 2015;199(1)39–59.

49. Petkova MD, Tkačik G, Bialek W, Wieschaus EF, Gregor T. Optimal decoding of information from a genetic network. 2016;

50. Small S, Blair A, Levine M. Regulation of even-skipped stripe 2 in the Drosophila embryo. EMBO J. 1992;11(11):4047–57.

51. Small S, Blair A, Levine M. Regulation of two pair-rule stripes by a single enhancer in the Drosophila embryo. Dev Biol. 1996;175(2):314–24.

52. Arnosti DN, Barolo S, Levine M, Small S. The eve stripe 2 enhancer employs multiple modes of transcriptional synergy. Development. 1996;122(1):205–14.

53. Fujioka M, Emi-Sarker Y, Yusibova GL, Goto T, Jaynes JB. Analysis of an even-skipped rescue transgene reveals both composite and discrete neuronal and early blastoderm enhancers, and multi-stripe positioning by gap gene repressor gradients. Development. 1999;126(11):2527–38.

54. Andrioli LPM, Vasisht V, Theodosopoulou E, Oberstein A, Small S. Anterior repression of a Drosophila stripe enhancer requires three position-specific mechanisms. Development. 2002;129(21):4931–40.

55. Janssens H, Hou S, Jaeger J, Kim AR, Myasnikova E, Sharp D,et al. Quantitative and predictive model of transcriptional control of the Drosophila melanogaster even skipped gene. Nat Genet. 2006;38(10)1159–65.

56. Ilsley GR, Fisher J, Apweiler R, De Pace AH, Luscombe NM. Cellular resolution models for even skipped regulation in the entire Drosophila embryo. Elife. 2013;2:e00522.

57. Vincent BJ, Estrada J, DePace AH. The appeasement of Doug: a synthetic approach to enhancer biology. Integr Biol. 2016;8:475–84.

58. Baumgartner S, Noll M. Network of interactions among pair-rule genes regulating paired expression during primordial segmentation of Drosophila. Mech Dev. 1990;33(1):1–18.

59. Manoukian AS, Krause HM. Concentration-dependent activities of the even-skipped protein in Drosophila embryos. Genes Dev. 1992;6(9):1740–51.

60. Manoukian AS, Krause HM. Control of segmental asymetry in Drosophila embryos. Development. 1993;118:785–96.

61. Fujioka M, Jaynes JB, Goto T. Early even-skipped stripes act as morphogenetic gradients at the single cell level to establish engrailed expression. Development. 1995;121(12):4371–82.

62. Mullen JR, DiNardo S. Establishing parasegments in Drosophila embryos: roles of the odd-skipped and naked genes. Vol. 169, Developmental biology. 1995. p. 295–308.

63. Sánchez L, Thieffry D. Segmenting the fly embryo: a logical analysis of the pair-rule cross-regulatory module. J Theor Biol. 2003;224:517–37.

64. Sander K. Specification of the Basic Body Pattern in Insect Embryogenesis. In: Treherne JE, Berridge MJ, Wigglesworth VB, editors. Advances in Insect Physiology. Elsevier Science; 1976. (Advances in Insect Physiology).

65. Damen WGM, Janssen R, Prpic NM. Pair rule gene orthologs in spider segmentation. Evol Dev. 2005;7(6):618–28.

66. Choe CP, Miller SC, Brown SJ. A pair-rule gene circuit defines segments sequentially in the short-germ insect Tribolium castaneum. Proc Natl Acad Sci U S A. 2006;103(17) 6560–4.

67. Janssen R, Budd GE, Prpic N-M, Damen WG. Expression of myriapod pair rule gene orthologs. Evodevo. 2011;2:5.

68. Green J, Akam M. Evolution of the pair rule gene network: Insights from a centipede. Dev Biol. 2013;382(1):235–45.

69. Sarrazin AF, Peel AD, Averof M. A segmentation clock with two-segment periodicity in insects. Science (80-). 2012;336(6079):338–41.

70. El-Sherif E, Averof M, Brown SJ. A segmentation clock operating in blastoderm and germband stages of Tribolium development. Development. 2012;139(23):4341–6.

71. Cooke J, Zeeman EC. A clock and wavefront model for control of the number of repeated structures during animal morphogenesis. J Theor Biol. 1976;58(2):455–76.

72. Liu PZ, Kaufman TC. Short and long germ segmentation: Unanswered questions in the evolution of a developmental mode. Evol Dev. 2005;7(6):629–46.

73. Jaeger J, Surkova S, Blagov M, Janssens H, Kosman D, Kozlov KN,et al. Dynamic control of positional information in the early Drosophila embryo. Nature. 2004330(6997)368–71.

74. Surkova S, Kosman D, Kozlov K, Manu, Myasnikova E, Samsonova A a.,et al. Characterization of the Drosophila segment determination morphome. Dev Biol. 2008;313(2)844–62.

75. El-Sherif E, Levine M. Shadow Enhancers Mediate Dynamic Shifts of Gap Gene Expression in the Drosophila Embryo. Curr Biol. 2016;1–6.

76. Jaeger J, Reinitz J. On the dynamic nature of positional information. BioEssays. 2006;28(11):1102–11.

77. Liu F, Morrison AH, Gregor T. Dynamic interpretation of maternal inputs by the Drosophila segmentation gene network. Proc Natl Acad Sci U S A. 2013;110(17):6724–9.

78. Perkins TJ, Jaeger J, Reinitz J, Glass L. Reverse engineering the gap gene network of Drosophila melanogaster. PLoS Comput Biol. 2006;2(5)417–28.

79. Keränen SVE, Fowlkes CC, Luengo Hendriks CL, Sudar D, Knowles DW, Malik J,et al. Three-dimensional morphology and gene expression in the Drosophila blastoderm at cellular resolution II: dynamics. Genome Biol. 2006;7(12)R124.

80. Benedyk MJ, Mullen JR, DiNardo S. Odd-paired: A zinc finger pair-rule protein required for the timely activation of engrailed and wingless in Drosophila embryos. Genes Dev. 1994;8(1):105–17.

81. Yu Y, Pick L. Non-periodic cues generate seven ftz stripes in the Drosophila embryo. Mech Dev. 1995;50(2–3):163–75.

82. Klingler M, Soong J, Butler B, Gergen JP. Disperse versus compact elements for the regulation of runt stripes in Drosophila. Dev Biol. 1996;177(1) 73–84.

83. Edgar BA, Weir MP, Schubiger G, Kornberg T. Repression and turnover pattern fushi tarazu RNA in the early Drosophila embryo. Cell. 1986;47(5)747–54.

84. Weir MP, Edgar B a, Kornberg T, Schubiger G. Spatial regulation of engrailed expression in the Drosophila embryo. Genes Deve. 1988;2:1194–203.

85. Vavra SH, Carroll SB. The zygotic control of Drosophila pair-rule gene expression. I. A search for new pair-rule regulatory loci. Development. 1989;107(3):663–72.

86. Nasiadka A, Krause HM. Kinetic analysis of segmentation gene interactions in Drosophila embryos. Development. 1999;126(7):1515–26.

87. Hiromi Y, Gehring WJ. Regulation and Function of the Drosophila Segmentation Gene Fushi-Tarazu. Cell. 1987;50(6):963–74.

88. Schier A, Gehring W. Direct homeodomain–DNA interaction in the autoregulation of the fushi tarazu gene. Nature. 1992;356(6372):804–7.

89. Ish-Horowicz D, Pinchin SM. Pattern abnormalities induced by ectopic expression of the Drosophila gene hairy are associated with repression of ftz transcription. Cell. 1987;51(3):405–15.

90. Kok K, Ay A, Li LM, Arnosti DN. Genome-wide errant targeting by Hairy. Elife. 2015;4(AUGUST2015):1–21.

91. Arnosti DN, Gray S, Barolo S, Zhou J, Levine M. The gap protein knirps mediates both quenching and direct repression in the Drosophila embryo. EMBO J. 1996;15(14):3659–66.

92. Barolo S, Levine M. hairy mediates dominant repression in the Drosophila embryo. EMBO J. 1997;16(10):2883–91.

93. Li LM, Arnosti DN. Long- and short-range transcriptional repressors induce distinct chromatin states on repressed genes. Curr Biol. 2011;21(5)406–12.

94. Meinhardt H. Models of biological pattern formation. Acad Press London. 1982;1–211.

95. Meinhardt H. Models for positional signalling, the threefold subdivision of segments and the pigmentation pattern of molluscs. J Embryol Exp Morphol. 1984;83 Suppl:289–311.

96. Meinhardt H. Hierarchical inductions of cell states: a model for segmentation in Drosophila. J Cell Sci Suppl. 1986;4:357–81.

97. DiNardo S, Sher E, Heemskerk-Jongens J, Kassis JA, O’Farrell PH. Two-tiered regulation of spatially patterned engrailed gene expression during Drosophila embryogenesis. Nature. 1988;332(6165):604–9.

98. Gergen JP, Wieschaus E. Dosage requirements for runt in the segmentation of Drosophila embryos. Cell. 1986;45(2):289–99.

99. Lardelli M, Ish-Horowicz D. Drosophila hairy pair-rule gene regulates embryonic patterning outside its apparent stripe domains. Development. 1993;118(1):255–66.

100. Fujioka M, Yusibova GL, Patel NH, Brown SJ, Jaynes JB. The repressor activity of Even-skipped is highly conserved, and is sufficient to activate engrailed and to regulate both the spacing and stability of parasegment boundaries. Development. 2002;129(19):4411–21.

101. Little SC, Tikhonov M, Gregor T. Precise developmental gene expression arises from globally stochastic transcriptional activity. Cell. 2013;154(4):789–800.

102. Desponds J, Tran H, Ferraro T, Lucas T, Perez Romero C, Guillou A,et al. Precision of Readout at the hunchback Gene: Analyzing Short Transcription Time Traces in Living Fly Embryos. PLOS Comput Biol. 2016;12(12):e1005256.

103. Edgar B a, Odell GM, Schubiger G. A genetic switch, based on negative regulation, sharpens stripes in Drosophila embryos. Dev Genet. 1989;10(3):124–42.

104. Edgar BA, O’Dell GM, Schubiger G. Cytoarchitecture and the patterning of fushi-tarazu expression in the Drosophila blastoderm. Genes Dev. 1987;1:1226–37.

105. Davis I, Ish-Horowicz D. Apical localization of pair-rule transcripts requires 3’ sequences and limits protein diffusion in the Drosophila blastoderm embryo. Cell. 1991;67(5):927–40.

106. Klingler M, Gergen JP. Regulation of runt transcription by Drosophila segmentation genes. Mech Dev. 1993;43(1):3–19.

107. Copeland JW, Nasiadka A, Dietrich BH, Krause HM. Patterning of the Drosophila embryo by a homeodomain-deleted Ftz polypeptide. Vol. 379, Nature. 1996. p. 162–5.

108. Bownes M. A photographic study of development in the living embryo of Drosophila melanogaster. J Embryol Exp Morphol. 1975;33(3):789–801.

109. Campos-Ortega JA, Hartenstein V. The Embryonic Development of Drosophila melanogaster. 1985.

110. Jürgens G, Wieschaus E, Nüsslein-Volhard C, Kluding H. Mutations affecting the pattern of the larval cuticle in Drosophila melanogaster II. Zygotic loci on the third chromosome. Wilhelm Roux’s Arch Dev Biol. 1984;193:283–95.

111. Fowlkes CC, Luengo Hendriks CL, Ker??nen SVE, Weber GH, R??bel O, Huang MY,et al. A Quantitative Spatiotemporal Atlas of Gene Expression in the Drosophila Blastoderm. Cell. 2008;133(2):364–74.

112. Nusslein-Volhard C, Kluding H, Jurgens G. Genes affecting the segmental subdivision of the Drosophila embryo. Cold Spring Harb Symp Quant Biol. 1985;50:145–54.

113. Harding K, Rushlow C, Doyle H, Hoey T, Levine M. Cross-regulatory interactions among pair-rule genes in Drosophila. Science (80-). 1986;233(4767):953–9.

114. Riechmann V, Irion U, Wilson R, Grosskortenhaus R, Leptin M. Control of cell fates and segmentation in the Drosophila mesoderm. Development. 1997;124(15):2915–22.

115. Coulter DE, Wieschaus E. Gene activities and segmental patterning in Drosophila: analysis of odd-skipped and pair-rule double mutants. Genes Dev. 1988;2(12B):1812–23.

116. True JR, Haag ES. Developmental system drift and flexibility in evolutionary trajectories. Evol Dev. 2001;3(2):109–19.

117. Lewis J. Autoinhibition with Transcriptional Delay. Curr Biol. 2003 Aug;13(16):1398–408.

118. Schröter C, Ares S, Morelli LG, Isakova A, Hens K, Soroldoni D,et al. Topology and dynamics of the zebrafish segmentation clock core circuit. PLoS Biol. 2012;10(7):11.

119. Bertrand E, Chartrand P, Schaefer M, Shenoy SM, Singer RH, Long RM. Localization of ASH1 mRNA Particles in Living Yeast. Mol Cell. 1998;2(4):437–45.

120. Forrest KM, Gavis ER. Live Imaging of Endogenous RNA Reveals a Diffusion and Entrapment Mechanism for nanos mRNA Localization in Drosophila. Curr Biol. 2003 Jul;13(14):1159–68.

121. Garcia HG, Tikhonov M, Lin A, Gregor T. Quantitative Imaging of Transcription in Living Drosophila Embryos Links Polymerase Activity to Patterning. Curr Biol. 2013;23(21):2140–5.

122. Andrioli LP, Oberstein AL, Corado MSG, Yu D, Small S. Groucho-dependent repression by Sloppy-paired 1 differentially positions anterior pair-rule stripes in the Drosophila embryo. Dev Biol. 2004;276(2):541–51.

123. Bothma JP, Garcia HG, Esposito E, Schlissel G, Gregor T, Levine M. Dynamic regulation of eve stripe 2 expression reveals transcriptional bursts in living Drosophila embryos. Proc Natl Acad Sci. 2014;111(29):10598–603.

124. Paré AC, Vichas A, Fincher CT, Mirman Z, Farrell DL, Mainieri A, et al. A positional Toll receptor code directs convergent extension in Drosophila. Nature. 2014;515(7528):523–7.

125. Benton MA, Pechmann M, Frey N, Stappert D, Conrads KH, Chen Y-T, et al. Toll Genes Have an Ancestral Role in Axis Elongation. Curr Biol. 2016;26.

126. Pueyo JI, Lanfear R, Couso JP. Ancestral Notch-mediated segmentation revealed in the cockroach Periplaneta americana. Proc Natl Acad Sci U S A. 2008;105(43):16614–9.

127. El-Sherif E, Zhu X, Fu J, Brown SJ. Caudal Regulates the Spatiotemporal Dynamics of Pair-Rule Waves in Tribolium. PLoS Genet. 2014;10(10).

128. Peel A, Akam M. Evolution of segmentation: Rolling back the clock. Curr Biol. 2003;13(18):708–10.

129. Rosenberg MI, Brent AE, Payre F, Desplan C. Dual mode of embryonic development is highlighted by expression and function of Nasonia pair-rule genes. Elife. 2014;2014(3):1–24.

130. Nakao H. Analyses of interactions among pair-rule genes and the gap gene Krüppel in Bombyx segmentation. Dev Biol. 2015;405(1):1–9.

131. Nakao H. Hunchback knockdown induces supernumerary segment formation in Bombyx. Dev Biol. 2016;413(2):207–16.

132. Rothschild JB, Tsimiklis P, Siggia ED, François P. Predicting Ancestral Segmentation Phenotypes from Drosophila to Anopheles Using In Silico Evolution. PLOS Genet. 2016;12(5):e1006052.

133. Wotton KR, Jimenez-Guri E, Crombach A, Janssens H, Alcaine-Colet A, Lemke S, et al. Quantitative system drift compensates for altered maternal inputs to the gap gene network of the scuttle fly megaselia abdita. Elife. 2015;2015(4):1–28.

134. Crombach A, Wotton KR, Jiménez-Guri E, Jaeger J. Gap Gene Regulatory Dynamics Evolve along a Genotype Network. Mol Biol Evol. 2016;33(5):1293–307.

135. Clark E, Akam M. Data from: Odd-paired controls frequency doubling in Drosophila segmentation by altering the pair-rule gene regulatory network [Internet]. Dryad Digital Repository. Dryad Digital Repository; 2016.

136. Schindelin J, Arganda-Carreras I, Frise E, Kaynig V, Longair M, Pietzsch T, et al. Fiji: an open-source platform for biological-image analysis. Nat Methods. 2012;9(7):676–82.

137. van der Walt S, Colbert SC, Varoquaux G. The NumPy Array: A Structure for Efficient Numerical Computation. Comput Sci Eng. 2011 Mar;13(2):22–30.

138. Hunter JD. Matplotlib: A 2D Graphics Environment. Comput Sci Eng. 2007 May;9(3):90–5.

139. Tracey WD, Ning X, Klingler M, Kramer SG, Gergen JP. Quantitative analysis of gene function in the Drosophila embryo. Genetics. 2000;154(i):273–84.

140. Cadigan KM, Grossniklaus U, Gehring WJ. Localized Expression of Sloppy Paired Protein Maintains the Polarity of Drosophila Parasegments. Genes Dev. 1994;8(8):899–913.

141. Farzana L, Brown SJ. Hedgehog signaling pathway function conserved in Tribolium segmentation. 2008;10:181–92.

142. Janssen R, Budd GE. Deciphering the onychophoran “segmentation gene cascade”: Gene expression reveals limited involvement of pair rule gene orthologs in segmentation, but a highly conserved segment polarity gene network. Dev Biol. 2013;382(1):224–34.

143. Peter IS, Faure E, Davidson EH. Predictive computation of genomic logic processing functions in embryonic development. Proc Natl Acad Sci. 2012;109(41):16434–42.

144. Pisarev A, Poustelnikova E, Samsonova M, Reinitz J. FlyEx, the quantitative atlas on segmentation gene expression at cellular resolution. Nucleic Acids Res. 2009;37(SUPPL. 1)560–6.

145. Saulier-Le Dréan B, Nasiadka a, Dong J, Krause HM. Dynamic changes in the functions of Odd-skipped during early Drosophila embryogenesis. Development. 1998;125(23):4851–61.

146. Hooper KL, Parkhurst SM, Ish-Horowicz D. Spatial control of hairy protein expression during embryogenesis. Development. 1989;107(3):489–504.

147. Riddihough G, Ish-Horowicz D. Individual stripe regulatory elements in the Drosophila hairy promoter respond to maternal, gap, and pair-rule genes. Genes Dev. 1991;5(5):840–54.

148. Hartmann C, Taubert H, Jäckle H. A two-step mode of stripe formation in the Drosophila blastoderm requires interactions among primary pair rule genes. Mech o. 1994;45(l):3–13.

149. Jiménez G, Pinchin SM, Ish-Horowicz D. In vivo interactions of the Drosophila Hairy and Runt transcriptional repressors with target promoters. EMBO J. 1996;15(24)7088–98.

150. Tsai C, Gergen JP. Gap gene properties of the pair-rule gene runt during Drosophila segmentation. Development. 1994;120(6):1671–83.

151. Chen H, Xu Z, Mei C, Yu D, Small S. A System of Repressor Gradients Spatially Organizes the Boundaries of Bicoid-Dependent Target Genes. Cell. 2012;149(3)618–29.

152. Carroll SB, Laughon a, Thalley BS. Expression, function, and regulation of the hairy segmentation protein in the Drosophila embryo. Genes Dev. 1988;2(7):883–90.

153. Nasiadka A, Grill A, Krause HM. Mechanisms regulating target gene selection by the homeodomain-containing protein Fushi tarazu. Development. 2000;127(13):2965–76.

154. Goldstein RE, Cook O, Dinur T, Pisante A, Karandikar UC, Virginia W. An eh1-Like Motif in Odd-skipped Mediates Recruitment of Groucho and Repression In Vivo. Mol Cell Biol. 2005;25(24):10711–20.

155. Warrior R, Levine M. Dose-dependent regulation of pair-rule stripes by gap proteins and the initiation of segment polarity. Development. 1990;110(3)759–67.

156. Kania MA, Bonner AS, Duffy JB, Kania MA, Bonner AS, Duffy JB, et al. The Drosophila segmentation gene runt encodes a novel nuclear regulatory protein that is also expressed in the developing nervous system. Genes Dev. 1990;1701–13.

157. Han K, Manley JL. Transcriptional repression by the Drosophila even-skipped protein: Definition of a minimal repression domain. Genes Dev. 1993;7(3):491–503.

158. Austin RJ, Biggin MD. A domain of the even-skipped protein represses transcription by preventing TFIID binding to a promoter: repression by cooperative blocking. Mol Cell Biol. 1995;15(9):4683–93.

159. Um M, Li C, Manley JL. The transcriptional repressor even-skipped interacts directly with TATA-binding protein. Mol Cell Biol. 1995;15(9):5007–16.

160. Lawrence PA, Johnston P. Pattern formation in the Drosophila embryo: allocation of cells to parasegments by even-skipped and fushi tarazu. Development. 1989;105(4):761–7.

161. Kauffman SA. Homeostasis and Differentiation in Random Genetic Control Networks. Nature. 1969;224(5215):177–8.

162. Thomas R. Boolean formalization of genetic control circuits. J Theor Biol. 1973;42(3):563–85.

163. Jong H De. Modeling and Simulation of Genetic Regulatory Systems: A Literature Review Modeling and Simulation of Genetic Regulatory Systems: A Literature Review. 2006;

164. Mbodj A, Junion G, Brun C, Furlong EEM, Thieffry D. Logical modelling of Drosophila signalling pathways. Mol Biosyst. 2013;9(9):2248–58.

165. Thomas R. Regulatory Networks Seen as Asynchronous Automata: A Logical Description. J Theor Biol. 1991;1–23.

